# Is epigenetics Lamarck’s revenge? The evolutionary and ecological impacts of DNA methylation in a California oak

**DOI:** 10.64898/2026.10.07.756922

**Authors:** Victoria L. Sork, Lily Peck, Luke Browne, Matteo Pellegrini

## Abstract

- Epigenetics has been proposed as a basis for Lamarckian inheritance of acquired characteristics. This hypothesis raises questions about environmental induction of DNA methylation, its persistence across generations, and associations with gene expression. We investigated these processes in the California oak, *Quercus lobata*.
- Based on 40 range-wide adult trees and their progeny grown in two common gardens, we tested four predictions: (i) DNA methylation is environmentally induced; (ii) methylation shows parent-offspring correspondence; (iii) environmentally induced patterns remain stable after transmission; and (iv) DNA methylation is associated with gene expression, varying by sequence context and TE proximity.
- Using methylomes and transcriptomes, we found support for all predictions except the third: CHH methylation was environmentally induced, but rarely showed parent-offspring correspondence, whereas CG methylation showed the opposite pattern. DNA methylation was associated with gene expression, depending on the sequence context and proximity to TEs.
- We identified one stress-associated gene with environmentally responsive CHH methylation that showed parent–offspring correspondence in both gardens. Still, our evidence for Lamarckian-like evolution is not compelling. Nevertheless, DNA methylation was environmentally responsive, showed parent–offspring correspondence, and was associated with gene expression, highlighting its importance for the ecological and evolutionary responses of long-lived trees to environmental change.

## Introduction

Is epigenetics “Lamarck’s revenge”, challenging the Darwinian concept of evolution (Ward, 2018) and providing a mechanism for the inheritance of environmentally induced phenotypic changes? Does epigenetics provide a molecular mechanism for Lamarck’s theory that the environment can induce heritable phenotypes? Skinner (2015) argues that environmental epigenetic processes can shape heritable phenotypic variation that can subsequently be acted upon by natural selection. With advances in 21^st^-century DNA sequencing, it is now possible to study genetic and epigenetic variation simultaneously in natural populations and evaluate Skinner’s (2015) “neo-Lamarckian” hypothesis. In contrast, Loison (2018) argues that epigenetic processes do not support the Lamarckian concept of the inheritance of acquired characteristics because epigenetic changes persist for only a few generations at best. Integrating genetic and epigenetic variation across environments can therefore help resolve this ongoing debate.

Epigenetic processes are likely to be especially important in plants, particularly sessile, long-lived species that experience substantial climatic variation over their lifespans (Bossdorf *et al*., 2008; Bräutigam *et al*., 2013; Duncan *et al*., 2014; Verhoeven *et al*., 2016; Richards *et al*., 2017; Rey *et al*., 2020). Epigenetic mutations may generate additional heritable phenotypic variation that can be acted upon by local selection pressures (Schmitz *et al*., 2011; Becker & Weigel, 2012; Schmitz, 2014). However, only environmentally induced epimutations satisfy a key criterion for Lamarckian inheritance, and distinguishing between spontaneous and induced epimutations remains difficult (Becker & Weigel, 2012). Environmentally induced epigenetic marks could lead to phenotypic plasticity that allows trees to respond to fluctuating climatic conditions (Romero-Mujalli *et al*., 2024; Miryeganeh & Armitage, 2025). Because these responses may be advantageous only under current environmental conditions, they need not be inherited. If environmentally induced epigenetic variation contributes to evolution, several testable predictions follow concerning the induction, inheritance, and functional consequences of DNA methylation. Thus, a Lamarckian perspective provides a useful framework for evaluating the evolutionary significance of environmentally induced epigenetic variation, regardless of whether epigenetic mechanisms ultimately require a revision of evolutionary theory (Skinner, 2015; Ward, 2018).

DNA methylation is one of the most studied epigenetic mechanisms in plants and can influence phenotypic variation through transposable element silencing and the regulation of gene expression (Law & Jacobsen, 2010; Schmitz *et al*., 2013a; Niederhuth & Schmitz, 2014; Verhoeven *et al*., 2016; Richards *et al*., 2017; Peck & Sork, 2024). Heritable DNA methylation has been widely documented in plants (Johannes *et al*., 2009; Paszkowski & Grossniklaus, 2011; Miryeganeh & Armitage, 2025), although its stability varies among methylation contexts (Schmitz *et al*., 2011). The evolutionary potential of environmentally responsive DNA methylation has evolutionary potential depends partly on sequence context because CG, CHG, and CHH methylation (where H is any non-guanine nucleotide) are established and maintained by different molecular pathways (Law & Jacobsen, 2010; Schmitz *et al*., 2013a). CG and CHG methylation occur at symmetric cytosine sites and can be maintained through mitosis and, in some cases, across generations. Consequently, phenotypic variation associated with these methylated states may be subject to natural selection until the methylation state is lost. In contrast, CHH methylation is asymmetric and must be continually re-established, making it more dynamic and less likely to be transmitted across generations. Most known epialleles, including those controlling floral phenotypes in *Linaria vulgaris* and *Arabidopsis thaliana* (Cubas *et al*., 1999; Cao & Jacobsen, 2002), are not environmentally induced. Notably, Song *et al*. (2025) found that hypomethylation of three CG and three CHG sites conferred acquired and heritable cold tolerance in rice **f**ollowing experimental cold treatment. This study represents one of the strongest examples to date of a pattern consistent with Lamarckian inheritance.

DNA methylation may provide a mechanism of phenotypic plasticity in response to environmental conditions regardless of its heritability (Bossdorf *et al*., 2010; Richards *et al*., 2010; Herman & Sultan, 2011; Zhang *et al*., 2013; Duncan *et al*., 2014; Schlichting & Wund, 2014; Sow, M. D. *et al*., 2018; Moler *et al*., 2019; Peck & Sork, 2024). Environmentally induced plasticity would be especially useful for tree populations because their long lifespans expose them to diverse and changing environments (Bräutigam *et al*., 2013; Sow, Mamadou Dia *et al*., 2018; Klupczynska & Ratajczak, 2021; Kurpisz & Pawlowski, 2022). Consistent with this potential for plasticity, numerous studies have documented methylation differences among populations or treatments that experience contrasting environments (Richards *et al*., 2012; Rico *et al*., 2014; Foust *et al*., 2016; Gugger *et al*., 2016; Keller *et al*., 2016; Galanti *et al*., 2022) and responses to environmental stress (e.g., Chinnusamy & Zhu, 2009; Kinoshita & Seki, 2014; Liang *et al*., 2014; Laanen *et al*., 2021). However, evidence that DNA methylation differs among environments does not, by itself, demonstrate environmental causation or that it contributes to phenotypic variation. Several studies provide such evidence. For example, in *Arabidopsis thaliana*, methylation changes in response to salicylic acid are associated with expression of stress-response genes (Dowen *et al*., 2012). In another Arabidopsis study, Zhang *et al*. (2013) show that DNA methylation can contribute to substantial heritable variation in ecologically important plant traits. In *Populus,* Zhou *et al*. (2023) report that geographically differentiated methylation patterns are associated with drought resistance and that drought induces methylation changes in promoter regions that alter the expression of drought-response genes. Together, these studies suggest that DNA methylation can contribute to environmentally induced phenotypic plasticity and, in some contexts, to heritable responses to environmental conditions.

A critical factor shaping the function of DNA methylation in generating phenotypic variation is the presence of transposable elements (TEs) (Klein & Anderson, 2022). TEs, which are repetitive DNA elements that can mobilize within genomes, can affect gene expression and thereby generate phenotypic variation that may be acted upon by natural selection and contribute to adaptation (Stapley *et al*., 2015; Schrader & Schmitz, 2019). Several studies have reported that TE insertions are associated with changes in phenotype (Studer *et al*., 2011; Ausin *et al*., 2016; Castanera *et al*., 2023). TEs could facilitate rapid adaptation, for example, through DNA methylation changes during colonization (Stapley *et al*., 2015). TEs can also be involved in stress responses. For example, in Arabidopsis, some TE families preferentially insert near stress-responsive genes and can suppress their expression (Ito *et al*., 2016). TEs can affect gene expression depending on whether they are inserted upstream or downstream of genes or within gene bodies, and their location may also influence whether CG, CHG, or CHH methylation is involved (Slotkin & Martienssen, 2007; Hollister & Gaut, 2009). Thus, to understand the functional role of DNA methylation in plant responses to the environment, it is necessary to consider TE position, DNA methylation sequence context, and gene expression together.

The overarching goal of this study is to evaluate the potential evolutionary and ecological consequences of DNA methylation in natural populations of the long-lived tree species, Q*uercus lobata.* We test three predictions central to a Lamarckian framework for the inheritance of acquired characteristics: (i) DNA methylation is environmentally induced; (ii) methylation is transmitted across generations; and (iii) environmentally induced patterns remain stable after transmission. We also test a fourth, functional prediction: DNA methylation is associated with gene expression, with these associations varying by sequence context and TE position.

Functional effects of methylation are relevant because they could generate the phenotypic variation underlying acquired characteristics. We test the first three predictions using reduced representation bisulphite sequencing (RRBS) data from 40 *Q. lobata* adult trees from natural populations and their 5-year-old progeny growing in two common gardens. We test the fourth prediction using whole-genome bisulphite sequencing (WGBS) and RNA-sequencing data of 42 7-year-old trees from these same families growing one of the common gardens. We find at least one case consistent with Lamarckian inheritance, but our broader results do not support Lamarckian inheritance as a widespread phenomenon in *Q. lobata*. Instead, our findings reveal important evolutionary, ecological, and functional consequences on DNA methylation, particularly through its interactions with transposable elements.

## Materials and Methods

### Study System

Valley oak, *Quercus lobata* Née (Fagaceae), is an endemic oak of California distributed along the foothills of the Coastal and Sierra-Nevada ranges (Pavlik et al., 1995). The species range includes a broad range of environmental conditions associated with latitudinal and longitudinal spread and elevational heterogeneity. This study system is supported by a high-quality, annotated reference genome, and methylome analyses (Sork *et al*., 2022), as well as prior evidence that DNA methylation is associated with environmental responses (Platt *et al*., 2015; Gugger *et al*., 2016; Browne *et al*., 2020; Browne *et al*., 2021). In fact, several studies have demonstrated correlations between environmental and genetic gradients (Sork *et al*., 2010; Gugger *et al*., 2013; Sork *et al*., 2016; Gugger *et al*., 2021). Moreover, under experimental drought conditions, seedlings from xeric and mesic parts of the species range show differences in gene expression, which may reflect underlying genetic or epigenetic variation (Gugger *et al*., 2017; Mead *et al*., 2019). Likewise, our prior studies of natural populations of *Q. lobata* indicate differences with leaf traits (Browne et al., 2021), which is suggestive that DNA methylation may be associated with phenotypic variation. Finally, Gugger *et al*. (2016) discovered several CG-differentially methylated positions that were correlated with climate gradients, in particular, several that were found in the gene bodies of DREB (Dehydration-responsive element-binding) genes, which are transcription factors shown to be related to drought tolerance (Hrmova & Hussain, 2021).

### Experimental design

To test the first three predictions, we selected 40 of 58 naturally occurring *Q. lobata* adult trees that were previously studied (Gugger *et al*., 2016) and had progeny growing in two common gardens (Figure 1). These gardens are part of a species-wide provenance study initiated in 2012 with progeny from 674 families and 95 provenances (i.e., sets of families from the same general locality) (Delfino Mix et al., 2015). The original planting design included five progeny per maternal seed source per garden, outplanted in 2014. We consider these progenies to be half-siblings (half-sibs) from the same maternal family, assuming that the open-pollinated offspring share the same mother but have multiple fathers. The two common gardens are located at the USDA Forest Service Chico Seed Orchard at Chico, California and the USDA Forest Service Institute of Forest Genetics, Placerville, California, respectively (hereafter, referred to as the Chico and Placerville gardens). The two sites differ environmentally in several respects. Chico, the warmer and dryer site (∼60 m above sea level), experiences longer growing seasons, higher summer maximum temperatures, and less precipitation than Placerville (∼600m above sea level), a slightly cooler and wetter site located outside the current species range of *Q. lobata*.

**Figure 1.**
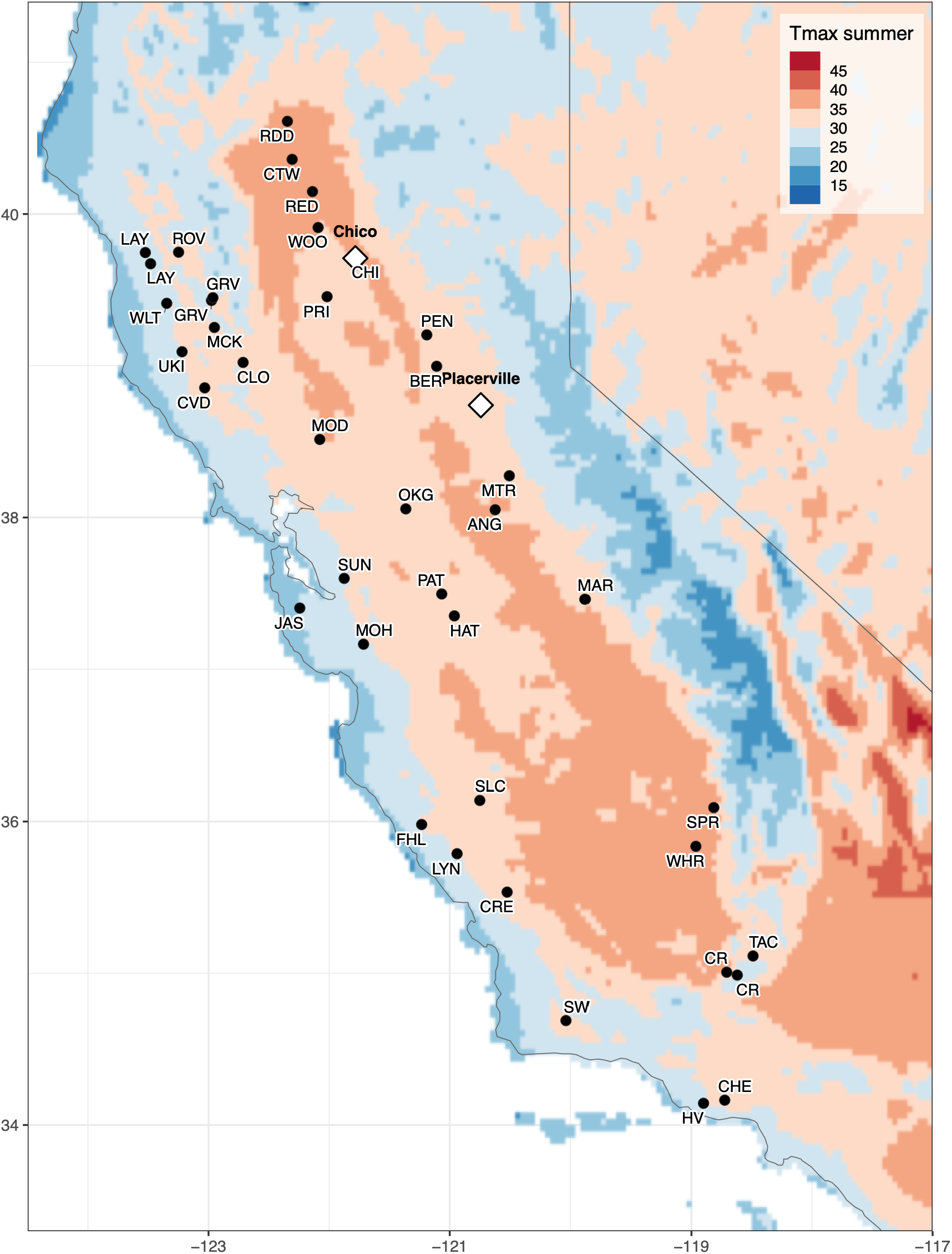
Locations of 40 valley oak (*Quercus lobata*) adult trees (IDs of localities indicated, see Table S1 for details on localities) used as seed sources for their half-sibling progeny grown in two common gardens. Gradients reflect the mean maximum temperature in summer.

For these tests, we collected leaf tissue from the 40 adults plus two five-year-old progeny per adult in both gardens from 40 families yielding 1200 total samples (Table S1). In addition, to assess methylation levels within individual trees between years, we selected 21 trees from16 families growing in the Chico garden and collected leaf tissue in Fall 2016 and Spring 2017. To test the fourth prediction concerning DNA methylation and gene expression, we selected 42 seven-year-old *Q. lobata* trees growing in the Placerville garden (Table S1) for which whole-genome bisulphite sequencing (WGBS) and bulk RNA-seq data were collected.

### Library preparation and sequencing

#### Reduced-Representation Bisulphite Sequencing (RRBS)

We followed methods established in the Pellegrini Lab as summarized by Fu et al. (2018) and described elsewhere (Gugger *et al*., 2016) for the adult seed source of the progeny studied here. Briefly, we extracted total genomic DNA from the leaves using a prewash protocol (Li *et al*., 2007) followed by Qiagen DNeasy Plant extraction kits. Total genomic DNA was digested with 20m of MspI (CCGG). Samples were pooled and purified using an equal volume of SPRI beads (Beckman Coulter, cat # B23318). Size selection was performed using SPRI beads to enrich for fragments from 200 to 400 bp. Bisulphite treatment was performed using Epitect Bisulphite kit (QIAGEN, cat # 59104) according to manufacturer’s protocol, except that two consecutive rounds of conversion are performed, for a total of 10 hours of incubation. Purified converted DNA was PCR amplified using MyTaq HS Mix (Bioline, cat# BIO-25045) and TruSeq PCR Primer Cocktail (Illumina, TruSeq Nano cat# FC-121– 4001). Amplified libraries were purified twice with an equal amount of SPRI beads to remove primer and adapter dimers. The resulting reduced-representation libraries were pooled in batches of 12, and single-end, 100-bp sequenced in multiple lanes of an Illumina HiSeq 2000 v3 according to the standard Illumina protocol. Bisulphite sequencing was conducted at core facilities of the Broad Stem Cell Center, UCLA.

#### Whole Genome Bisulphite Sequencing

We extracted total genomic DNA from the 40 *Q. lobata* trees by grinding leaf tissue in liquid nitrogen using a Mixer Mill MM301 (Retsch, Germany). We used a prewash method (Li *et al*., 2007) followed by a modified CTAB protocol (Doyle & Doyle, 1987). The prewash method was repeated up to 3x until a clear supernatant was achieved. The resultant pellet was then used in a modified CTAB protocol in which the chloroform-isoamyl (24:1) step was repeated twice. DNA was quantified using the Qubit dsDNA BR Assay Kit on the Qubit 3.0 Fluorometer (Life Technologies, Carlsbad, CA).

Total genomic DNA at a concentration of 500 ng in 60uL was sonicated using an S2 Focused-ultrasonicator (Covaris, Woburn, MA, USA) for 60 sec to obtain fragments in the 200-300 bp range (Duty cycle: 10%, intensity: 5, cycles/burst: 200, Mode: Frequency sweeping). Kapa DNA hyper kit (Kapa Biosystems, Inc., MA, USA) was used to repair the ends of the DNA fragments and to ligate the DNA fragments with Illumina TruSeq DNA LT Nano adapters. Subsequently, the adapter-ligated DNA was treated with the Epitect bisulphite kit (Qiagen, Hilden, Germany). Finally, bisulphite-treated DNA was PCR amplified once with MyTaq DNA polymerase (Bioline, MA, USA) at 15 cycles.

#### Transcriptome sequencing

To conduct RNA-seq, total RNA was extracted from the same tree as used in WGBS (Table S1). Three separate RNA extractions were performed using a modified version of the Conifer RNA prep protocol from the Cronn Lab and the Spectrum Plant Total RNA kit (Sigma, St. Louis, MO, USA). Leaf tissues were flash frozen in liquid nitrogen and ground using a Mixer Mill MM301 (Retsch, Germany). Powdered tissues were transferred to cold 2mL tubes, and 1.8 mL of cold RNA Extraction Buffer + DTT was added. RNA Extraction Buffer consists of 8 M Urea, 3 M LiCl, 1% polyvinylpyrrolidone K-60, 5 mM DTT (added just before use, 1M stock). The tubes were then vortexed for 30 sec, incubated at 4° for 30 min, then centrifuged at 4° for 30 min at 20,000 x *g*. The supernatant was discarded and the pellet was used as the starting material for the Spectrum Plant Total RNA kit, protocol A, adding 750uL of Binding Solution, and performing on-column DNase I digestion. RNA quality and quantity were assessed using the Agilent RNA ScreenTape System on the Agilent 2200 TapeStation system (Agilent Technologies, Santa Clara, CA, USA).

RNAseq libraries were constructed using the TruSeq RNA Library Prep Kit v2 (Illumina, San Diego, CA, USA) following the Low Sample (LS) Protocol in the TruSeq RNA Sample Preparation v2 Guide. Briefly, mRNA was purified and fragmented from total RNA at a concentration of 1 µg in 50 µL using poly-T oligo-attached magnetic beads. First-strand and second-strand cDNA synthesis, end repair, dA-tailing, ligation, purification, and enrichment steps were performed following the manufacturer’s instructions. Libraries were analyzed using the Agilent D1000 Screen Tape System on the Agilent 2200 TapeStation (Agilent Technologies, Santa Clara, CA, USA). Libraries were quantified using the Qubit dsDNA BR Assay Kit on the Qubit 3.0 Fluorometer (Life Technologies, Carlsbad, CA). Libraries were pooled and sequenced on the Illumina HiSeq 4000 sequencer using paired-end 100 bp read in August 2018.

### Data Processing

#### Reduced-Representation Bisulphite Sequencing (RRBS) Read processing and alignment

Raw qseq files were converted to FASTQ with EAMSS correction and demultiplexed. Reads were trimmed with Trim Galore using ‘--quality 20 --length 80 --rrbs --clip_R1 3’. The ‘—rrbs’ option conditionally removes 2 bp from the 3′ end of reads where adapter is detected, eliminating the filled-in non-genomic cytosine introduced during MspI end repair; ‘--clip_R1 3’ additionally hard-clips 3 bp from the 5′ end of every read. Trimmed reads were aligned with bwa-meth v0.2.0 to the *Quercus lobata* reference assembly (Sork et al., 2022; NCBI Accession: GCF_001633185)), with phiX sequence appended (Qlobata.v3.0.phiX.fasta), using default parameters.

#### Methylation calling

Per-cytosine methylation was called with MethylDackel extract, run separately for each sequence context. Position-specific inclusion bounds were tuned empirically from MethylDackel ‘mbias’ plots generated for each cohort, giving ‘--nOT 4,2,0,0 --nOB 2,4,0,0’ for CG, ‘--nOT 4,0,0,0 --nOB 0,4,0,0’ for CHG, and ‘--nOT 2,0,0,0 --nOB 0,2,0,0’ for CHH. All contexts used ‘--maxVariantFrac 0.2 --minOppositeDepth 2’ to exclude positions where the opposite-strand base call indicated a likely C→T sequence variant rather than an unmethylated cytosine, together with ‘-q 10’ (minimum MAPQ) and ‘-p 5’ (minimum base quality). Duplicate reads were not removed. Because MspI cuts at fixed sites, independent RRBS fragments share identical start coordinates, so coordinate-based deduplication would discard genuine reads.

#### Bisulphite conversion efficiency

Non-conversion was estimated from cytosines on approximately 42 chloroplast-derived scaffolds, which are natively unmethylated; no exogenous spike-in control was used. Mean non-conversion was 0.39% for the field adults (range 0.27– 0.68%), 0.82% for Chico progeny, and 0.83% for Placerville progeny. Progeny from both gardens used in the between-garden comparison were sequenced together on a single run, so the between-garden comparison is not confounded by conversion efficiency or sequencing batch.

#### Site filtering

Positions were retained if covered by at least 10 reads and if coverage did not exceed a per-sample upper bound computed from the coverage distribution. Positions with more than 20% missing data across the samples used in each test were excluded. Remaining values were capped using a Tukey-style outlier rule: the 10th and 90th percentiles of each position’s distribution were taken as box edges, a range was computed as the difference between them, fences were placed at 1.5 × that range beyond each edge, and values outside the fences were reset to the nearest observed value inside them. Finally, positions were retained only if they exceeded a minimum-variation threshold, set separately by context at 0.045 (CG), 0.024 (CHG) and 0.011 (CHH). These thresholds were derived empirically as the mean per-position standard deviation across the two garden cohorts, computed after the missing-data filter. Sample sets were intersected before the minimum-variation threshold was applied, so that positions varying in only one of the two sample sets were retained rather than discarded. This method applies to the parent–offspring comparisons, for which each garden was intersected with the maternal set separately, giving the site counts reported in Table S2.

#### Methylomes using whole genome bisulphite sequencing

Whole-genome bisulphite sequenced (WGBS) libraries were pooled and sequenced on an Illumina HiSeq 4000 machine. The reads were aligned to the *Quercus lobata* genome (Sork *et al*., 2022), using BSseeker2 (Guo et al., 2013) and bowtie with default parameters. After alignment, BSseeker2 was used again to call methylation with default parameters. CgmapTools (Guo et al., 2018) were then used to analyze the methylation levels for each sample. DNA methylation was analyzed separately for each sequence context using per-cytosine methylation fractions. Cytosines were assigned to three genomic features based on gene models: the gene body; 1.5 kb upstream; and 1.5 kb downstream. Mean methylation for each feature was calculated as the mean of non-missing cytosine methylation fractions. Features containing < 3 cytosines were treated as missing. Additionally, weighted methylation levels were calculated in 100 bp windows reflecting the sum of methylated cytosines divided by the sum of all cytosines / thymines. Sequence data were aligned to the ValleyOak3.2 reference genome (GCF_001633185.2), and analyses were restricted to the 12 assembled chromosomes. Gene coordinates were obtained from the NCBI LOC annotation.

#### Transcriptomes using RNAseq data

Raw sequenced reads were converted from qseq to fastq format, reads failing the Illumina quality filter were removed, and the reads were demultiplexed using custom scripts (available at github.com/alaynamead/RNAseq_scripts). Sample quality was checked using FastQC 0.11.8 (Andrews, 2010). Cutadapt 2.3 (Martin, 2014) was used to trim adapters and low-quality ends (using a cutoff quality score of 27), and the reads shorter than 30 bp after trimming were removed. Reads were aligned to Valley oak genome v3.2 (NCBI Accession: GCF_001633185) using STAR 2.7.1 (Dobin et al., 2013). Reads which mapped to multiple locations within the genome (quality score < 255) were removed using Samtools 1.9 (Li *et al*., 2009).

### Statistical Analyses

#### Prediction 1. DNA methylation is environmentally induced

First, we compared mean genome-wide methylation levels separately for each context between 80 five-year-old trees (two half-sib progeny per family per garden; see sample details in Table S3 B, C) growing in each of the two common gardens. If methylation is environmentally induced, methylation levels within a family should differ between the two gardens. We therefore calculated the mean methylation level for each tree across the methylome and tested for differences between gardens using paired *t*-tests, with family as the pairing factor.

Second, we assessed whether the mean genome-wide levels of CG, CHG, and CHH methylation changed across years for 21 trees growing in the Chico Garden (see sample details in Table S3 D, E). We used paired *t*-tests to compare mean methylome-wide methylation levels for the same at the two time points, separately for CG, CHG, and CHH methylation.

Third, and most importantly for Prediction 1, we identified DMPs within each of the sequence contexts that showed evidence of environmentally induced methylation differences. We refer to these DMPs as “DMP plasticity outliers”. For each DMP, we first calculated the mean methylation percentage of the two progenies from each family within each garden and then calculated the difference in mean methylation between gardens for each of the 40 families. We tested these paired family-level differences using paired *t*-tests, separately for each sequence context. DMP plasticity outliers were identified using a stringent significance criterion of *P* < 0.0001.

#### Prediction 2. Methylation is transmitted across generations

To test whether DNA methylation of *Q. lobata* was transmitted across generations, we correlated the methylation level at each DMP between the maternal parent and the mean level of their offspring, separately for each context and garden, re-estimated Spearman’s Rho, and pooled the resulting null values for each context and garden nonparametric correlation test (Smeeton et al., 2025). This correlation test is advantageous because it does not require the assumption of normality between generations. To test and compare the observed genome-wide patterns of correlations with the null hypothesis that DNA methylation is random between parents and offspring, we permuted maternal methylation values across families within each DMP, separately for each context and garden, re-estimated Spearman’s Rho, and pooled the resulting null values for each context and garden. This breaks the family structure while preserving the distribution of methylation values, creating a null distribution under the hypothesis of no parent-offspring correlation. Given the computational time required for such a large number of loci, we used only 100 permutations to observe the general patterns. We also examined the individual p-values for each correlation test.

Parent-offspring regression can also be used to estimating narrow sense heritability, *h*n^2^, (Falconer & Mackay, 1983). We were concerned about the non-normal distribution of observed values of *h*n^2^ for the three sets of DMPs and also whether epigenetic data meet the assumptions of quantitative genetic models. Nonetheless, for heuristic purposes, we conducted parent-offspring regressions and repeated the permutation tests. The findings are reported in the Supplemental Information.

#### Prediction 3. Induced patterns remain stable after transmission

For DNA methylation patterns to be consistent with the Lamarckian expectation of the inheritance of acquired characteristics, the DMP plasticity outliers identified in Prediction 1 should be transmitted from parent to offspring, as assessed in Prediction 2. Because applying equally stringent probability thresholds to both plasticity and evidence of heritability would result in combined probabilities too small to identify potential candidate DMPs, we used relaxed criteria for evidence of heritability. Specifically, we required P < 10⁻⁴ for plasticity combined with evidence of heritability based on either a correlation at P < 0.1 in both gardens or on a correlation at P < 0.05 in one garden. Thus, these two criteria yielded a combined P values of P < 10⁻⁶ and P < 5 × 10⁻⁶, respectively. For DMPs meeting these criteria, we reported functional annotations for the nearest genes and repeats using the published genome assembly, Valley oak 3.2 (Sork *et al*., 2022) (Accession: GCF_001633185), and relative expression values using data from Prediction 4 (see below).

#### Prediction 4. DNA methylation is associated with gene expression, varying by sequence context and TE position

##### Transcriptome analysis

Gene expression was quantified from the mapped libraries with Salmon v.1.5.2 (Patro *et al*., 2017) with transcript-level counts summarized to gene level. matrix. Genes with expression >= 10 in at least 25% trees were retained and gene-level read counts were first normalized for sequencing depth across all samples, then expression values were log2(x+1) transformed for all downstream analyses.

##### Methylome analyses

Genome-wide methylation profiles were generated using DeepTools 3.5.1 ‘computeMatrix scale-regions’ (Ramírez et al., 2016). To examine relationships between methylation and gene expression, genes were ranked independently within each tree according to expression level. The top and bottom 20% expressed genes were compared by calculating mean methylation for each genomic feature (upstream, gene body, downstream) within each tree. Differences between the two expression classes were evaluated using paired *t*-tests across the 42 trees and effect sizes were calculated as the median paired difference with 95% confidence intervals. To assess whether these relationships extended across the full expression range, we additionally divided genes into ten equal-sized expression deciles within each tree and calculated mean methylation for each.

##### Transposable element analyses

TE annotations were obtained from the published annotation of the ValleyOak3.2 genome (Sork et al., 2022). For each gene, TE presence was determined independently for the gene body, upstream, and downstream regions, with any overlap classified as TE present. Associations between TE presence and DNA methylation were evaluated by comparing methylation profiles and mean methylation levels between genes containing and lacking TEs within each genome feature. To investigate interactions between TEs and gene expression, genes were further classified according to both expression level (top and bottom 20%) and TE presence. We also ranked genes by their distance to the nearest upstream or downstream TE and divided them into ten equal-sized distance bins. Mean CHH methylation was calculated within each distance bin, and associations were quantified between TE distance and methylation using Spearman rank correlations.

## Results

### DNA methylation is induced by the environment

The levels of genome-wide DNA methylation per family differ across CG-, CHG-, and CHH DMPs, with CHH levels being the lowest (∼60%, ∼20%, and ∼8-16 %, respectively for field adult and progeny garden samples; Figure 2A). Mean levels of CG- and CHG methylation per family was similar between field adults and their garden progeny and did not differ between progeny growing in the two common gardens (paired *t*-test, df=39, P > 0.10). In contrast, the CHH-DMP levels in the progeny were lower than those of the field adults and were significantly higher in the Chico than Placerville Garden (paired *t*-test, df=39, *P* = 3.1 x 10^-11^, Figure 2A). Between years, mean family levels of mCG or mCHG were similar between the end of one season and the beginning of the following season, whereas the mCHH levels differed between seasons (Figure 2B). Thus, CHH-methylation was consistently the least stable context. None of these patterns were attributable to differences in sequencing depth or genome coverage.

**Figure 2.**
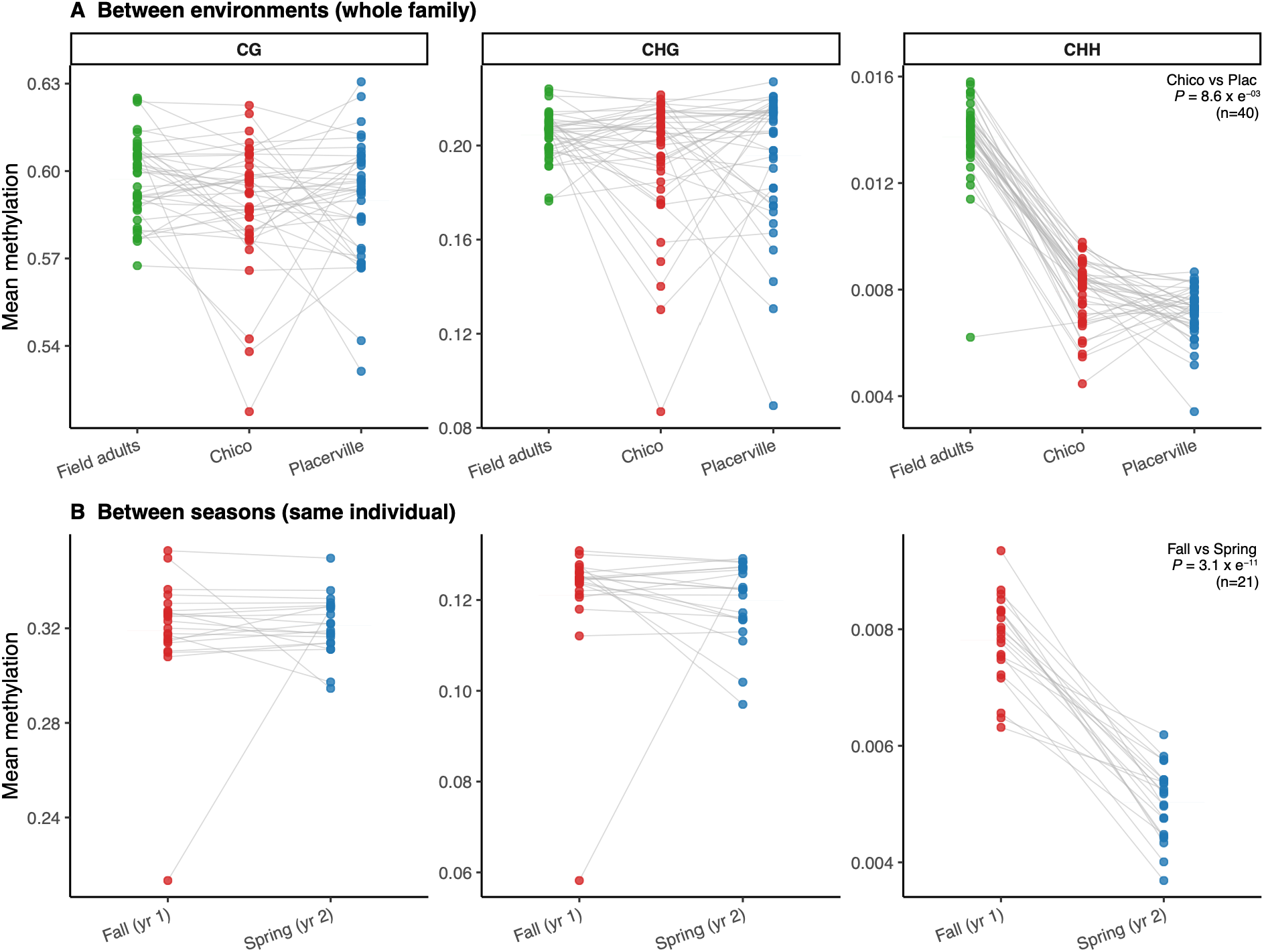
Mean genome-wide methylation across all differentially methylated positions (DMPs) for each methylation context (CG, CHG, CHH). (**A**) Comparison of methylation among 40 field-grown adult trees (green) and the corresponding half-sib progeny grown in the Chico (red) and Placerville (blue) common gardens. Points represent family means and grey lines connect the same family across environments. CHH methylation differed significantly between the two common gardens (paired *t*-test, n = 40, *P* = 8.6 x 10^-3^), whereas CG and CHG methylation did not differ. Field-grown adults are included as a reference for natural methylation levels. (**B)** Seasonal comparison of methylation within the same individuals sampled in Fall of year 1 and Spring of year 2 (n = 21). CHH methylation declined significantly between sampling points (paired *t*-test, *P* = 3.1 x 10^-11^), whereas CG and CHG methylation did not change. Note that the y-axis scales differ among contexts.

Our tests that for differences between the two gardens within families identified 84 individual DMPs that showed evidence of environmentally responsive methylation (paired *t*-test, df=39, P < 10^-4^; Figure 3, Table S3): most of these loci (71%) were in the CHH sequence context. We found 13 CG-DMP and 9 CHG-DMP outliers, and all of which had higher methylation levels at the Placerville garden (Figure 3A-B). Among the 62 CHH-DMP plasticity outliers, all but four had higher methylation in the Chico Garden (Figure 3C). The higher frequency of mCHH outliers suggests that this context to most likely to exhibit plasticity across environments.

**Figure 3.**
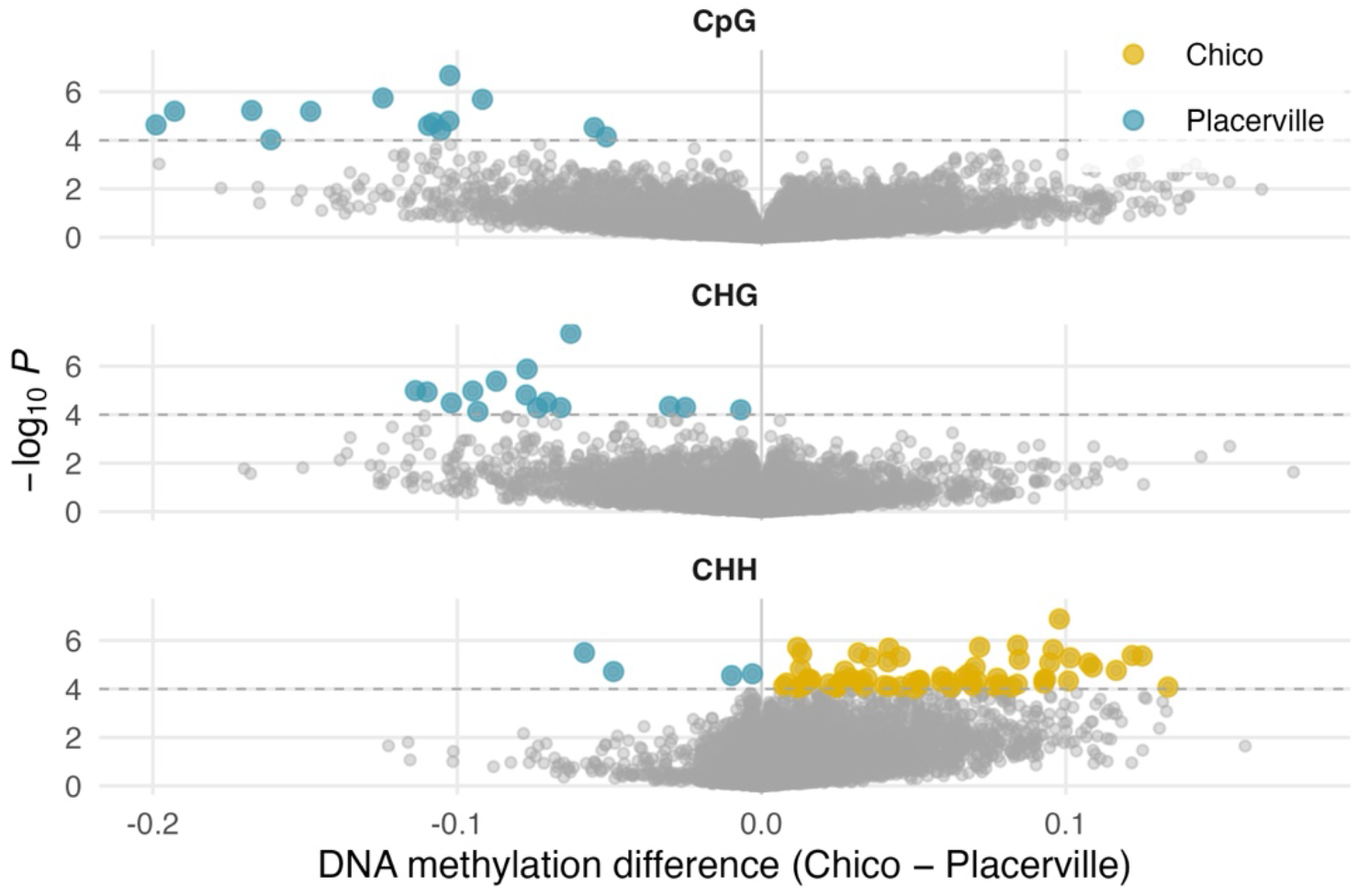
Volcano plots showing the difference in mean methylation between half-sib progeny grown in the Chico and Placerville common gardens (Chico – Placerville) against statistical significance (-log10P) for CpG, CHG, and CHH DMPs. Each point represents a single DMP; samples sizes are 12,266 CG DMPs; 17,461 CHG DMPs, and 145,913 CHH DMPs. Colored points indicate DMPs that exceeded the significance threshold (*P* < 0.0001) and showed higher methylation in either the Chico (gold) or Placerville (blue) common garden; grey points were not significant. The horizontal dashed line denotes the significance threshold (P = 0.0001). DMP outliers identified at *P* < 0.0001 are listed in Table S2.

To determine whether methylation levels were coordinated among sequence contexts, we ranked each tree according to its maximum methylation level within each sequence context. Peak CG and CHG methylation were strongly correlated among trees (Spearman’s rho = 0.95, *P* < 0.001), whereas peak CHH methylation was not correlated with either CG (rho = 0.12, *P* =0.45) or CHG methylation (ρ = 0.11, *P* = 0.49). These results indicate that CHH methylation varies independently of the more stable CG and CHG methylation landscapes among individuals.

#### DNA methylation is transmitted from parent to offspring

Of the three sequence contexts, only CG-DMPs had an overall distribution of parent-offspring correlation that was dramatically different than a randomly sampled distribution (Figure 4A). Out of 13,547 positions at Chico and 13,716 at Placerville, we found 322 and 363 CG-DMPs at Chico and Placerville gardens, respectively, that were significant (Spearman’s Rho, df=39, P < 10^-4^; Table 1). At both gardens, the distributions of observed Spearman’s correlations include some CHG DMPs that fall outside of the randomly generated distributions (Figure 4B), with 33 and 40 DMPs showing significant correlations (Spearman’s Rho, df=39, *P* < 10^-4^; Table 1). The observed CHH-DMP distributions of Spearman’s Rho almost completely overlapped with the random distribution (Figure 4C), with 34 DMPs at both gardens showing significant correlations ((Spearman’s Rho, df=39, P < 10^-4^; Table 1). Genome-wide patterns of narrow sense heritability (*h*^2^) showed the same trends for sequence contexts, except that their distributions are highly and symmetrically leptokurtic (Figure S1). Overall, the associations between parent and offspring indicate that of the three sequence contexts, CG-DMPs are the most likely to be heritable from parent to offspring.

**Figure 4.**
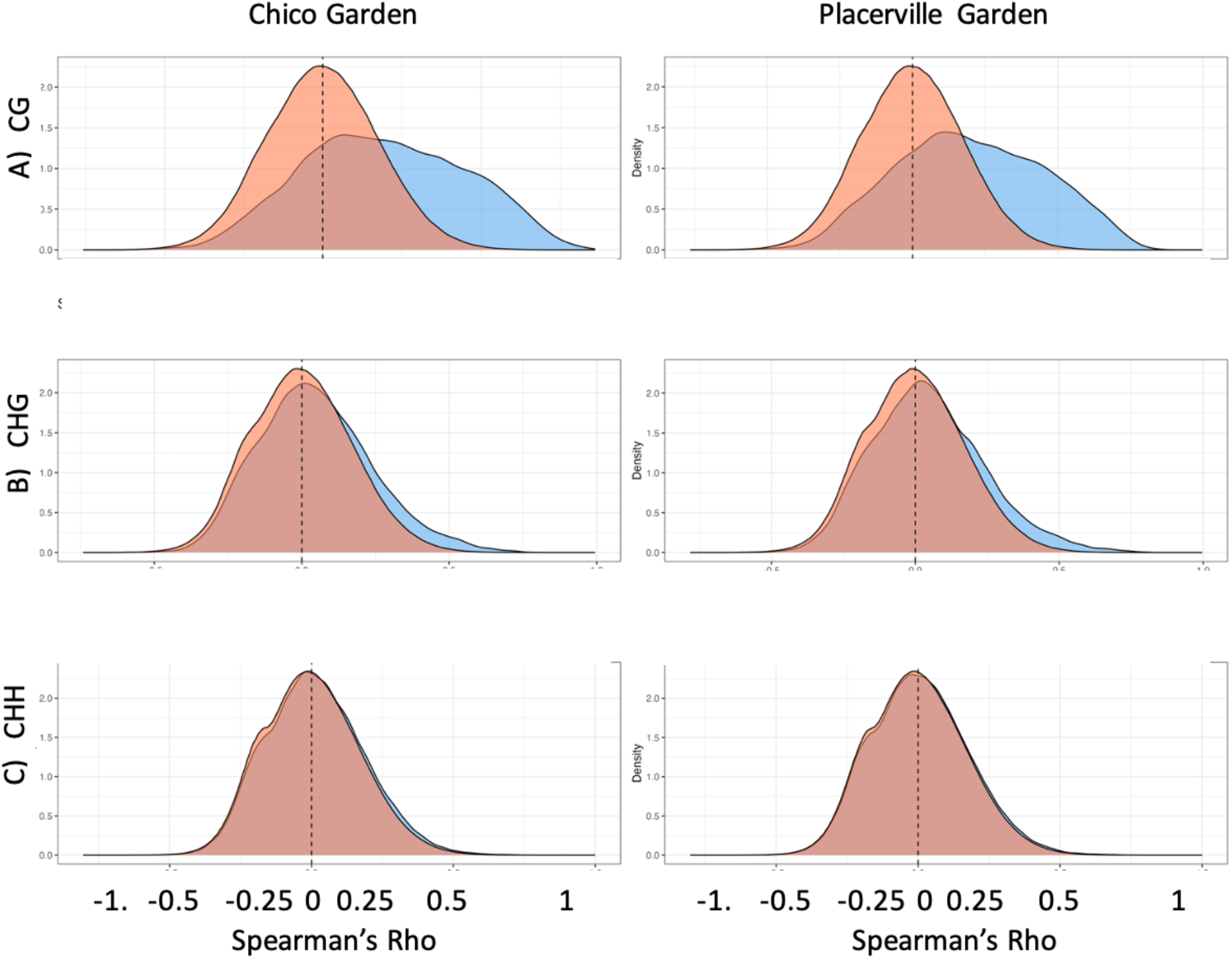
Frequency distribution of observed correlations been methylation level in parent and offspring (blue) versus random correlations (red) between methylation levels separately for (**A**) CG methylation, (**B**) CHG methylation, and (**C)** CHH methylation (bottom row).in leaves from maternal tree collected from species-wide natural population and progeny growing in Chico (left column) and Placerville (right column) gardens.

**Table 1.** Summary of significant Spearman’s Rho correlations (*P* < 0.0001) in levels of methylation between maternal seed source and the mean progeny value of 40 trees growing in the Placerville common garden. (See Table S1 for description of trees.

|  | Context | # total DMPs | # significant correlations | % significant correlations |
| --- | --- | --- | --- | --- |
| **Chico** | CG | 13,547 | 322 | 82.8% |
|  | CHG | 28,304 | 33 | 8.5% |
|  | CHH | 188,956 | 34 | 8.7% |
|  | Total | 230,807 | 389 |  |
| **Placerville** | CG | 13,716 | 363 | 83.1% |
|  | CHG | 28,848 | 40 | 9.2% |
|  | CHH | 193,153 | 34 | 7.8% |
|  | Total | 235,717 | 437 |  |

#### Environmentally induced methylation patterns are rarely stably inherited

Of the 84 putatively environmentally induced DMP outliers among the 22,586 DMPs tested for significant differences between paired progeny sets across the two common gardens (Table S3), we identified only 15 DMPs that met both criteria for pattens consistent with Lamarckian-like evolution (Table 2): evidence of environmental induction (i.e., a plasticity outlier at P < 1 x 10^-4^) and a significant parent-offspring methylation correlation (i.e., P < 0.1 in both gardens or P < 0.05 in one garden). These candidate DMPs were most commonly CHH-methylated: 2 CG-, 2 CHG-, and 11 CHH DMPs (Table 2). Using transcriptomes from 42 trees sampled in the Placerville common garden, we found detectable gene expression for genes nearest the 11 of the 15 DMPs.

**Table 2.** Subsample of 15 DMPS out of 84 plasticity outliers that met the threshold of P < 0.0001 (See Table S3), their locations to closes genes and repeats, and the correlation tests as evidence of transgenerational inheritance that are significant at either the Placerville or Chico Gardens with P < 0.05 or both with P < 0.10. Significant values are bolded. DMP outliers are sorted by context. We grouped genes into four expression categories from RNAseq data of trees growing in the Placerville Garden. Using mean depth-normalized (log2) expression for each gene, we classed the top 20% as High and the bottom 20% as Low, with Moderate covering all genes in between. Silent denotes genes that did not pass the expression filter and were excluded. Ty3 TEs were formally known as Gypsy (see Wei *et al*., 2022).

| Chr. | position | Con-<br>text | Plasticity outliers |  | Parent-offspring correlation |  |  |  | closest<br>gene | gene function | distance<br>from<br>gene | closest<br>repeat<br>family | closest repeat<br>superfamily | distance<br>from<br>repeat | Gene<br>expression |
| --- | --- | --- | --- | --- | --- | --- | --- | --- | --- | --- | --- | --- | --- | --- | --- |
|  |  |  | paired t-test |  | Placerville |  | Chico |  |  |  |  |  |  |  |  |
|  |  |  | t-stat | P value | Rho | P-value | Rho | P-value |  |  |  |  |  |  |  |
| 6 | 16286242 | CG | -5.6367 | 0.000002 | -0.084 | 0.6095 | 0.360 | 0.0265 | LOC-115994925 | Cellulose<br>synthase-like<br>protein G1 | -3231 | rnd-<br>4_family-23 | LINE_L1 | 0 | high |
| 11 | 18274950 | CG | -4.7881 | 0.000024 | 0.369 | 0.0190 | 0.386 | 0.0140 | LOC-115968872 | 25.3 kDa<br>vesicle<br>transport<br>protein<br>SEC22-1 | 0 | s1RF0451 | Tandem_(C)n | 184 | medium |
| 4 | 60006623 | CHG | -4.9379 | 0.000033 | 0.192 | 0.2687 | 0.594 | 0.0005 | LOC-115988013 | Auxin<br>response<br>factor 6 | 0 | rnd-<br>1_family-228 | LINE_L1 | 508 | silent |
| 6 | 16286267 | CHG | -5.0436 | 0.000012 | 0.004 | 0.9824 | 0.377 | 0.0196 | LOC-115994925 | Cellulose<br>synthase-like protein G1 | -3206 | rnd-<br>4_family-23 | LINE_L1 | 0 | high |
| 2 | 11547789 | CHH | 5.1439 | 0.000008 | 0.009 | 0.9555 | 0.334 | 0.0378 | LOC-115977973 | Snakin-2 | 1612 | rnd-<br>1_family-267 | Unknown<br>unclassified | 0 | moderate |
| 2 | 11547801 | CHH | 5.3237 | 0.000005 | 0.073 | 0.6614 | 0.326 | 0.0454 | LOC-115977973 | Snakin-2 | 1600 | rnd-<br>1_family-267 | Unknown<br>unclassified | 0 | moderate |
| 4 | 39038509 | CHH | 4.8705 | 0.000040 | 0.706 | 0.0000 | 0.607 | 0.0002 | LOC-115985165 | . | -515 | s1RF0016 | Tandem_(A)n | -109 | moderate |
| 5 | 7557828 | CHH | 4.6036 | 0.000053 | 0.319 | 0.0659 | 0.331 | 0.0518 | LOC-115992010 | AP2-like<br>ethylene<br>-responsive<br>transcription<br>factor AIL1 | -5885 | rnd-<br>1_family-442 | DNA_MuLE-MuDR | -760 | silent |
| 6 | 1116145 | CHH | 4.8830 | 0.000046 | 0.458 | 0.0109 | 0.212 | 0.2362 | LOC-115994608 | . | -1362 | rnd-<br>6_family-165 | Unknown<br>unclassified | 0 | silent |
| 6 | 8637559 | CHH | 4.4685 | 0.000079 | 0.223 | 0.2043 | 0.343 | 0.0352 | LOC-115994083 | Acyl<br>transferase 4 | -191 | rnd-<br>1_family-359 | Unknown<br>unclassified | -39 | medium |
|  |  |  | paired t-test |  | Placerville |  | Chico |  |  |  |  |  |  |  |  |
|  |  |  | t-stat | P value | Rho | P-value | Rho | P-value |  |  |  |  |  |  |  |
| 7 | 157616 | CHH | -5.0123 | 0.000019 | 0.325 | 0.0527 | 0.351 | 0.0359 | LOC-115954121 | Zinc finger AN1 and C2H2 domain-containing stress-associated protein 11 | -74 | s1RF0016 | Tandem_(A)n | -64 | high |
| 7 | 157627 | CHH | -5.0749 | 0.000016 | 0.527 | 0.0010 | 0.449 | 0.0060 | LOC-115954121 | Zinc finger AN1 and C2H2 domain-containing stress-associated protein 11 | -85 | s1RF0016 | Tandem_(A)n | -53 | high |
| 7 | 27807409 | CHH | 5.5392 | 0.000003 | 0.259 | 0.1169 | 0.340 | 0.0490 | LOC-115951832 | . | 660 | rnd-5_family-3370 | LTR_Ty3* | 525 | n/a |
| 9 | 36665030 | CHH | 4.8954 | 0.000044 | 0.141 | 0.4410 | 0.407 | 0.0187 | LOC-115959226 | . | -2564 | rnd-1_family-69 | Unknown unclassified | 0 | very high |
| 12 | 22354830 | CHH | 5.0224 | 0.000017 | 0.196 | 0.2385 | 0.358 | 0.0322 | LOC-115972330 | Adenylate isopentenyltransferase 3, chloroplastic | -1209 | rnd-1_family-396 | Unknown unclassified | 0 | moderate |

Among these 15 positions, the most compelling candidate was the Zinc finger AN1 and C2H2 domain-containing stress-associated protein 11, which is associated with stress response (Giri *et al*., 2011). This gene is proximal to two CHH-DMPs, which are located 1and 12 bp upstream of the gene, and 53 and 64 bp, respectively, from a repeat (Table 2). In addition, the gene was highly expressed in the trees sampled in Placerville (Table 2).

One other candidate gene linked to DNA methylation in Table 2, with significant parent-offspring correlation in both gardens, is 25.3 kDa vesicle transport protein SEC22-1. The associated CG-DMP is located within the gene and 184 bp from the nearest repeat. We found a third gene on chromosome 4, LOC-115985165, which showed moderate expression in the following year, although its function is unknown.

We found another DMP of possible interest located on chromosome 5 that exhibited significant parent-offspring correlation in both gardens. The nearest gene, the AP2-like ethylene-responsive transcription factor AIL1, is located -5885 bp upstream from the CHH-DMP, whereas and the nearest DNA_MuLE-MuDR repeat is 760 bp upstream. Moreover, gene expression was indetectable, although that evidence alone would not eliminate this DMP as a candidate. Nonetheless, this DMP does not appear to be a strong candidate for Lamarckian-like evolution.

Thus, this analysis of environmentally responsive DMPs detected by RRBS identifies at least one strong candidate stress-response gene that may warrant further study as a potential example of environmentally induced, parent-offspring-associated CCH-methylation in a Lamarckian-like process.

#### DNA methylation is associated with gene expression and varies by sequence context and TE position

The whole-genome methylation profiles across 44,957 genes in 42 seven-year-old valley oak trees growing in the Chico garden (Figures 5A-C) were consistent with previous reports for valley oak and other angiosperms (Niederhuth *et al*., 2016; Sork *et al*., 2022): CG methylation was depleted at the start and stop sites of genes while peaking across gene bodies; CHG methylation was reduced within the gene body; and CHH methylation concentrated immediately upstream of the transcription start site (TSS). The 42 trees exhibited conserved methylation profiles, but individual trees varied in their levels of methylation (Figure 5). The differences were greatest within the gene-flanking regions, whereas methylation levels within gene bodies were comparatively consistent among trees (Figures 5A-C). Differences among individuals therefore primarily reflected changes in methylation intensity rather than in spatial distribution across genes.

**Figure 5.**
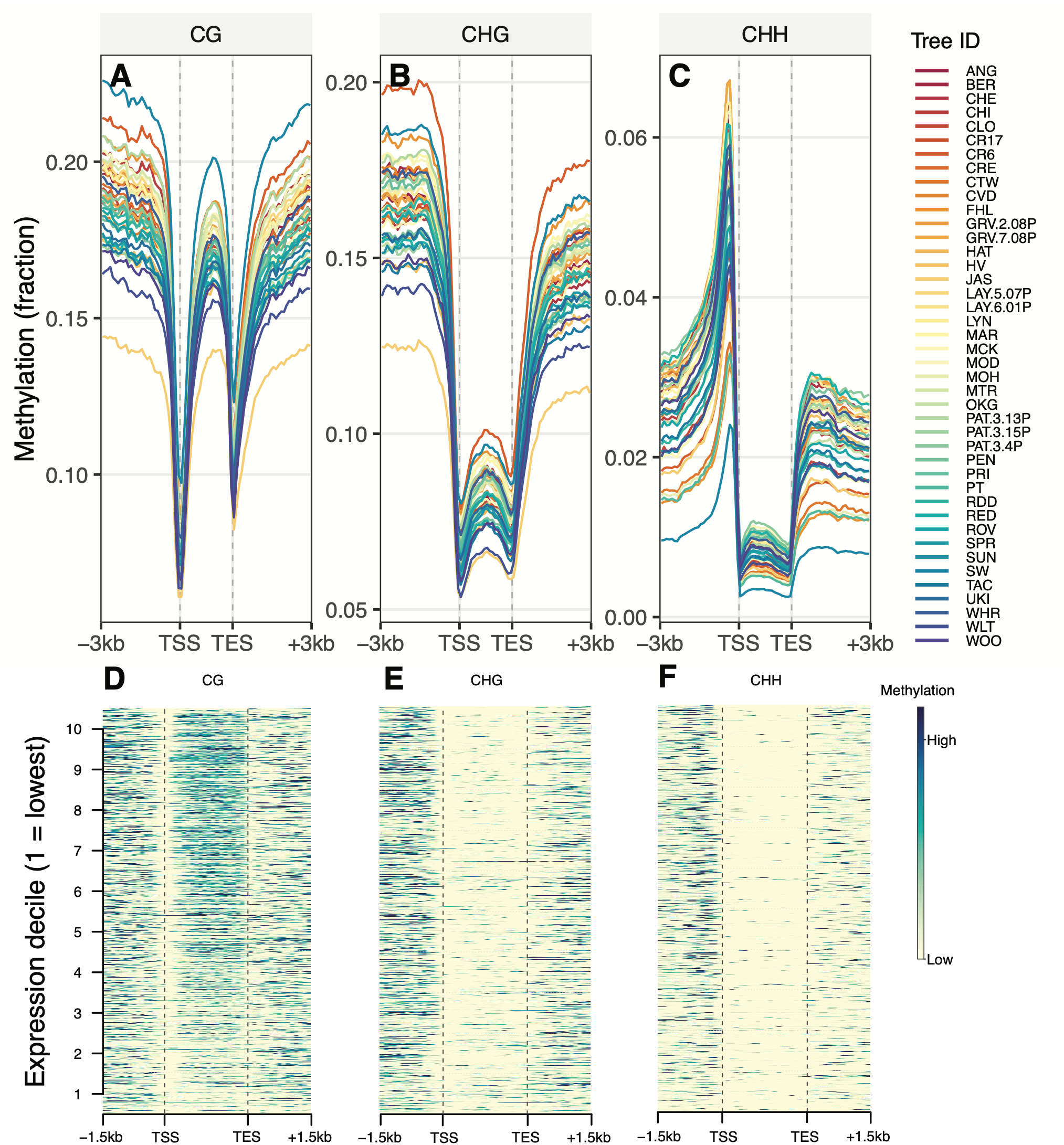
Genome-wide DNA methylation profiles across genes and their relationship with gene expression in 42 *Quercus lobata* trees in the Placerville garden. (A-C) Mean metagene methylation profiles for CG, CHG, and CHH methylation across 44,957 annotated genes in 42 eight-year-old trees grown. Methylation levels are shown across gene bodies together with the 3 kb up- and downstream flanking regions. Each colored line represents one tree. Vertical dashed lines denote the transcription start site (TSS) and transcription end site (TES). (**D-F**) Heatmaps showing mean methylation levels across genes ranked by expression decile (1 = lowest expression, 10 = highest expression) for CG, CHG, and CHH methylation. Methylation values represent the mean across all trees ranked by expression decile.

##### Associations with specific sequence context

The relationship between DNA methylation and gene expression differed markedly among the three sequence contexts (Figure 5D-F). We observed a positive association between gene expression and CG methylation within transcribed regions (Figure 5D), and between gene expression and CHH methylation immediately upstream of the TSS (Figure 5F), whereas CHG methylation showed no such pattern. To quantify these relationships, we compared mean methylation levels for the 20% most highly expressed and 20% least expressed genes within each of the 42 trees (Figure S2). For CG methylation, highly expressed genes showed significantly greater gene body methylation than weakly expressed genes (median difference = +0.165, *P* < 2 x 10^-47^, paired *t*-test across 42 trees, Figure S2), while downstream methylation was significantly lower in highly expressed genes. No significant difference was observed within the upstream flanking region. In contrast, CHG methylation showed the opposite pattern within gene bodies, with highly expressed genes exhibiting significantly lower methylation than weakly expressed genes (median difference -0.039, *P* < 2 x 10^-38^, paired *t*-test, Figure S2). Highly expressed genes also showed slightly higher upstream CHG methylation but significantly reduced downstream CHG methylation. CHH methylation was associated almost exclusively with the upstream flanking region. Highly expressed genes exhibited significantly greater CHH methylation within 1.5 kb upstream of the TSS than weakly expressed genes (median difference +0.033, *P* < 5 x 10^-32^, paired *t*-test, Figure S2), whereas methylation levels within gene bodies and downstream regions differed little between expression classes.

Thus, these findings reveal that the association between DNA methylation and gene expression is strongly context dependent in *Q. lobata.* High gene expression is associated with elevated CG methylation within gene bodies, reduced CHG methylation across gene bodies, and increased CHH methylation in the immediate upstream gene-flanking regions.

##### Relationship with proximity of TEs

We compared methylation profiles between genes containing repeats, which are largely TEs, and genes lacking repeats (Figure 6A, Figure S3). Repeat-containing genes consistently exhibited higher methylation than repeat-absent genes across all three sequence contexts, although the magnitude of this effect differed. CG methylation showed the strongest positive association with repeat presence, particularly within gene bodies where methylation approximately doubled in repeat-containing genes (median difference = +0.229, *P* < 5 x 10^-81^, paired *t*-test across 42 trees, Figure S3). CHG methylation exhibited more modest increases across gene bodies and flanking regions, whereas CHH methylation remained concentrated immediately upstream of the TSS and reached significantly higher levels in repeat-containing genes than repeat-absent genes (median difference = +0.027, *P* < 9 x 10^-15^, paired *t*-test, Figure S3). All three sequence contexts showed a significant positive association between methylation levels and repeats present downstream of genes. These results demonstrate that repetitive sequences influence DNA methylation differently between contexts.

**Figure 6.**
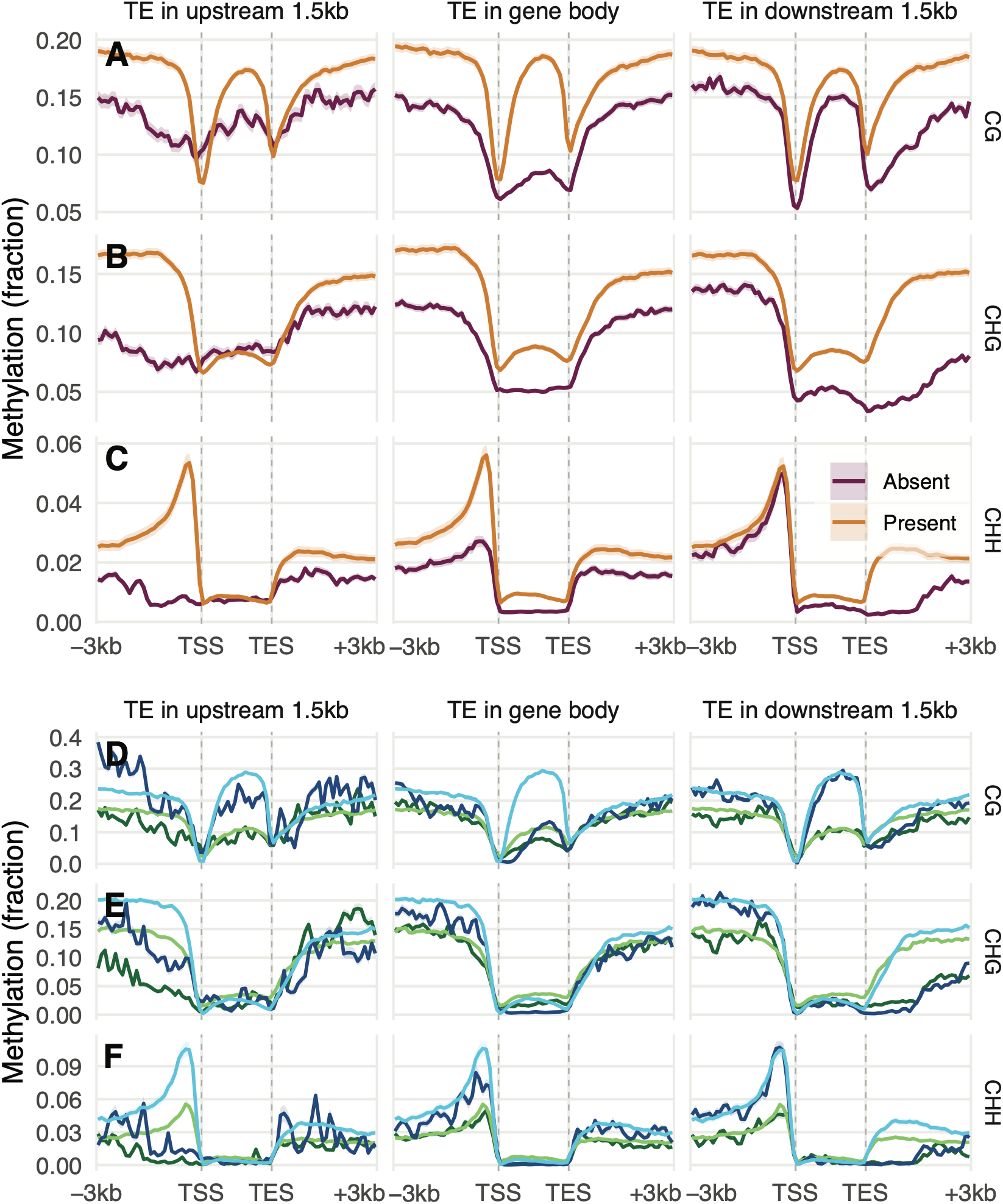
Repetitive sequences modify DNA methylation patterns and their relationship with gene expression in *Quercus lobata.* (**A-C**) Mean methylation profiles for genes containing repetitive sequences (simple repeats or transposable elements (TEs) and genes lacking repeats, shown separately for CG, CHG, and CHH. Profiles represent the mean across 42 eight-year-old trees grown in the Placerville common garden: shaded ribbons indicate 95% confidence intervals. Rows show genes containing repeats within the gene body, upstream 1.5 kb flanking region or downstream 1.5 kb flanking region. Vertical dashed lines indicate the transcription start site (TSS) and transcription end site (TES). **(D-F**) Mean methylation profiles for genes grouped by both expression level (top and bottom 20% of expressed genes) and repeat presence. Profiles are shown separately for CG, CHG, and CHH methylation and averaged across all 42 trees.

Our findings revealed that repeat-associated methylation also varied with transcriptional activity, depending on the presence of repeats. CG methylation exhibited the strongest association between repeat content and gene expression, particularly for highly expressed genes containing repeats (Figure 6B). In contrast, CHG methylation remained associated primarily with gene expression rather than repeat presence. CHH methylation continued to exhibit pronounced peaks immediately upstream of the TSS in highly expressed genes irrespective of repeat content, while gene bodies and downstream regions showed relatively little variation among the four expression-repeat categories. These results indicate that repeat-associated CG methylation and upstream CHH methylation show the strongest relationships with transcription.

The analysis of association between methylation and expression persisted across the full expression gradient using expression deciles, rather than only the upper and lower expression classes (Figure S4). All profiles support the interpretation that the effect of DNA methylation on gene expression is influenced by repeat presence. CG gene body methylation increased progressively with gene expression, whereas CHH methylation increased specifically within the upstream flanking region. In contrast, CHG methylation decreased across gene bodies with increasing gene expression, confirming that the observed relationships represent continuous trends rather than differences confined to the most and least highly expressed genes.

CHH methylation differed among transposable elements classes. We found that CHH methylation in gene flanking region declined progressively with increasing distance from both the upstream (Spearman’s ρ = -0.19, *P* <=8.9 x 10^-259^) and downstream TEs (ρ = -0.16, *P* <= 2.8 x 10^-177^), indicating that genes located closest to TEs exhibit the highest levels of upstream CHH methylation (Figure S5).

Variation in TE body methylation reflected the TE family identity consistently more than TE geometric properties (including length, distance to genes, and genic overlap), accounting for approximately 40% of the residual variance in CHH methylation, compared with only approximately 3% explained by TE geometry (Figure S6A-B). Resolving TEs at the family level, rather than the broader superfamily level, substantially improved prediction of CHH methylation. Among TE families, copy number was positively associated with CHH methylation (partial R^2^ = 0.13, Spearman’s ρ = 0.3, *P* <= 2.8 x 10^-39^), indicating that high copy families tended to be more methylated than low-copy families (Figure S6C). These results indicate that CHH methylation patterns are largely conserved across TE classes and are influenced more strongly by intrinsic properties of individual TE families with, with TE family abundance contributing to this pattern, while size and genic features contributed comparatively little additional explanatory power.

Together, these analyses provide strong evidence that DNA methylation affects gene expression, although the relationships are highly context-specific and depend on proximity to TE. CG methylation is most strongly associated with repeat-containing gene bodies, whereas CHH methylation is concentrated immediately upstream of genes, particularly those located near TEs, suggesting distinct regulatory roles for each methylation context.

## Discussion

“Lamarck’s revenge,” as framed in (Ward, 2018), may be too strong a phrase to describe the role of DNA methylation in the inheritance of environmentally induced characteristics in this *Quercus lobata* study. Our findings suggest that such inheritance is possible, but uncommon. We evaluated two criteria for a “Lamarckian” mechanism: environmental induction of DNA methylation and its transmission from maternal seed source to offspring. Although we identified one stress-associated gene that met both criteria, our reduced representation sequence analysis of thousands of differentially methylated positions provided little evidence that environmentally induced, heritable methylation is widespread. Instead, DNA methylation was highly responsive to environmental conditions, suggesting that it contributes to phenotypic plasticity across environments. Heritable environmentally induced methylation, however, may represent only a small component of this plasticity. However, out inclusion of expression analyses demonstrates that, even when uncommon, such methylation could generate additional heritable variation underlying phenotypic differences on which natural selection could act. Thus, although Lamarck’s revenge may be premature, DNA methylation could still provide trees with an additional route for responding to rapid environmental change.

### DNA methylation can be environmentally induced

Mean methylation levels of genome-wide DNA methylation differed between the two gardens differed between gardens for CHH differentially methylated positions (DMPs), but not for CG- or CHG DMPs. Mean CHH methylation at these positions was significantly higher in the Chico garden than in Placerville across the 40 families, while family averages for CG and CHG DMPs did not differ significantly. Additional evidence for CHH methylation plasticity was the observation that genome-wide CHH methylation was higher in fall than in spring within the same individual trees, while CG and CHG levels within the same tree were constant. A similar pattern was observed in *Quercus robur* (Sanchez-Lucas *et al*., 2025). Because CHH methylation is asymmetric and must be actively re-established following DNA replication (Law & Jacobsen, 2009), its maintenance may require ongoing activity of methylation pathways. Several studies have recently reported a role for CHH methylation in environmental response. In *Arabidopsis thaliana* in the western Himalaya, levels of CHH methylation were more likely to change across populations at different elevations than methylation in other sequence contexts (Singh *et al*., 2024). Studies of stress response have shown that drought treatment can trigger CHH hypermethylation of transposable elements in *Populus nigra* (Peña-Ponton *et al*., 2024) and in *Quercus lobata* (Peck & Sork, 2024), although CHH-methylation did not increase following temperature stress in the asexual plant, *Lemna minor* (Van Antro *et al*., 2023). Nonetheless, these studies, along with ours, support the conclusion that CHH methylation is an important component of environmentally responsive DNA methylation

Our search for differentially methylated positions (DMPs) at which progeny within families differed significantly between gardens revealed 84 highly significant candidate plasticity outliers, 73% of which occurred in the CHH context. Many of the DMPs identified “plasticity outliers” may represent environmentally responsive methylation states. We highlight below DMPs that overlap genes or repeats and are associated with genes with published environmental-response functions that may be relevant to differences between the two gardens.

One example involves the gene Snakin-2, which has 11 CHH-DMP outliers located about 1600 bp downstream of the gene, all of which overlap a repeat (Table S3). This Snakin-2 gene is upregulated during defense responses in tomato (Herbel *et al*., 2017). Another candidate plasticity outlier is a CHH-DMP on chromosome 12, which was 9,831 bp downstream of the closest gene, transcription factor MYB61, and overlapped a repeat in the LTR_Tyr3 superfamily (Table S3). A third example, a CHG-DMP on chromosome 1, overlaps the gene ammonium transporter 1 member 3 (AMT 1;3) and the closest repeat to the DMP was 752 bp downstream (Table S3). In Arabidopsis, AMT 1;3 contributes to ammonium uptake in plants grown under nitrogen-deficient conditions. Two of these examples involved CHH-methylation and overlapped repeats, whereas one CHG-DMP overlapped the gene body.

Whether functions documented for homologs of these genes are conserved in *Q. lobata* is not known. However, all three genes were detectably expressed in transcriptomes collected the following year in the Chico garden, providing evidence that they were transcriptionally active in that sample; none showed a significant parent-offspring correlation in methylation level in either garden. Thus, the significant methylation differences between gardens are consistent with environmentally responsive methylation, although the specific environmental factors responsible remain unknown.

In sum, CHH outliers predominated, but CG and CHG plasticity outliers also occurred within or near genes and transposable elements (TEs). It is remarkable that we found such candidates despite the fact that our RRBS data covered just 5% of genomic positions. Together, these candidates suggest that environmentally responsive methylation may affect genes involved in defense, nutrient acquisition, and other environmental-responsive pathways. Further experimental studies are needed to demonstrate whether methylation of these loci or TE affects gene expression and ultimately phenotypes that shape how trees respond to changing environments.

### DNA methylation is transmitted from parent to offspring

DNA methylation, especially in the CG context, appears to be transmitted from maternal trees to offspring for a large portion of DMPs in both garden, consistent with substantial transmission of CG methylation. The symmetry of CG sites facilitates maintenance of methylation through mitotic and meiotic cell divisions (Law & Jacobsen, 2010), but not all correlations between parent and offspring are significant for all CG-DMPs, and almost none for CHG-methylation, which has the same replication mechanism (Law & Jacobsen, 2009; Law & Jacobsen, 2010). For those positions with high correlations, natural selection could increase their frequency if they are beneficial in their local environment.

In *Q. lobata*, CG methylation at drought-associated DREB genes correlated with temperature gradients (Gugger et al., 2016). Specifically, individuals from warmer regions showed higher CG methylation at DREB genes. Such an association could reflect Darwinian selection acting on pre-existing epigenetic variation (Schmitz *et al*., 2011; Schmitz *et al*., 2013b), or environmentally induced methylation differences among individuals. In Arabidopsis, Cortijo *et al*. (2014) provided compelling evidence that epiallelic variation may contribute to the heritability of complex traits, thereby generating phenotypic variation on which selection could act. Together these findings support Prediction 2, but the number of generations over which these methylation states remain stable is unknown. Even transiently inherited methylation states could contribute to phenotypic variation, but their evolutionary significance depends on their persistence. Nonetheless, this uncertainty reflects Loison’s (2018) critique of claims about epigenetic inheritance. Moreover, Moreover, parent-offspring correlations alone do not support Lamarckian-like inheritance. To meet Prediction 3, DMPs must show both environmental induction (Prediction 1) and transmission from parent to offspring (Prediction 2).

### Lamarckian concept revisited

Testing Prediction 3 is challenging because most of the environmentally plastic DMPs are CHH methylated positions (mCHH), which are rarely inherited, whereas heritable CG methylation is rarely environmentally induced. Among the 200,000+ DMPs we examined and the 84 significant plasticity outliers, we found only 15 candidate positions that met the combined criteria for environmental plasticity and parent-offspring correlation. Two of those positions were neighboring CHH-DMP plasticity outliers associated with the zinc-finger stress-associated protein, for which methylation levels showed high parent-offspring correlations in both gardens (Table 2). The estimated narrow sense heritabilities were 48% (*P* = 0.005537) and 84% (*P* = 0.00036) in the Chico and Placerville gardens, respectively, indicating substantial resemblance in methylation level between offspring and their maternal tree. In addition, these two DMPs satisfy all three predictions and are located closed to the gene and repeat: 53-83 bp upstream both the gene and a repeat on chromosome 7. One other CHH-DMP appears to be consistent with the three predictions (Table 2), and it is associated with a gene that belongs to the AP2/ERF family of stress-responsive transcription factors involved in drought, salinity, cold, heat, and developmental plasticity (Wang *et al*., 2024). However, because the parent-offspring correlations are only marginally significant in both common gardens (P < 0.1) and the CHH-DMP is located 5,885 bp upstream of the gene and 760 bp upstream a TE, this latter case is not a strong example. Nonetheless, the discovery of two candidate CHH-DMPs that fit the predictions of the Lamarckian model is consistent with the possibility of inheritance of environmentally induced characteristics.

Very few studies describe cases of inheritance of acquired characteristics. Among the most convincing is a study by Song *et al*. (2025), who showed that experimental cold treatment induced hypomethylation at an epiallele that can confer heritable cold tolerance in rice. This study is impressive because it identifies a beneficial trait that could facilitate the cultivation of rice at colder latitudes and demonstrates that the trait is heritable through the F2 generation. The molecular mechanism by which hypomethylation contributes to the cold tolerance phenotype remains unclear, and further work is needed to determine how the epiallele affects cold-stress responses. Nevertheless, this study provides unusually direct evidence for environmentally induced, heritable epigenetic variation.

Another example potentially consistent with environmentally acquired characteristics comes from Norway spruce, where Viejo *et al*. (2023) report that offspring clones of adults exposed to warmer temperatures showed differences in the timing of budburst. They propose that the trees have an epigenetic memory of parental temperature exposure based on the correlation between DNA methylation and phenology, but further work is needed to test relationships among methylation gradients, gene expression, and phenology, to establish a causal mechanism. More such examples are needed to determine the prevalence of Lamarckian evolution and the interactions among DNA methylation, transposable elements, and gene regulation.

### DNA methylation is associated with gene expression

Regardless of its potential role in Lamarckian evolution, our study reveals that DNA methylation was associated with gene expression, although the strength and direction of this relationship differed markedly among methylation contexts. Our genome-wide analyses of methylation profiles in each sequence context across genes provide compelling evidence that DNA methylation is associated with gene regulation. Across 42 trees growing in the Chico common garden, CG methylation was positively associated with gene expression within gene bodies, whereas CHH methylation peaks were concentrated immediately upstream of highly expressed genes. These distinct genomic distributions also varied with the presence of nearby TEs, suggesting that the regulatory roles of DNA methylation depend on both sequence context and genomic location. These findings are consistent with the growing view that DNA methylation contributes to the regulation of plant transcriptional responses.

CG-methylation exhibited the strongest association with gene bodies and TEs, consistent with patterns reported across many angiosperms (Zilberman *et al*., 2007; Niederhuth *et al*., 2016; Bewick *et al*., 2017; Xu *et al*., 2018). In apple, for example, CG-methylation of the gene body was associated with higher gene expression for water-deficit genes (Xu *et al*., 2018), while comparative analyses across flowering plants have shown that gene body methylation is one of the most evolutionarily conserved features of plant methylomes (Niederhuth *et al*., 2016; Takuno *et al*., 2016; Bewick & Schmitz, 2017). Comparing these results with our RRBS data, we found that although relatively few CG-methylated sites showed differences between common gardens, CG-methylation was highly stable among individuals, consistent with its maintenance through symmetric methylation pathways. Such stable epialleles could therefore contribute to heritable phenotypic variation upon which natural selection can act (Schmitz *et al*., 2011; Schmitz, 2014). For example, the cold-tolerance epiallele discussed above (Song *et al*., 2025) provides an example of how environmentally responsive CG and CHG methylation could potentially contribute to adaptive evolution if maintained across generations (Quadrana & Colot, 2016).

CHH methylation appears to play a fundamentally different role from CG-methylation. Rather than accumulating within gene bodies, CHH methylation was concentrated immediately upstream of highly expressed genes, where it was strongly associated with proximity to TEs and only weakly influenced by TE identity or size. These upstream regions may contain cis-regulatory elements that coordinate transcriptional responses (Ricci *et al*., 2019), suggesting that CHH methylation could influence gene expression through modification of the regulatory landscape surrounding genes rather than directly through gene-body methylation. Similar enrichment of CHH methylation around gene-flanking TEs has been reported in several plant genomes and is thought to reflect RNA-directed DNA methylation acting to suppress TE activity while limiting the spread of heterochromatin into neighboring genes (Chan *et al*., 2005; Matzke & Mosher, 2014; Sigman & Slotkin, 2016; Niederhuth & Schmitz, 2017; Zhang *et al*., 2018). This mechanism could permit continued expression of nearby genes while maintaining TE silencing and could thereby contribute to phenotypic variation in plant response to environmental conditions.

Unlike CG methylation, however, CHH methylation is asymmetric and must be continually re-established (Law & Jacobsen, 2010). Our results suggest that CHH methylation primarily contributes to environmentally responsive phenotypic plasticity, enabling individuals to adjust transcriptional responses as environmental stimuli change. For example, drought induced CHH methylation was associated with expression of genes that may reduce the effects of water limitation (Peck & Sork, 2026), and widespread phosphate starvation-induced gene expression drove transient CHH methylation (Secco *et al*., 2015). Given that CHH methylation is less likely to be stably inherited because it requires continual reestablishment (Law & Jacobsen, 2010), it may contribute more strongly to environmentally responsive phenotypic plasticity rather than to stable transgenerational inheritance. This distinction may explain why CHH methylation is often the most environmentally responsive methylation context yet comparatively rarely implicated in long-term evolutionary change.

Therefore, taken together, our results suggest that the evolutionary significance of DNA methylation depends on sequence context CG provides a source of relatively stable epigenetic variation that may contribute to long-term adaptation, whereas CHH methylation primarily mediates dynamic transcriptional responses that facilitate its environmental responsiveness.

### Limitations

Our survey of genes affected by DNA methylation was conducted using reduced representation library sequencing (RRBS), which captures about 5% of the valley oak genome. It is possible that we would have identified other candidate loci for Lamarckian-like evolution had we surveyed the entire genome. However, while this small percentage means that we may have missed possible candidates, it does not weaken the candidates reported here. A second limitation is that we assessed gene expression at a single time point, two years later than the RRBS data were collected. This separation in sampling times means that our methylation-gene expression associations would not include short-term expression responses present at the time of the DNA methylation sampling. A further limitation is that our stress-associated candidate gene might represent a beneficial response to the environmental stress, but further research is needed to determine how this gene affects phenotypic variation and whether any resulting phenotype affects tree fitness before we can conclude that the stress phenotype generated by DNA methylation is beneficial.

## Conclusions

DNA methylation is associated with gene expression and may contribute to phenotypic variation associated with plant responses to the environmental conditions. The extent to which heritable epigenetic variation is as important as genetic variation remains unknown, but epimutations may generate new opportunities for phenotypic change, even if they are stable for only a few generations. Environmentally responsive methylation may be especially important than heritable epigenetic variation for facilitating rapid responses to environmental conditions. For long-lived trees, CHH methylation may be important for adjustment to changing climatic conditions, and its role warrants future attention.

## Acknowledgments

We acknowledge the native peoples of California as the traditional caretakers (past, present, and future) of the oak ecosystems sampled for this project. We are grateful to the following people for research assistance: Dylan Burge, Claudia Henriquez, Alayna Mead, and Scott O’Donnell. We thank Jessica Wright for her significant contribution to this study and the USDA Forest Service staff who provided logistical support for the provenance study (especially A Delfino-Mix and C Canning at the Institute of Forest Genetics at Placerville, California, and L Crane at the Genetic Resource and Conservation Center at Chico, California). We thank Sorel Fitz-Gibbon for significant contribution to the bioinformatic analysis. Genome and methylome studies were supported by NSF PGRP grant (IOS-1444661) to VLS, MP, and colleagues. LB was supported by the UCLA La Kretz Center for California Conservation Science. LDP was supported by funding to VLS from The Seaver Institute and NSF PGRP (IOS-2512127). Support for graduate student researchers and maintenance of the common gardens was provided by NSF-LTEB (DEB-2232974) and the USDA Forest Service. Any use of product names is for informational purposes only and does not imply endorsement by the US Government.

## Data availability

Following publication, raw reads for the reduced representation and whole genome bisulphite sequencing data, and RNA sequencing data will be deposited in the relevant International Nucleotide Sequence Database Collaboration (INSDC) database with BioProject ID XXXXXXX and the SRA run accession XXXXX. Assembled methylation and transcriptome data analyzed in this study will be published in a publicly available Dryad digital repository (DOI XXXXX). Custom analyses and plotting scripts used in this study will be available in a Zenodo digital repository (DOI XXXXX).

## Author Contribution

Conceptualization, VLS, MP; methodology, LDP, LB, VLS; bioinformatic and data analyses, LDP; resources, VLS, and MP; writing—original draft preparation, VLS, LDP; writing—review and editing, LDP, LB, VLS, MP; funding acquisition, VLS, MP. All authors have read and agreed to the published version of the manuscript

## Supplemental Information

**Figure S1.**
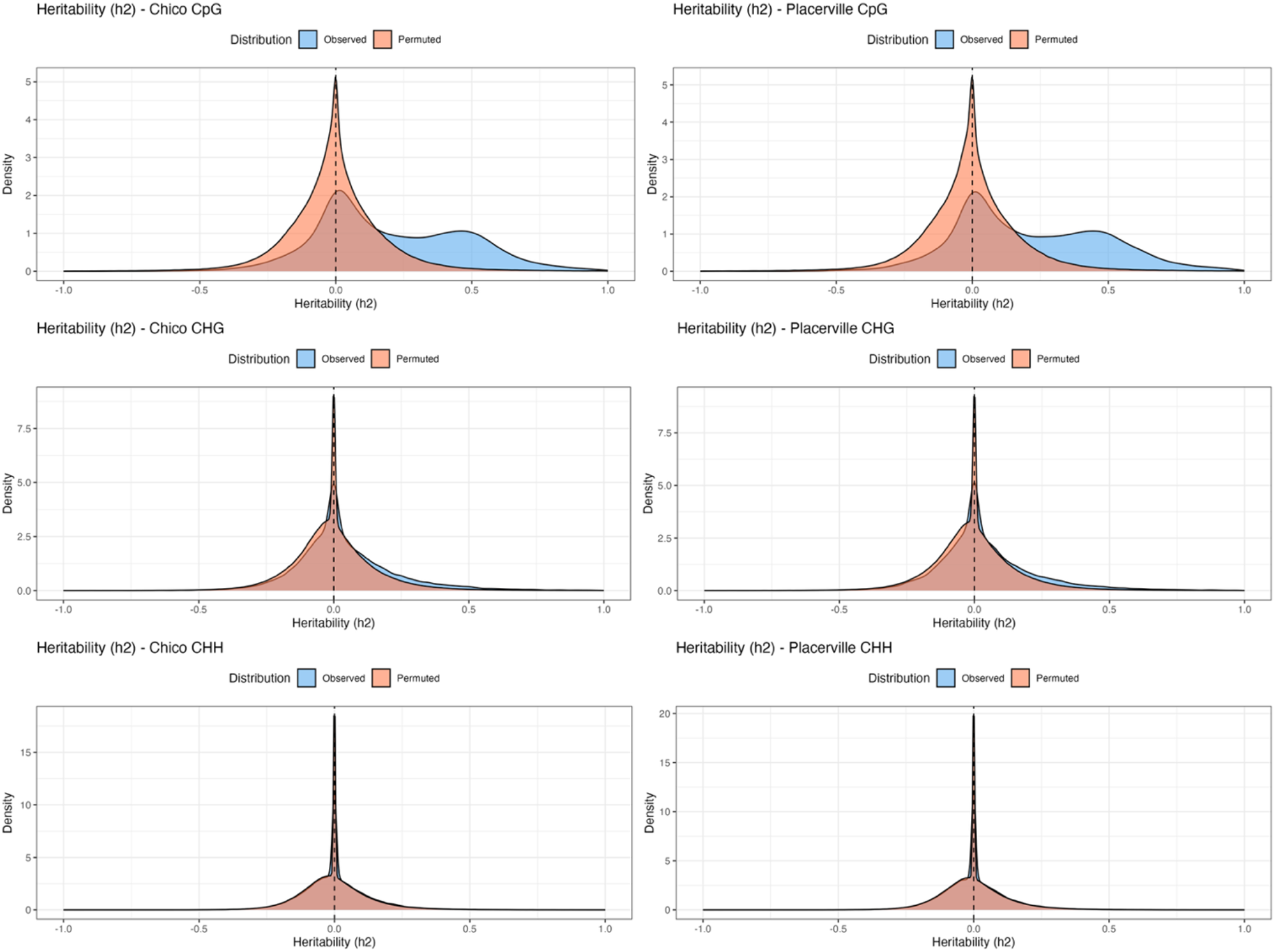
Distribution of observed (blue) and randomly permutated narrow-sense heritability for XX *Quercus lobata* trees. Heritability values (*h*^2^) for the CG-, CHG-, and CHH-DMPs (rows) growing in the two common gardens (left column: Chico Garden and right: Placerville Garden) using the slope (m) of the regression where Y = mX + b where Y=progeny methylation level and X=seed source methylation level. (See Table S4 for individual values.)

**Figure S2.**
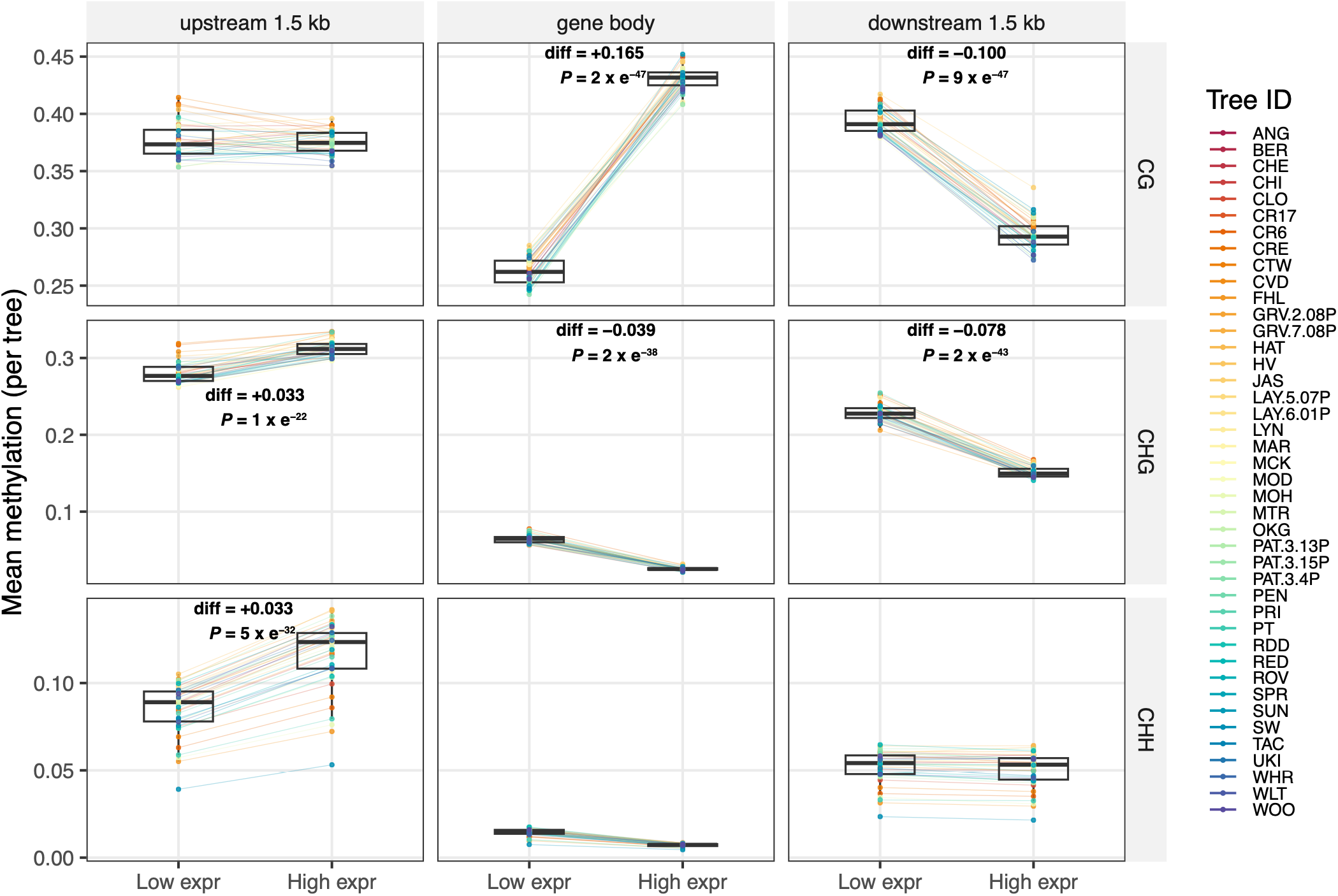
Associations between DNA methylation and gene expression across 42 *Quercus lobata* trees. Mean methylation levels were compared between the 20% most highly expressed and 20% least expressed genes within each tree for the upstream 1.5 kb flanking region, gene body and downstream 1.5 kb flanking region. Rows correspond to CG, CHG, and CHH methylation contexts. Each colored point represents one tree, with paired points connected by lines to illustrate within-tree differences between expression classes. Boxplots summarize the distribution across all trees, with lines indicating medians and boxes representing the interquartile range. Differences between expression classes were tested using paired statistical tests (df – 41), with effect sizes (median difference) and *P-*values shown for each significant panel.

**Figure S3.**
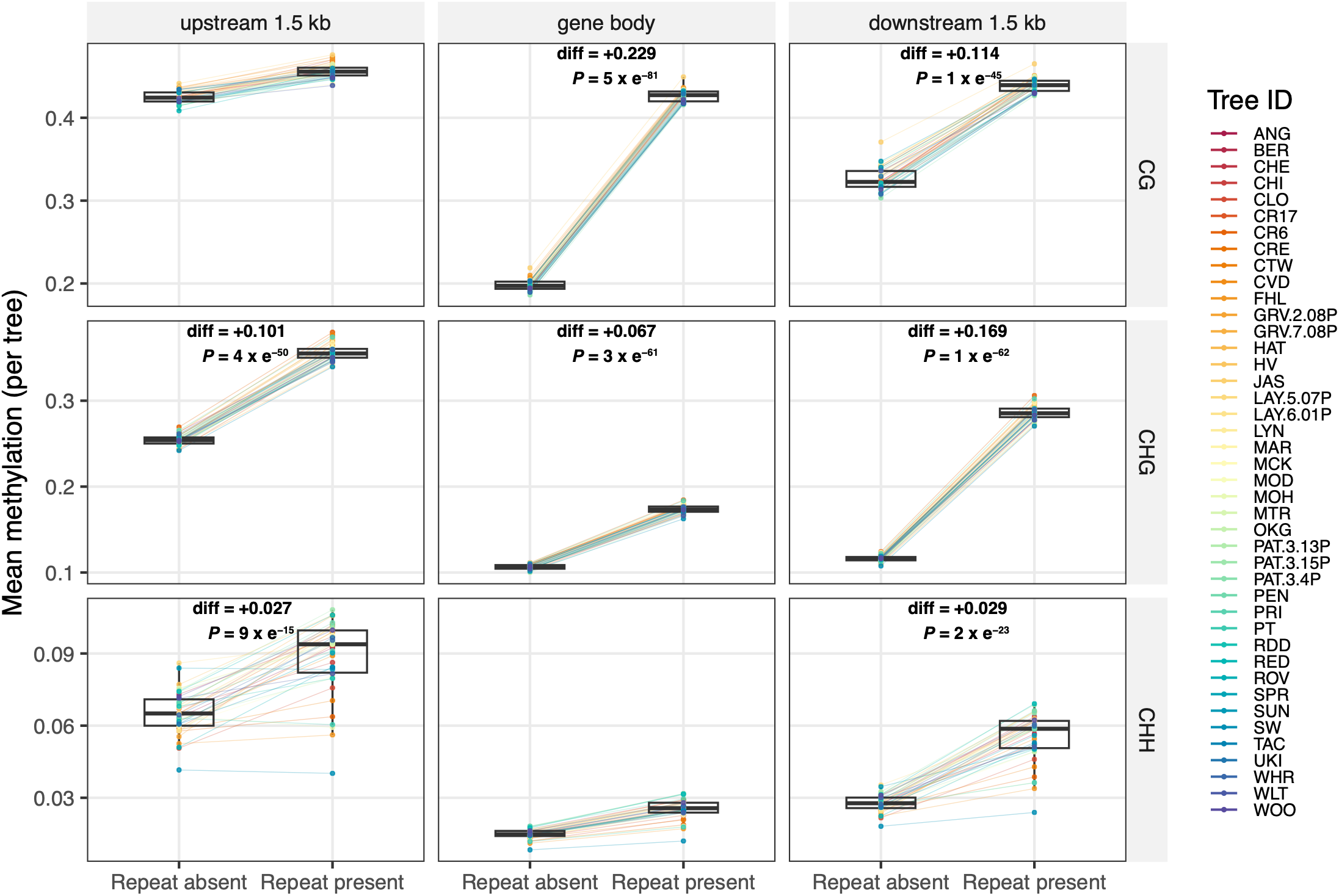
Associations between DNA methylation and repeat presence across 42 *Quercus lobata* trees. Mean methylation levels were compared between genes containing repeats and those lacking repeats. Rows correspond to CG, CHG, and CHH methylation contexts. Each colored point represents one tree, with paired points connected by lines to illustrate within-tree differences between expression classes. Boxplots summarize the distribution across all trees, with lines indicating medians and boxes representing the interquartile range. Differences between expression classes were tested using paired statistical tests (df – 41), with effect sizes (median difference) and *P-*values shown for each significant panel.

**Figure S4.**
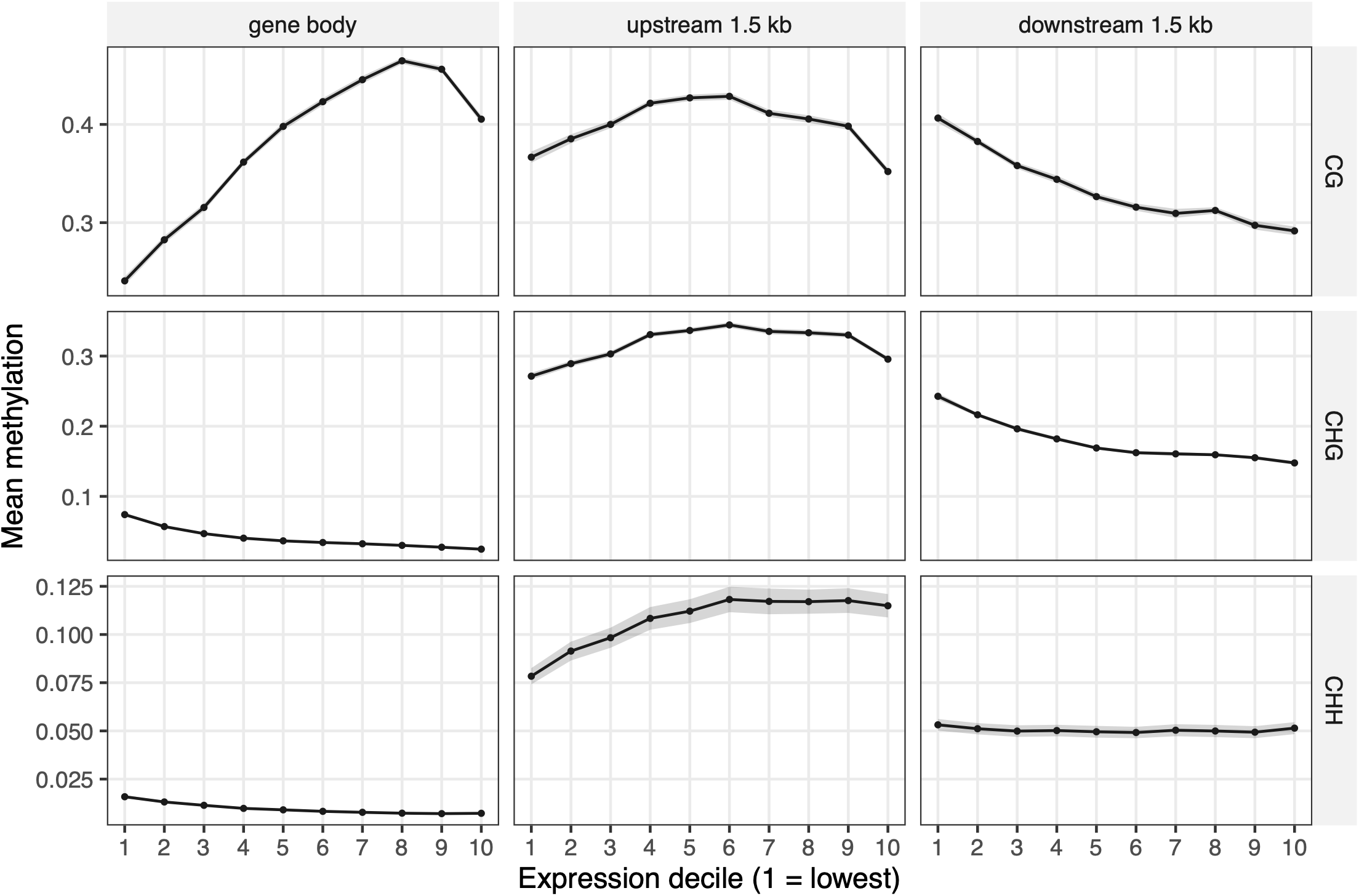
DNA methylation varies continuously with gene expression. Genes were ranked by mean expression and divided into ten equal-sized expression deciles (1 = lowest expression; 10 = highest expression). Mean methylation profiles are shown separately for CG, CHG, and CHH methylation across gene bodies and 1.5 kb upstream and downstream flanking regions. Profiles represent the mean across all 42 *Quercus lobata* trees. These data demonstrate that the relationships between DNA methylation and gene expression occurr across the full range of expression levels rather than being restricted to the highest and lowest expressed genes.

**Figure S5.**
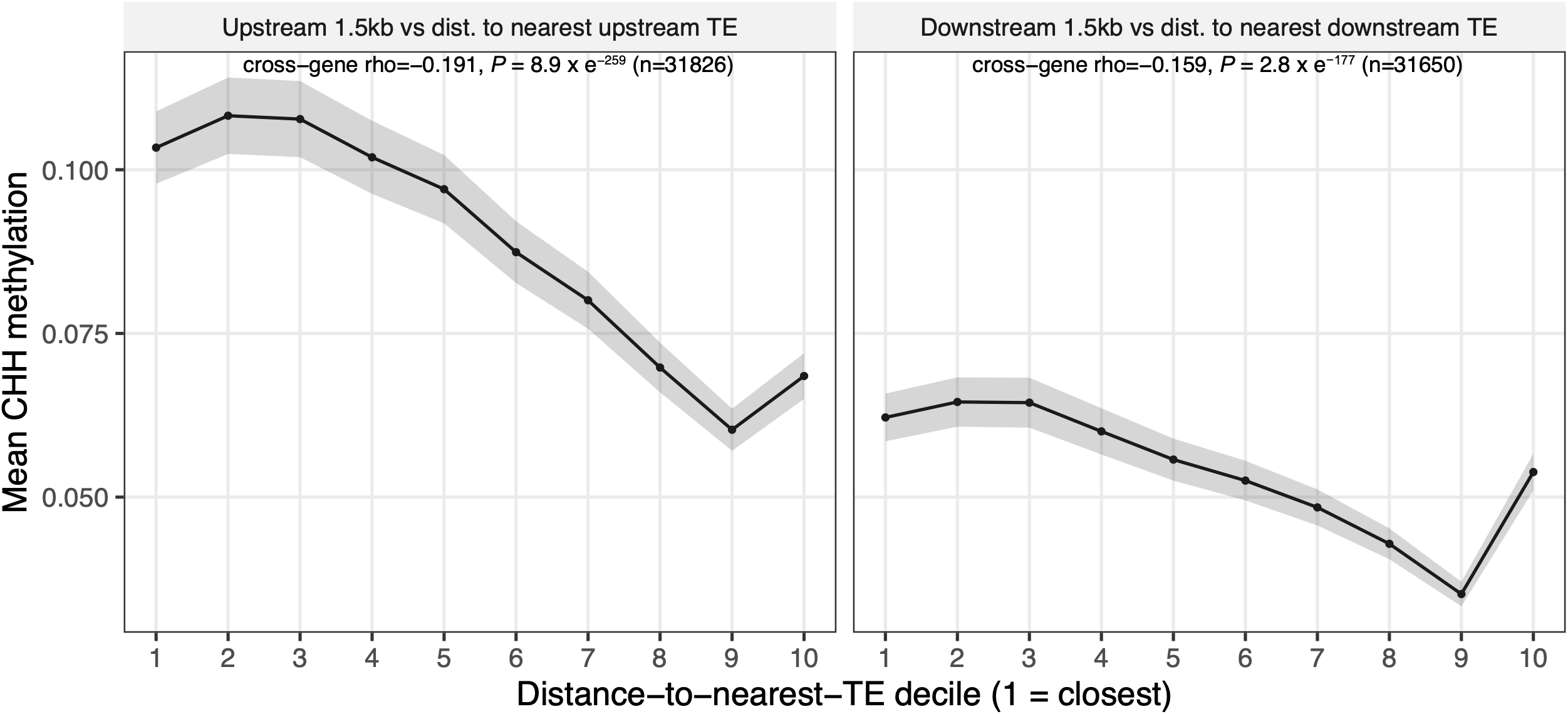
CHH methylation decreases with increasing distance from neighboring transposable elements (TEs). Mean CHH methylation profiles are shown for genes grouped according to their distance from the nearest upstream or downstream TE. Panels show upstream and downstream TE distances, respectively. Genes were binned by TE distance, and profiles represent the mean across all 42 *Quercus lobata* trees. CHH methylation is highest for genes located closest to TEs and declines progressively with increasing TE distance, demonstrating that elevated CHH methylation is concentrated around TE-proximal genes.

**Figure S6.**
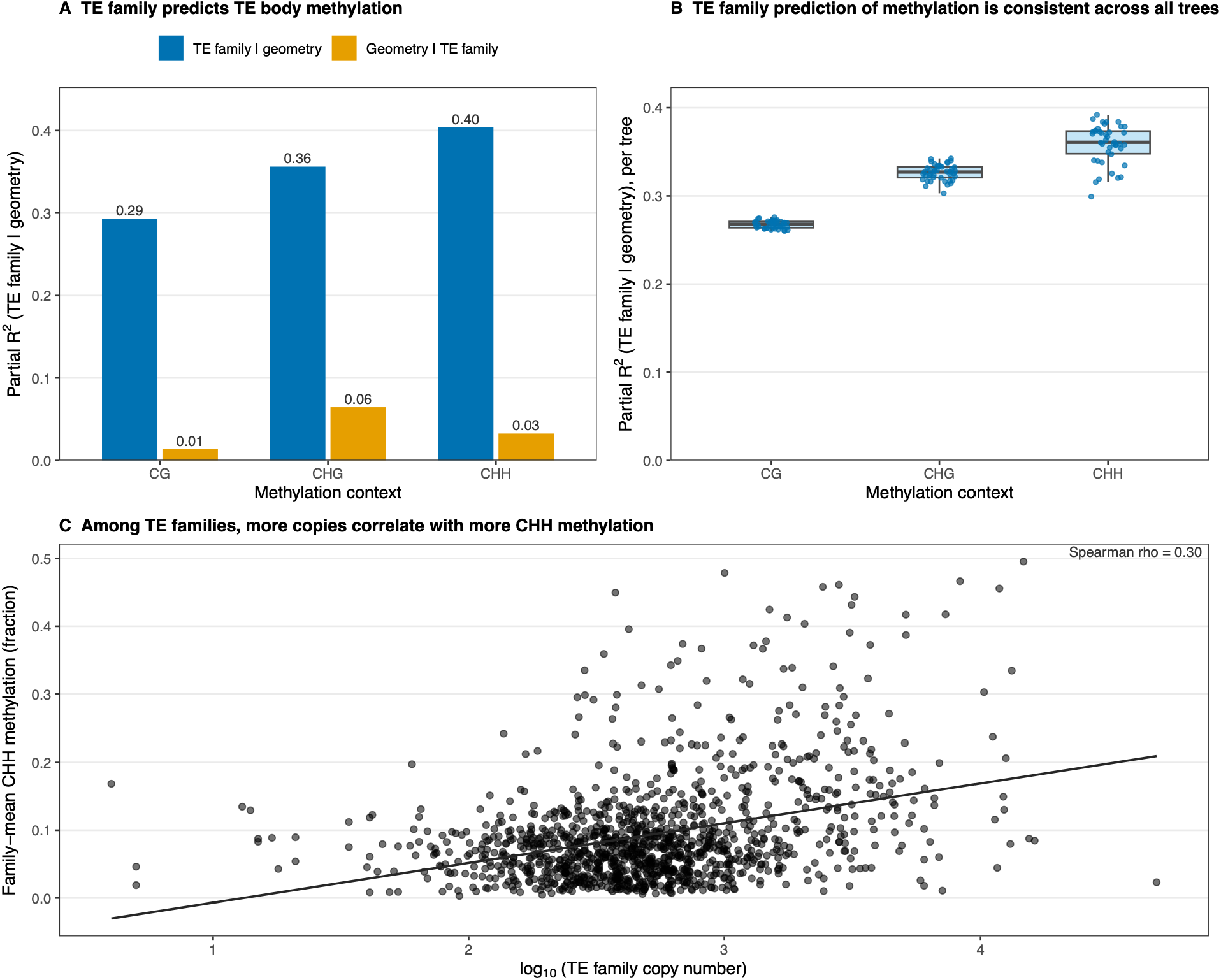
Transposable element (TE) CHH methylation is determined primarily by TE family identity rather than TE features. **A)** Variance partitioning of TE body methylation for CG, CHG and CHH contexts. Bars show the partial R² explained by TE family after accounting for TE geometry (log10 TE length, log10 distance to the nearest gene, and genic overlap; blue) and, conversely, the partial R^2^ explained by TE geometry after accounting for TE family (orange). TE family was the dominant predictor of CHH methylation (partial R^2^ = 0.40), whereas TE geometry explained only 3% of the remaining variance. **B)** Partial R^2^ of TE family after accounting for TE geometry calculated independently for each of the 42 trees, demonstrating that the strong effect of TE family is consistent among individuals. **C)** Relationship between family-mean CHH methylation and genome-wide TE family copy number for 1,227 TE families. Each point represents one TE family; the fitted line shows the ordinary least-squares regression. Families with higher copy numbers exhibited greater CHH methylation (Spearman’s ρ = 0.30), and copy number was the strongest predictor of family-level CHH methylation in the multivariate model after accounting for mean TE length, distance to the nearest gene and genic overlap (partial R^2^ = 0.13).

**Table S1.** *Quercus lobata* trees used in this study. List of sampled *Q. lobata* trees, locality details of seed source (See **Figure 1** for map of adults), and the specific analysis conducted. Maternal ID represents the family of trees growing in common gardens. Gard. Diffs. Represents the paired t-test of methylation levels of progeny in two gardens within family (Figs.2a, 3); Seas. Diff. represents the paired t-test of methylation levels within tree between seasons (Fig. 2b); Parent-offspring represents correlation and heritability tests (Figs. 4, S1). Sample sizes of total number of progenies per family used for each of the five analyses are listed within each cell.

| Locality ID | Maternal ID | Locality: Nearby town or area | latitude | Longitude | Elevation | County | Gard. diffs. Fig 2,3 | Seas. diffs. Fig 2 | Parent-offspring Fig 4, S1 | Methy -lome Fig 5, 6 | Transcrip -tome Fig 5, 6 |
| --- | --- | --- | --- | --- | --- | --- | --- | --- | --- | --- | --- |
| ANG | ANG.5 | Angel's Camp | 38.051 | -120.623 | 448 | Tuolumne | 4 |  | 4 | 1 | 1 |
| BER | BER.1 | Meadow Vista | 38.9966 | -121.1084 | 402 | Placer | 4 |  | 4 | 1 | 1 |
| CHE | CHE.1 | Chesebro, Agoura | 34.1636 | -118.7241 | 327 | Los Angeles | 4 | 1 | 4 | 1 | 1 |
| CHI | CHI.3b | Chico | 39.7119 | -121.7842 | 79 | Placer | 4 |  | 4 | 1 | 1 |
| CLO | CLO.4 | Clear Lakes Oaks | 39.0219 | -122.7135 | 401 | Colusa | 4 |  | 4 | 1 | 1 |
| CR | CR.17 | Tehachapi Mtns | 34.9879 | -118.6199 | 1766 | Los Angeles | 4 | 1 | 4 | 1 | 1 |
| CR | CR.6 | Tehachapi Mtns | 35.0069 | -118.7076 | 590 | Kern | 4 |  | 4 | 1 | 1 |
| CRE | CRE.5 | Creston | 35.5349 | -120.524 | 335 | San Luis Obispo | 2 | 1 | 2 | 1 | 1 |
| CTW | CTW.6 | Cottonwood | 40.3602 | -122.3067 | 134 | Shasta | 4 | 2 | 4 | 1 | 1 |
| CVD | CVD.8 | Russian River | 38.8544 | -123.0319 | 131 | Mendocino | 4 | 1 | 4 | 1 | 1 |
| FHL | FHL.5 | Fort Hunter Leggett | 35.9804 | -121.2328 | 300 | Monterey | 4 | 1 | 4 | 1 | 1 |
| GRV | GRV.2.08P | Gravelly Valley | 39.4302 | -122.9754 | 550 | Colusa | 4 |  | 4 | 1 | 1 |
| GRV | GRV.7.08P | Gravelly Valley | 39.4485 | -122.964 | 557 | Colusa | 4 |  | 4 | 1 | 1 |
| HAT | HAT.4 | Hills Ferry | 37.3517 | -120.9617 | 18 | Merced | 4 | 1 | 4 | 1 | 1 |
| HV | HV.1 | Hidden Valley | 34.1431 | -118.8999 | 305 | Los Angeles | 4 |  | 4 | 1 | 1 |
| JAS | JAS.5 | Jasper Ridge Reserve | 37.4032 | -122.2436 | 123 | San Mateo | 4 | 2 | 4 | 1 | 1 |
| LAY | LAY.5.07zp | Laytonville | 39.746 | -123.5242 | 457 | Mendocino | 4 |  | 4 | 1 | 1 |
| LAY | LAY.6.01P | Laytonville | 39.6722 | -123.4807 | 500 | Mendocino | 4 |  | 4 | 1 | 1 |
| LYN | LYN.4 | Arroyo Grande | 35.7878 | -120.9391 | 282 | San Luis Obispo | 4 |  | 4 | 1 | 1 |
| MAR | MAR.B | Bootjack | 37.4611 | -119.8797 | 672 | Mariposa | 4 | 1 | 4 | 1 | 1 |
| MCK | MCK.5 | Middle Creek | 39.2524 | -122.9516 | 443 | Colusa | 4 |  | 4 | 1 | 1 |
| MOD | MOD.8 | St. Helena | 38.5145 | -122.0775 | 55 | Napa | 4 | 1 | 4 | 1 | 1 |
| MOH | MOH.3 | Coyote | 37.1649 | -121.7148 | 95 | Santa Clara | 4 |  | 4 | 1 | 1 |
| MTR | MTR.3 | Pine Grove | 38.275 | -120.5058 | 730 | Amador | 4 |  | 4 | 1 | 1 |
| OKG | OKG.3 | Stockton | 38.0566 | -121.3643 | -6 | Sacramento | 4 |  | 4 | 1 | 1 |
| PAT | PAT.3 | Oakdale | 37.4959 | -121.0663 | 26 | Stanislaus | 4 |  | 4 | 3 | 3 |
| PEN | PEN.5 | Penn Valley | 39.2034 | -121.1902 | 422 | Tahama | 4 | 2 | 4 | 1 | 1 |
| PRI | PRI.3 | Princeton | 39.4555 | -122.0172 | 39 | Glen | 4 |  | 4 | 1 | 1 |
| SLC | PT.5 | San Lorenzo Creek | 36.1389 | -120.7506 | 446 | Monterey | 4 |  | 4 | 1 | 1 |
| RDD | RDD.7 | Redding | 40.611 | -122.3462 | 176 | Tahama | 4 |  | 4 | 1 | 1 |
| RED | RED.1 | Red Bluff | 40.1478 | -122.1382 | 66 | Butte | 4 | 2 | 4 | 1 | 1 |
| ROV | ROV.3 | Round Valley | 39.7483 | -123.2484 | 438 | Mendocino | 4 |  | 4 | 1 | 1 |
| SPR | SPR.h2 | Springville | 36.0919 | -118.8148 | 267 | Inyo | 4 | 1 | 4 | 1 | 1 |
| SUN | SUN.5 | Sunol | 37.5987 | -121.8751 | 82 | Santa Clara | 4 | 1 | 4 | 1 | 1 |
| SW | SW.786 | Sedgwick Reserve | 34.6882 | -120.0362 | 323 | Santa Barbara | 4 | 1 | 4 | 1 | 1 |
| TAC | TAC.8 | Tehachapi Mtns | 35.1139 | -118.4893 | 1292 | Kern | 4 |  | 4 | 1 | 1 |
| UKI | UKI.5 | Ukiah | 39.0924 | -123.2197 | 196 | Mendocino | 4 |  | 4 | 1 | 1 |
| WHR | WHR.7 | Three Rivers | 35.8365 | -118.9634 | 190 | Tulare | 4 |  | 4 | 1 | 1 |
| WLT | WLT.2 | Willits | 39.4114 | -123.3469 | 416 | Mendocino | 4 |  | 4 | 1 | 1 |
| WOO | WOO.4 | Woodson | 39.9111 | -122.0912 | 55 | Butte | 4 | 2 | 4 | 1 | 1 |
| Sample sizes |  |  |  |  |  |  | 160 | 21 | 160 | 42 | 42 |
| No. families |  |  |  |  |  |  | 40 | 16 | 40 | 40 | 40 |

**Table S2.** Summary of methylation statistics. Five datasets generated by reduced representation bisulfite sequencing and used to assess plasticity and parent-offspring transmission for testing predictions 1-3: number and percent of cytosine-containing sites; number of differentially methylated positions (DMPs); percent of DMPs per context out of total number of DMPs across all contexts. All DMPs had a minimum read coverage of 10x, maximum read coverage = mean + (5 x standard deviation), and maximum missing data across samples = 20%. Additional filters applied include: DMPs: Outliers winsorized at 10%ile and 90%ile, with a minimum range of variation = mean of standard deviation of variation at each site). We only used methylated positions with greater than minimum variation for DMPs at each context: CG=0.045, CHG=0.024, CHH=0.011.

| Methylation context | # Positions | % of Total Positions | # DMPs | % DMPs of Total DMPs | % DMPs of # Positions |
| --- | --- | --- | --- | --- | --- |
| **A. 40 adults, species wide, Fall 2012** |  |  |  |  |  |
| CG | 59,447 | 14% | 19,477 | 8% | 33% |
| CHG | 65,446 | 15% | 27,923 | 11% | 43% |
| CHH | 305,432 | 71% | 211,616 | 82% | 69% |
| Total | 430,325 | 100% | 259,016 | 100% | 60% |
| **B. 80 five-year-old trees (2 per 40 families per garden): Chico Garden, Spring 2017** |  |  |  |  |  |
| CG | 52,846 | 15% | 14,568 | 7% | 28% |
| CHG | 55,019 | 16% | 26,301 | 12% | 48% |
| CHH | 233,751 | 68% | 171,669 | 81% | 73% |
| Total | 341,616 | 100% | 212,538 | 100% | 62% |
| **C. 80 five-year-old trees (2 per 40 families per garden): Placerville Garden, Spring 2017** |  |  |  |  |  |
| CG | 54,387 | 15% | 15,238 | 7% | 28% |
| CHG | 56,654 | 16% | 27,355 | 12% | 48% |
| CHH | 241,523 | 69% | 176,692 | 81% | 73% |
| Total | 352,564 | 100% | 219,285 | 100% | 62% |
| **D. 21 4-year-old trees (16 families): Chico Garden, Fall 2016** |  |  |  |  |  |
| CG | 52,858 | 15% | 16,471 | 7% | 31% |
| CHG | 55,983 | 16% | 27,358 | 12% | 49% |
| CHH | 247,081 | 69% | 178,574 | 80% | 72% |
| Total | 355,922 | 100% | 222,403 | 100% | 62% |
| **E. 21 5-year-old trees (16 families): Chico Garden, Spring 2017** |  |  |  |  |  |
| CG | 49,950 | 16% | 14,268 | 8% | 29% |
| CHG | 51,645 | 16% | 20,860 | 12% | 40% |
| CHH | 217,837 | 68% | 133,105 | 79% | 61% |
| Total | 319,432 | 100% | 168,233 | 100% | 53% |

**Table S3.**
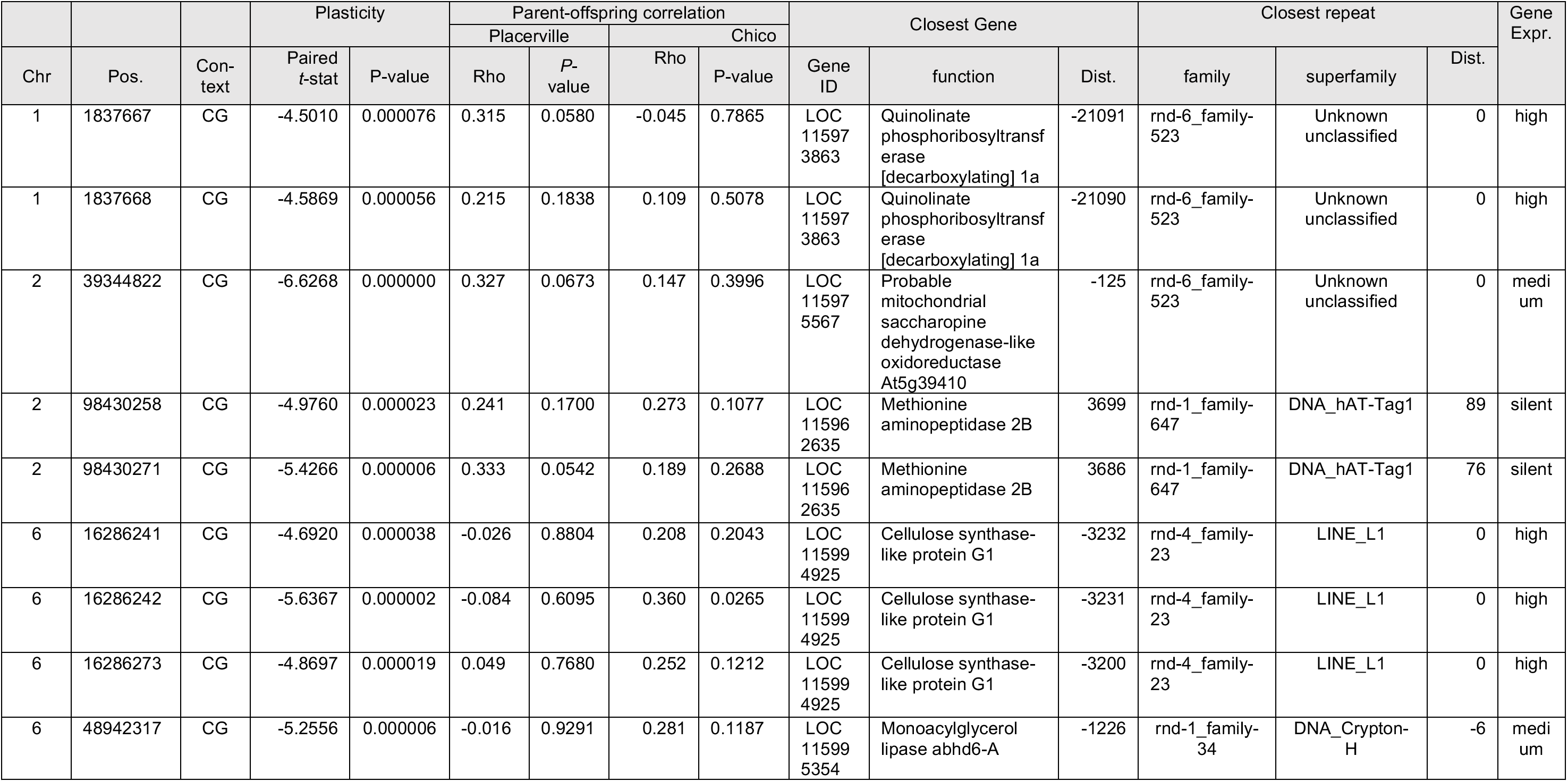
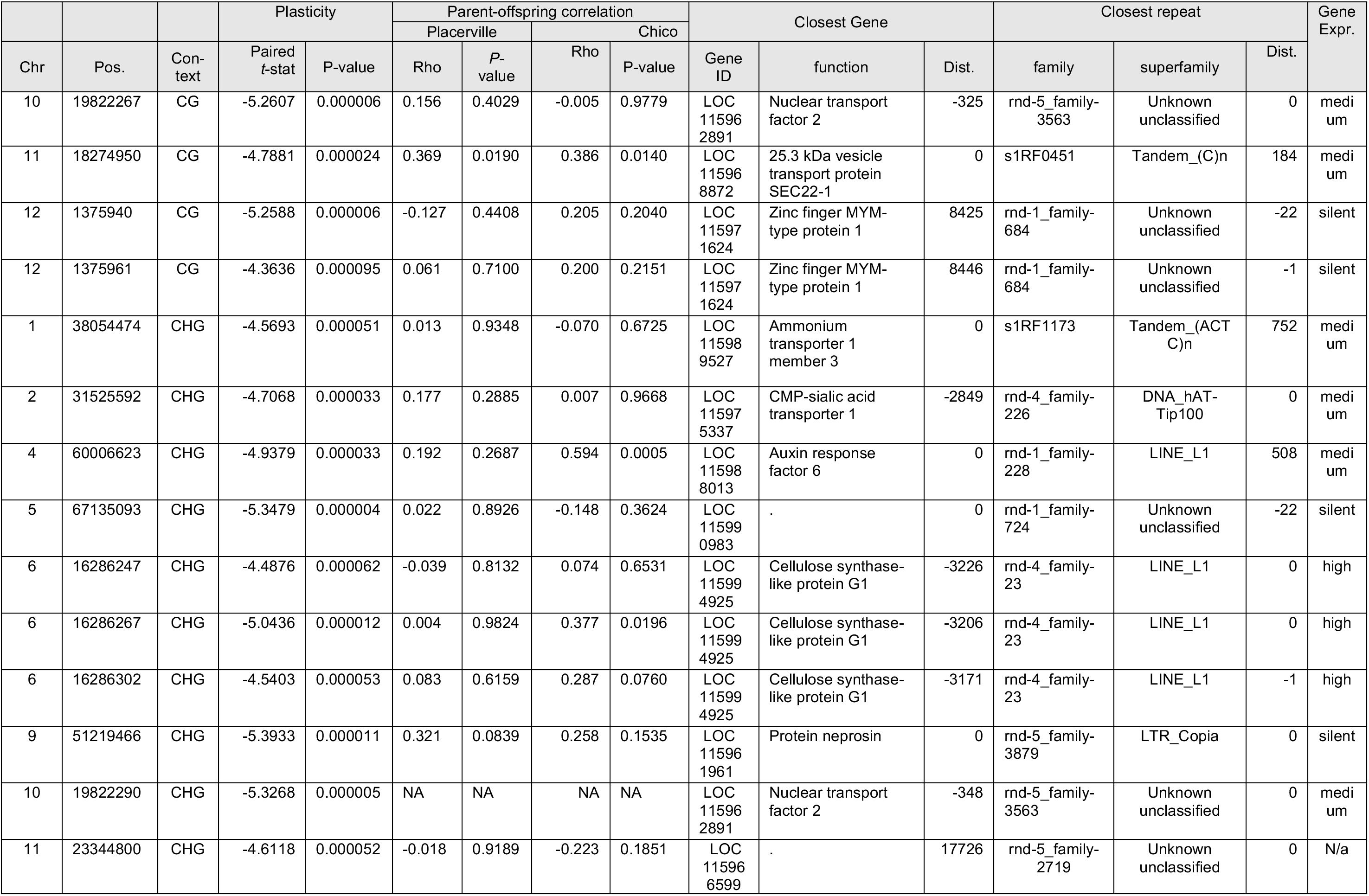
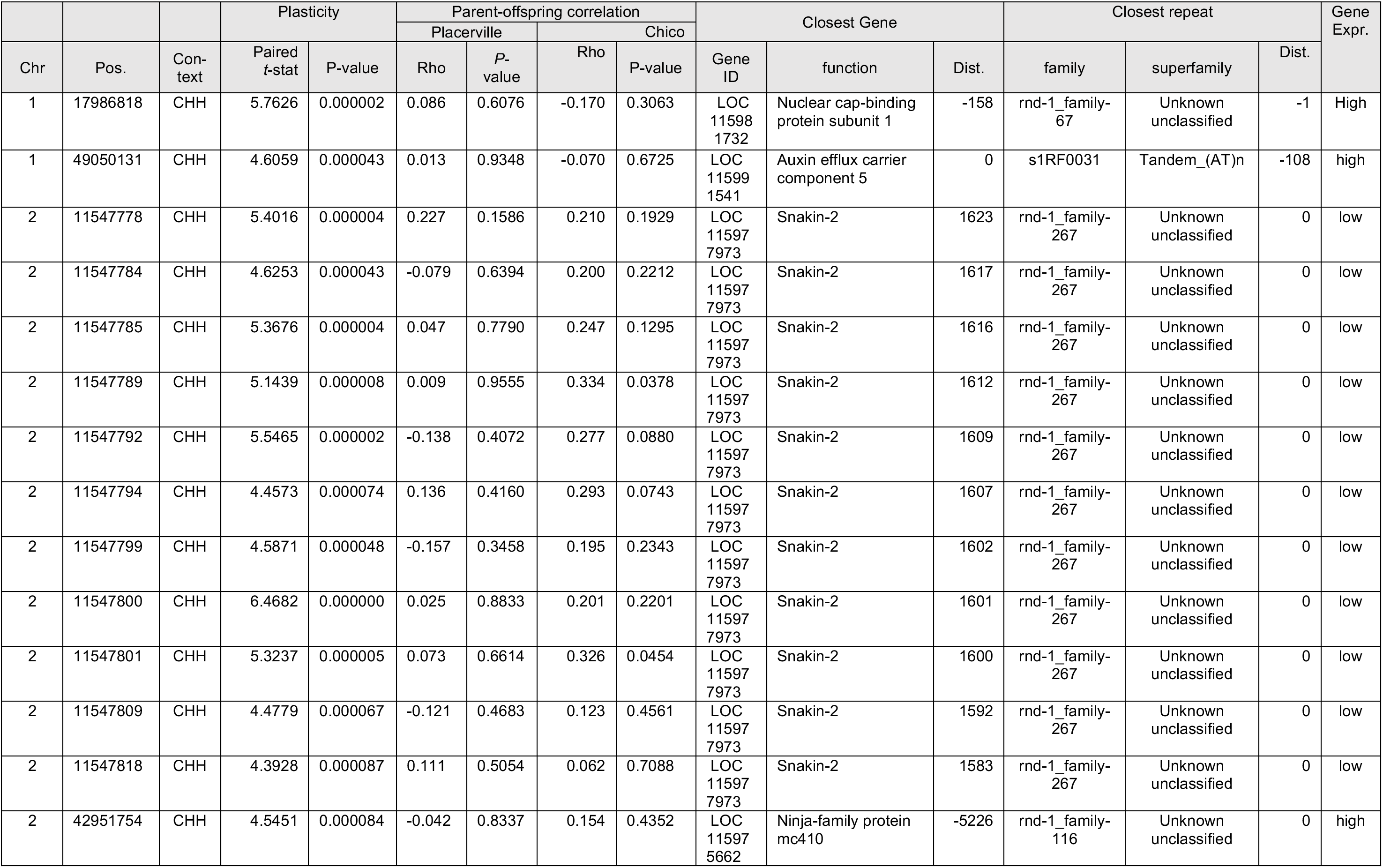
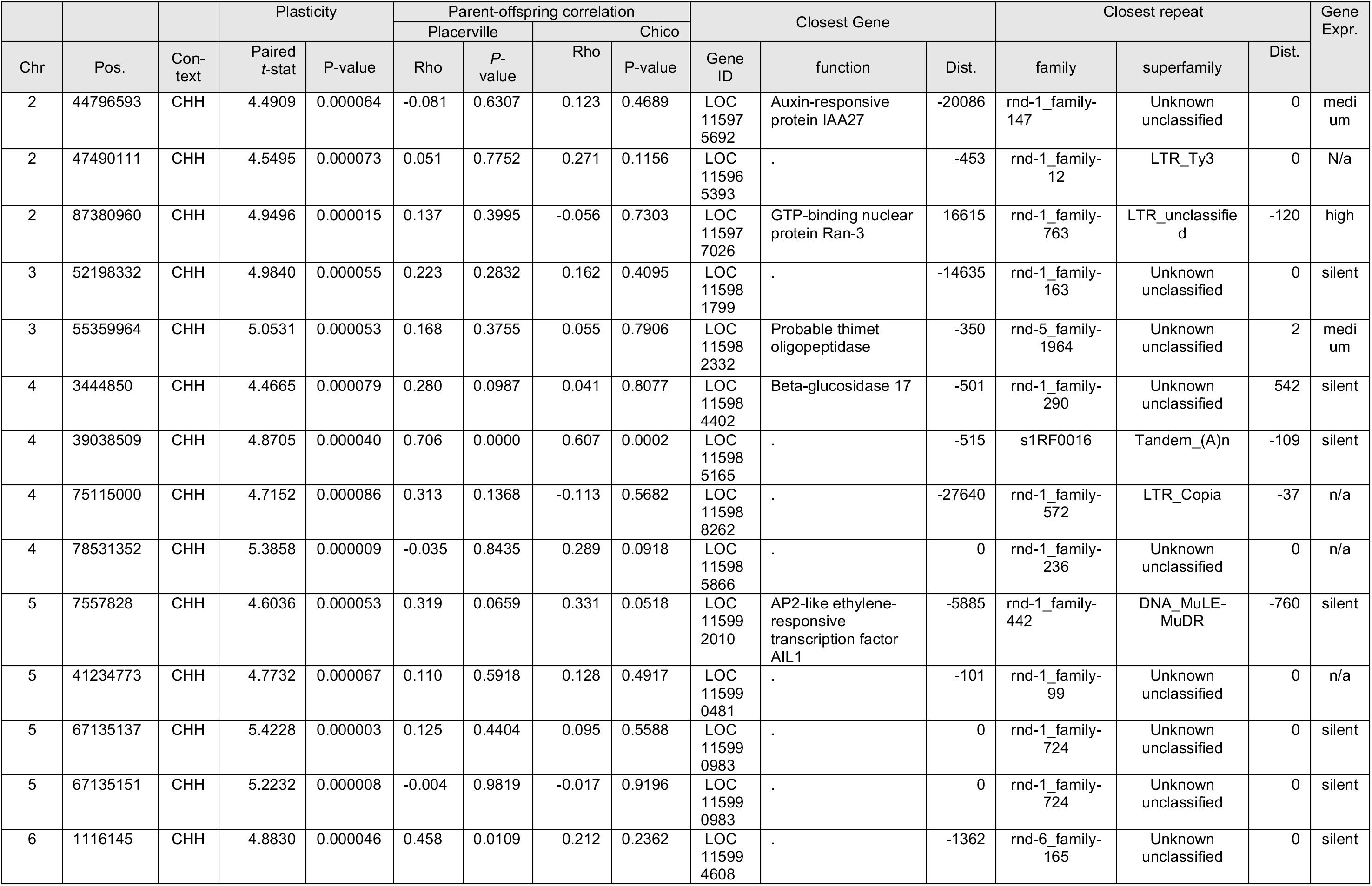
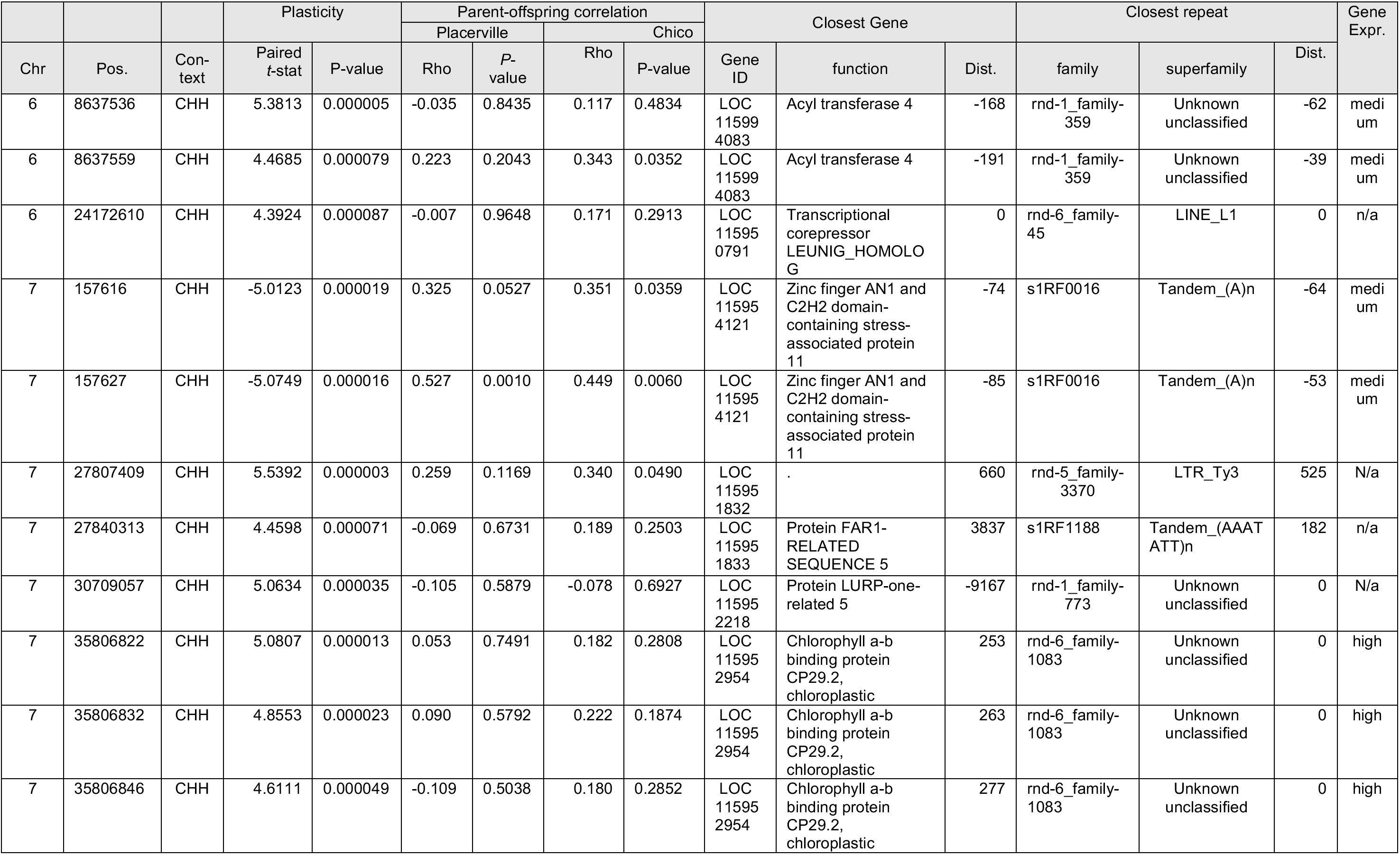
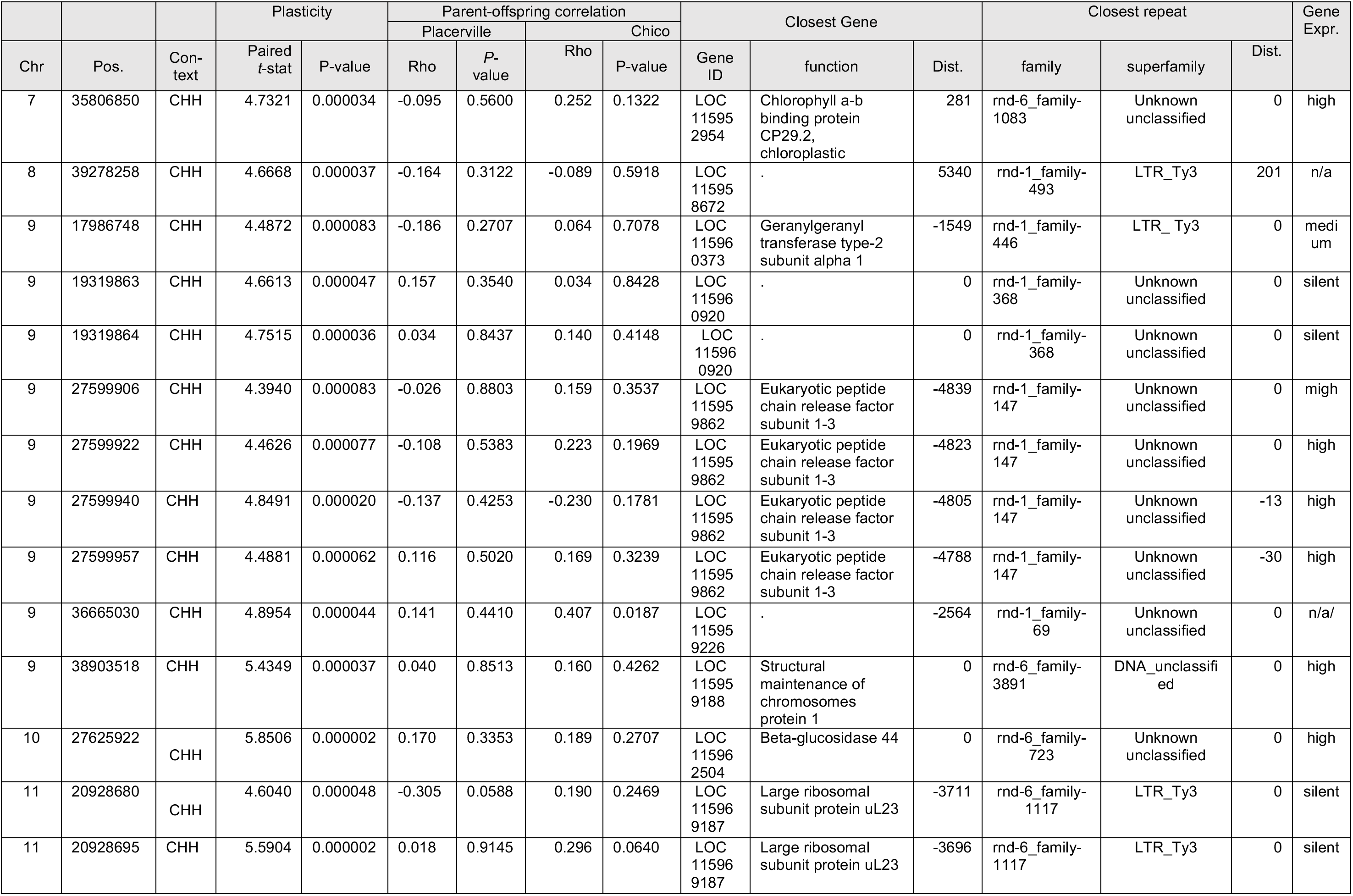
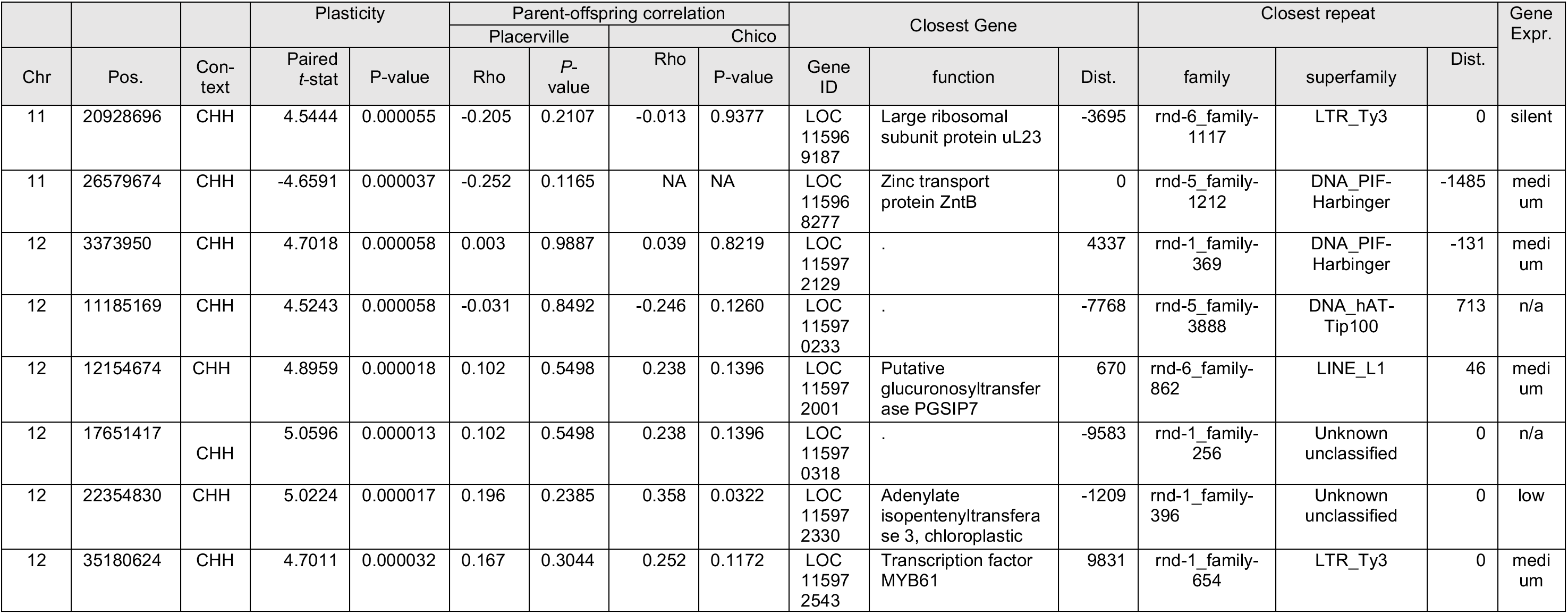
List of 84 DMP plasticity outliers include 13 out of 12,266 CG-DMPs, 9 out of 17,461 CHG-DMPs, and 62 out of 145,913 CHH-DMPs generated from Reduced Representation Library Sequencing. The plasticity outliers are defined as comprised of DMPs of 40 families differing significantly between the two Gardens based on paired t-tests with P < 0.0001. The parent-offspring correlations are based on Spearman’s rank correlation and P-values for each garden. We grouped genes into four expression categories from RNAseq data of trees growing in the Placerville Garden. Using mean depth-normalized (log2) expression for each gene, we classed the top 20% as High and the bottom 20% as Low, with Moderate covering all genes in between. Silent denotes genes that did not pass the expression filter and were excluded. Note that Ty3 TEs were formally known as Gypsy (see Wei et al., 2022).

**Table S4.** Summary of outliers of parent-offspring Spearman Rho correlations (df =39, P < 1 x 10^-5^) between levels of methylation by context in field seed source adult and mean value of two progeny for 40 families growing in two common gardens. Analyses were done separately by garden because estimates are specific to the environment where they are measured.

| Chico Garden |  |  |  |  |  | Placerville Garden |  |  |  |  |  |
| --- | --- | --- | --- | --- | --- | --- | --- | --- | --- | --- | --- |
| Chromo-<br>some | position | context | Spearman<br>Rho | P-value | $h^2$ | Chromo-<br>some | position | context | Spearman<br>Rho | P-value | $h^2$ |
| 1 | 775,681 | CG | 0.7095 | 2.97E-07 | 0.652 | 1 | 2,287,623 | CG | 0.6466 | 8.70E-06 | 0.525 |
| 1 | 2,287,616 | CG | 0.6832 | 6.11E-06 | 0.527 | 1 | 3,014,543 | CG | 0.7738 | 8.51E-07 | 0.556 |
| 1 | 4,963,699 | CG | 0.7177 | 3.94E-07 | 0.540 | 1 | 3,014,696 | CG | 0.6506 | 5.50E-06 | 0.467 |
| 1 | 7,125,371 | CG | 0.7660 | 8.42E-09 | 0.585 | 1 | 3,014,732 | CG | 0.6492 | 5.84E-06 | 0.447 |
| 1 | 11,062,737 | CG | 0.6676 | 4.69E-06 | 0.508 | 1 | 3,014,782 | CG | 0.6453 | 6.95E-06 | 0.378 |
| 1 | 13,559,895 | CG | 0.7128 | 1.55E-06 | 0.708 | 1 | 8,002,344 | CG | 0.6746 | 1.81E-06 | 0.564 |
| 1 | 28,077,888 | CG | 0.6925 | 7.41E-07 | 0.629 | 1 | 12,413,047 | CG | 0.6690 | 3.22E-06 | 0.665 |
| 1 | 29,261,169 | CG | 0.6385 | 9.27E-06 | 0.715 | 1 | 12,413,077 | CG | 0.6524 | 6.78E-06 | 0.686 |
| 1 | 30,423,848 | CG | 0.7119 | 3.74E-07 | 0.563 | 1 | 12,536,961 | CG | 0.6413 | 8.26E-06 | 0.504 |
| 1 | 31,378,233 | CG | 0.6686 | 2.42E-06 | 0.669 | 1 | 13,559,871 | CG | 0.7126 | 1.57E-06 | 0.519 |
| 1 | 38,569,979 | CG | 0.6380 | 9.49E-06 | 0.711 | 1 | 13,559,876 | CG | 0.7252 | 8.31E-07 | 0.502 |
| 1 | 39,725,092 | CG | 0.6758 | 4.44E-06 | 0.599 | 1 | 16,361,581 | CG | 0.6710 | 2.16E-06 | 0.609 |
| 2 | 7,977,491 | CG | 0.7338 | 3.54E-07 | 0.602 | 1 | 20,119,664 | CG | 0.7751 | 4.60E-08 | 0.610 |
| 2 | 8,840,427 | CG | 0.6502 | 5.61E-06 | 0.513 | 1 | 21,269,693 | CG | 0.7333 | 7.36E-08 | 0.689 |
| 2 | 12,063,923 | CG | 0.6810 | 2.53E-06 | 0.420 | 1 | 21,269,696 | CG | 0.6774 | 1.58E-06 | 0.663 |
| 2 | 12,063,924 | CG | 0.7151 | 1.39E-06 | 0.351 | 1 | 29,261,169 | CG | 0.6487 | 5.98E-06 | 0.656 |
| 2 | 12,063,960 | CG | 0.6788 | 7.38E-06 | 0.349 | 1 | 30,423,724 | CG | 0.6945 | 6.64E-07 | 0.748 |
| 2 | 12,609,086 | CG | 0.7194 | 5.10E-06 | 0.561 | 1 | 30,423,730 | CG | 0.7166 | 1.99E-07 | 0.752 |
| 2 | 18,716,289 | CG | 0.7872 | 1.70E-09 | 0.737 | 1 | 30,423,736 | CG | 0.6440 | 7.34E-06 | 0.662 |
| 2 | 19,437,162 | CG | 0.6605 | 3.52E-06 | 0.572 | 1 | 32,163,274 | CG | 0.6742 | 1.85E-06 | 0.698 |
| 2 | 19,651,390 | CG | 0.6891 | 9.24E-06 | 0.380 | 1 | 33,509,195 | CG | 0.6756 | 2.37E-06 | 0.422 |
| 2 | 21,772,719 | CG | 0.7506 | 2.07E-07 | 0.518 | 1 | 34,774,555 | CG | 0.8361 | 1.04E-07 | 0.677 |
| 2 | 22,503,872 | CG | 0.6806 | 1.35E-06 | 0.522 | 1 | 34,774,559 | CG | 0.8379 | 9.24E-08 | 0.614 |
| 2 | 22,503,966 | CG | 0.6549 | 6.09E-06 | 0.513 | 1 | 37,689,052 | CG | 0.7126 | 2.49E-07 | 0.660 |
| 2 | 24,253,425 | CG | 0.7344 | 6.88E-08 | 0.569 | 1 | 45,470,991 | CG | 0.7455 | 2.78E-07 | 0.716 |
| 2 | 24,420,266 | CG | 0.7136 | 2.35E-07 | 0.500 | 1 | 46,646,113 | CG | 0.6620 | 4.44E-06 | 0.604 |
| Chromosome | position | context | Spearman Rho | P-value | $h^2$ | Chromosome | position | context | Spearman Rho | P-value | $h^2$ |
| 2 | 32,076,074 | CG | 0.7761 | 6.91E-08 | 0.666 | 1 | 47,296,469 | CG | 0.8118 | 9.89E-09 | 0.893 |
| 2 | 32,076,087 | CG | 0.7624 | 1.61E-07 | 0.645 | 1 | 55,100,611 | CG | 0.6725 | 9.66E-06 | 0.520 |
| 2 | 32,923,988 | CG | 0.8602 | 3.53E-08 | 0.551 | 2 | 100,830 | CG | 0.6467 | 6.52E-06 | 0.868 |
| 2 | 38,816,785 | CG | 0.7984 | 1.73E-06 | 0.500 | 2 | 1,629,719 | CG | 0.6536 | 4.81E-06 | 0.588 |
| 2 | 46,503,895 | CG | 0.6465 | 6.58E-06 | 0.550 | 2 | 1,629,744 | CG | 0.7021 | 4.44E-07 | 0.641 |
| 2 | 47,490,081 | CG | 0.7059 | 9.15E-06 | 0.695 | 2 | 1,683,033 | CG | 0.6659 | 3.72E-06 | 0.537 |
| 2 | 52,591,427 | CG | 0.6952 | 6.43E-07 | 0.572 | 2 | 4,458,329 | CG | 0.6503 | 7.44E-06 | 0.265 |
| 2 | 54,330,472 | CG | 0.7529 | 3.79E-06 | 1.097 | 2 | 7,198,066 | CG | 0.6942 | 5.28E-06 | 0.569 |
| 2 | 59,430,027 | CG | 0.7004 | 6.91E-07 | 0.548 | 2 | 7,198,077 | CG | 0.7210 | 1.51E-06 | 0.566 |
| 2 | 64,042,516 | CG | 0.6627 | 4.31E-06 | 0.578 | 2 | 8,787,134 | CG | 0.7185 | 3.78E-07 | 0.785 |
| 2 | 69,679,495 | CG | 0.7921 | 8.55E-07 | 0.368 | 2 | 8,979,643 | CG | 0.6959 | 9.78E-06 | 0.483 |
| 2 | 72,627,449 | CG | 0.6764 | 5.94E-06 | 0.561 | 2 | 10,724,168 | CG | 0.6471 | 6.40E-06 | 0.464 |
| 2 | 72,983,480 | CG | 0.7229 | 1.38E-07 | 0.764 | 2 | 12,063,923 | CG | 0.6719 | 9.89E-06 | 0.318 |
| 2 | 72,983,483 | CG | 0.6475 | 6.31E-06 | 0.611 | 2 | 12,063,924 | CG | 0.7576 | 8.84E-08 | 0.427 |
| 2 | 77,499,679 | CG | 0.6857 | 1.46E-06 | 0.513 | 2 | 12,063,960 | CG | 0.7568 | 9.29E-08 | 0.427 |
| 2 | 77,499,697 | CG | 0.6725 | 2.74E-06 | 0.704 | 2 | 14,237,417 | CG | 0.6605 | 4.76E-06 | 0.684 |
| 2 | 77,499,723 | CG | 0.6705 | 7.67E-06 | 1.098 | 2 | 20,154,966 | CG | 0.6662 | 6.76E-06 | 0.485 |
| 2 | 77,499,755 | CG | 0.7269 | 1.60E-07 | 1.383 | 2 | 22,503,872 | CG | 0.6807 | 1.34E-06 | 0.536 |
| 2 | 77,499,769 | CG | 0.7573 | 2.41E-08 | 0.777 | 2 | 24,456,384 | CG | 0.6501 | 7.48E-06 | 0.536 |
| 2 | 77,499,771 | CG | 0.7336 | 1.08E-07 | 0.810 | 2 | 25,315,990 | CG | 0.7041 | 4.00E-07 | 0.676 |
| 2 | 78,435,005 | CG | 0.6678 | 2.51E-06 | 0.562 | 2 | 41,753,757 | CG | 0.7002 | 2.83E-06 | 0.505 |
| 2 | 79,911,111 | CG | 0.6558 | 5.85E-06 | 0.637 | 2 | 48,384,995 | CG | 0.7682 | 2.89E-06 | 0.482 |
| 2 | 79,911,112 | CG | 0.6609 | 3.46E-06 | 0.627 | 2 | 52,591,427 | CG | 0.7299 | 9.07E-08 | 0.502 |
| 2 | 80,837,920 | CG | 0.7096 | 1.82E-06 | 1.342 | 2 | 59,430,017 | CG | 0.6808 | 2.56E-06 | 0.582 |
| 2 | 84,618,637 | CG | 0.7202 | 3.45E-07 | 0.505 | 2 | 59,430,027 | CG | 0.7429 | 9.27E-08 | 0.621 |
| 2 | 85,113,397 | CG | 0.6986 | 8.72E-06 | 0.544 | 2 | 61,530,937 | CG | 0.6456 | 6.85E-06 | 0.707 |
| 2 | 85,555,168 | CG | 0.7679 | 4.60E-08 | 0.660 | 2 | 63,900,794 | CG | 0.6733 | 6.81E-06 | 0.430 |
| 2 | 86,447,637 | CG | 0.6711 | 2.15E-06 | 0.921 | 2 | 67,625,832 | CG | 0.6998 | 5.04E-07 | 0.759 |
| 2 | 89,426,877 | CG | 0.6853 | 4.01E-06 | 0.635 | 2 | 69,266,361 | CG | 0.6962 | 1.21E-06 | 0.817 |
| Chromo-<br>some | position | context | Spearman<br>Rho | P-value | $h^2$ | Chromo-<br>some | position | context | Spearman<br>Rho | P-value | $h^2$ |
| 2 | 91,965,103 | CG | 0.6471 | 8.51E-06 | 0.428 | 2 | 70,428,942 | CG | 0.6606 | 3.51E-06 | 0.517 |
| 2 | 92,730,033 | CG | 0.6522 | 5.12E-06 | 1.058 | 2 | 71,438,883 | CG | 0.6495 | 5.77E-06 | 0.535 |
| 2 | 92,730,037 | CG | 0.6950 | 6.47E-07 | 0.685 | 2 | 72,627,449 | CG | 0.6918 | 1.51E-06 | 0.523 |
| 2 | 93,846,367 | CG | 0.7467 | 4.82E-08 | 0.678 | 2 | 74,565,046 | CG | 0.6663 | 2.69E-06 | 0.499 |
| 2 | 93,931,520 | CG | 0.7059 | 1.06E-06 | 0.666 | 2 | 76,450,217 | CG | 0.6549 | 4.53E-06 | 0.602 |
| 2 | 93,931,525 | CG | 0.6801 | 3.65E-06 | 0.595 | 2 | 76,450,508 | CG | 0.6531 | 4.92E-06 | 0.524 |
| 2 | 97,705,344 | CG | 0.7670 | 7.82E-09 | 0.649 | 2 | 76,450,521 | CG | 0.6748 | 1.80E-06 | 0.552 |
| 2 | 97,705,358 | CG | 0.6807 | 1.34E-06 | 0.522 | 2 | 77,408,861 | CG | 0.6899 | 8.45E-07 | 0.405 |
| 2 | 100,638,222 | CG | 0.7522 | 1.23E-07 | 0.605 | 2 | 77,499,697 | CG | 0.6697 | 3.13E-06 | 0.887 |
| 3 | 3,863,316 | CG | 0.7230 | 4.35E-06 | 0.657 | 2 | 77,499,723 | CG | 0.6954 | 8.93E-07 | 1.538 |
| 3 | 12,692,506 | CG | 0.6554 | 4.44E-06 | 0.618 | 2 | 77,499,755 | CG | 0.6827 | 1.69E-06 | 0.907 |
| 3 | 12,692,808 | CG | 0.6750 | 6.32E-06 | 0.655 | 2 | 77,499,756 | CG | 0.7324 | 1.89E-06 | 1.553 |
| 3 | 18,604,044 | CG | 0.7020 | 1.83E-06 | 0.584 | 2 | 77,499,769 | CG | 0.7052 | 3.75E-07 | 1.268 |
| 3 | 32,333,109 | CG | 0.7534 | 3.70E-06 | 0.820 | 2 | 77,499,771 | CG | 0.7025 | 4.36E-07 | 1.359 |
| 3 | 33,905,174 | CG | 0.6900 | 4.51E-06 | 0.637 | 2 | 78,549,159 | CG | 0.6830 | 8.55E-06 | 0.752 |
| 3 | 38,767,288 | CG | 0.6448 | 7.10E-06 | 0.517 | 2 | 86,447,637 | CG | 0.7459 | 3.32E-08 | 0.851 |
| 3 | 43,508,674 | CG | 0.6454 | 6.92E-06 | 0.662 | 2 | 93,206,519 | CG | 0.7292 | 2.20E-06 | 0.594 |
| 3 | 43,508,677 | CG | 0.6754 | 1.74E-06 | 0.651 | 2 | 93,846,338 | CG | 0.6698 | 4.24E-06 | 0.527 |
| 3 | 46,044,491 | CG | 0.6576 | 4.02E-06 | 0.457 | 2 | 93,846,367 | CG | 0.6746 | 3.41E-06 | 0.572 |
| 3 | 49,016,337 | CG | 0.6611 | 6.24E-06 | 0.551 | 2 | 93,931,525 | CG | 0.7343 | 5.16E-07 | 0.693 |
| 3 | 49,016,400 | CG | 0.6494 | 5.81E-06 | 0.548 | 2 | 94,346,057 | CG | 0.7032 | 4.18E-07 | 0.577 |
| 3 | 50,439,234 | CG | 0.7010 | 7.87E-06 | 0.547 | 2 | 94,346,150 | CG | 0.7410 | 1.04E-07 | 0.491 |
| 3 | 54,584,064 | CG | 0.6608 | 4.68E-06 | 0.735 | 2 | 97,705,344 | CG | 0.6553 | 4.47E-06 | 0.454 |
| 3 | 54,706,699 | CG | 0.6786 | 2.05E-06 | 0.647 | 2 | 101,818,581 | CG | 0.6612 | 3.41E-06 | 0.508 |
| 3 | 58,558,297 | CG | 0.7301 | 8.99E-08 | 0.737 | 2 | 101,841,362 | CG | 0.7131 | 2.42E-07 | 0.757 |
| 3 | 65,943,024 | CG | 0.6802 | 2.63E-06 | 0.458 | 3 | 12,692,499 | CG | 0.6984 | 5.43E-07 | 0.570 |
| 3 | 67,952,830 | CG | 0.7037 | 3.43E-06 | 0.613 | 3 | 24,195,406 | CG | 0.6742 | 1.85E-06 | 0.459 |
| 3 | 68,121,571 | CG | 0.6386 | 9.23E-06 | 0.409 | 3 | 25,012,910 | CG | 0.6850 | 5.63E-06 | 0.675 |
| 3 | 69,564,275 | CG | 0.7874 | 7.39E-09 | 0.796 | 3 | 28,880,816 | CG | 0.7891 | 6.06E-07 | 0.641 |
| Chromosome | position | context | Spearman Rho | P-value | $h^2$ | Chromosome | position | context | Spearman Rho | P-value | $h^2$ |
| 3 | 69,564,276 | CG | 0.7748 | 4.68E-08 | 0.734 | 3 | 33,923,276 | CG | 0.7585 | 8.32E-08 | 0.646 |
| 3 | 69,846,517 | CG | 0.8307 | 1.50E-07 | 0.581 | 3 | 35,023,831 | CG | 0.6949 | 1.30E-06 | 0.852 |
| 4 | 4,797,045 | CG | 0.6600 | 3.60E-06 | 0.529 | 3 | 35,821,780 | CG | 0.7285 | 1.46E-07 | 0.459 |
| 4 | 10,174,723 | CG | 0.6845 | 5.77E-06 | 0.651 | 3 | 38,767,288 | CG | 0.6811 | 1.32E-06 | 0.521 |
| 4 | 10,601,410 | CG | 0.8118 | 4.51E-06 | 0.662 | 3 | 38,970,818 | CG | 0.7247 | 1.82E-07 | 0.678 |
| 4 | 17,910,973 | CG | 0.7211 | 1.54E-07 | 0.649 | 3 | 39,160,591 | CG | 0.6900 | 1.18E-06 | 0.685 |
| 4 | 30,919,409 | CG | 0.7329 | 9.19E-06 | 0.646 | 3 | 43,153,001 | CG | 0.6667 | 4.88E-06 | 0.415 |
| 4 | 32,678,546 | CG | 0.7487 | 3.53E-07 | 0.671 | 3 | 43,471,835 | CG | 0.7198 | 2.42E-07 | 0.644 |
| 4 | 33,540,050 | CG | 0.6926 | 1.45E-06 | 0.570 | 3 | 43,471,836 | CG | 0.6489 | 5.92E-06 | 0.659 |
| 4 | 41,480,372 | CG | 0.7509 | 2.40E-08 | 0.634 | 3 | 43,508,644 | CG | 0.6430 | 7.68E-06 | 0.584 |
| 4 | 41,480,373 | CG | 0.7109 | 1.18E-06 | 0.677 | 3 | 43,508,677 | CG | 0.6566 | 4.21E-06 | 0.666 |
| 4 | 41,480,379 | CG | 0.8122 | 5.50E-09 | 0.805 | 3 | 46,044,491 | CG | 0.6958 | 6.21E-07 | 0.414 |
| 4 | 41,480,384 | CG | 0.6801 | 2.64E-06 | 0.661 | 3 | 48,436,157 | CG | 0.6661 | 2.71E-06 | 0.568 |
| 4 | 43,179,413 | CG | 0.8002 | 5.45E-07 | 0.649 | 3 | 50,439,282 | CG | 0.7528 | 1.18E-07 | 0.647 |
| 4 | 48,417,218 | CG | 0.7891 | 2.95E-08 | 0.698 | 3 | 52,410,220 | CG | 0.7055 | 3.70E-07 | 0.595 |
| 4 | 53,005,283 | CG | 0.6704 | 2.22E-06 | 0.577 | 3 | 54,881,037 | CG | 0.7663 | 3.17E-07 | 0.704 |
| 4 | 53,005,285 | CG | 0.6763 | 1.67E-06 | 0.537 | 3 | 55,849,032 | CG | 0.7465 | 1.42E-06 | 0.950 |
| 4 | 53,348,901 | CG | 0.7128 | 3.57E-07 | 0.586 | 3 | 55,849,040 | CG | 0.7210 | 4.75E-06 | 0.839 |
| 4 | 53,436,340 | CG | 0.7574 | 8.92E-08 | 0.685 | 3 | 58,260,918 | CG | 0.7393 | 3.06E-06 | 0.794 |
| 4 | 58,861,597 | CG | 0.7268 | 2.47E-06 | 0.709 | 3 | 58,558,315 | CG | 0.6540 | 4.73E-06 | 0.643 |
| 4 | 59,571,510 | CG | 0.6599 | 3.61E-06 | 0.607 | 3 | 65,698,209 | CG | 0.6830 | 1.20E-06 | 0.697 |
| 4 | 59,571,881 | CG | 0.7400 | 3.78E-07 | 0.631 | 3 | 65,943,024 | CG | 0.7317 | 3.98E-07 | 0.620 |
| 4 | 67,049,220 | CG | 0.6589 | 5.10E-06 | 0.645 | 3 | 65,943,025 | CG | 0.7250 | 3.88E-07 | 0.613 |
| 4 | 69,992,625 | CG | 0.7312 | 1.25E-07 | 0.664 | 3 | 66,361,387 | CG | 0.6434 | 7.54E-06 | 0.568 |
| 4 | 70,256,921 | CG | 0.6806 | 1.35E-06 | 0.589 | 3 | 68,121,571 | CG | 0.6401 | 8.69E-06 | 0.499 |
| 4 | 75,217,680 | CG | 0.7108 | 2.76E-07 | 0.470 | 3 | 69,564,275 | CG | 0.8356 | 2.29E-10 | 0.829 |
| 4 | 77,298,883 | CG | 0.6841 | 8.17E-06 | 0.506 | 3 | 69,564,276 | CG | 0.7728 | 5.35E-08 | 0.730 |
| 5 | 9,722,824 | CG | 0.7828 | 1.19E-07 | 0.638 | 3 | 70,846,096 | CG | 0.6561 | 4.30E-06 | 0.592 |
| 5 | 9,722,908 | CG | 0.7725 | 2.22E-07 | 0.677 | 4 | 4,190,267 | CG | 0.8085 | 1.01E-06 | 0.657 |
| Chromo-<br>some | position | context | Spearman<br>Rho | P-value | $h^2$ | Chromo-<br>some | position | context | Spearman<br>Rho | P-value | $h^2$ |
| 5 | 9,722,955 | CG | 0.8122 | 2.90E-08 | 0.604 | 4 | 10,213,260 | CG | 0.7532 | 3.16E-08 | 0.624 |
| 5 | 9,722,994 | CG | 0.8044 | 4.97E-08 | 0.678 | 4 | 17,910,891 | CG | 0.6872 | 9.68E-07 | 0.521 |
| 5 | 9,722,996 | CG | 0.8055 | 4.60E-08 | 0.662 | 4 | 17,910,994 | CG | 0.6422 | 7.93E-06 | 0.589 |
| 5 | 9,723,012 | CG | 0.7099 | 7.74E-06 | 0.546 | 4 | 19,128,508 | CG | 0.6598 | 4.91E-06 | 0.884 |
| 5 | 9,723,014 | CG | 0.7222 | 4.50E-06 | 0.602 | 4 | 30,345,343 | CG | 0.6730 | 1.96E-06 | 0.618 |
| 5 | 14,169,231 | CG | 0.6808 | 1.34E-06 | 0.464 | 4 | 31,202,956 | CG | 0.7066 | 3.48E-07 | 0.671 |
| 5 | 28,778,228 | CG | 0.6465 | 6.58E-06 | 0.499 | 4 | 38,708,914 | CG | 0.7604 | 1.81E-07 | 0.587 |
| 5 | 39,731,972 | CG | 0.7190 | 7.82E-07 | 0.662 | 4 | 39,097,437 | CG | 0.6490 | 5.90E-06 | 0.578 |
| 5 | 44,180,057 | CG | 0.6634 | 3.09E-06 | 0.466 | 4 | 41,480,379 | CG | 0.7100 | 1.78E-06 | 0.594 |
| 5 | 61,024,545 | CG | 0.6757 | 4.46E-06 | 0.536 | 4 | 41,480,384 | CG | 0.6806 | 6.83E-06 | 0.536 |
| 5 | 61,024,547 | CG | 0.6618 | 8.18E-06 | 0.495 | 4 | 41,480,420 | CG | 0.7076 | 4.73E-07 | 0.486 |
| 5 | 72,806,747 | CG | 0.7094 | 6.16E-07 | 0.418 | 4 | 41,480,421 | CG | 0.7262 | 5.35E-07 | 0.522 |
| 5 | 72,806,752 | CG | 0.6752 | 2.42E-06 | 0.425 | 4 | 53,436,217 | CG | 0.6689 | 3.24E-06 | 0.585 |
| 5 | 80,563,996 | CG | 0.6866 | 1.00E-06 | 0.513 | 4 | 55,696,212 | CG | 0.6881 | 9.27E-07 | 0.681 |
| 5 | 81,019,447 | CG | 0.6757 | 1.72E-06 | 0.543 | 4 | 55,696,213 | CG | 0.7473 | 3.04E-08 | 0.641 |
| 5 | 83,924,280 | CG | 0.7095 | 1.82E-06 | 0.729 | 4 | 60,021,367 | CG | 0.6915 | 1.53E-06 | 0.608 |
| 5 | 85,095,638 | CG | 0.6841 | 1.58E-06 | 0.579 | 4 | 67,049,220 | CG | 0.6588 | 5.12E-06 | 0.503 |
| 5 | 88,737,273 | CG | 0.6608 | 3.47E-06 | 0.595 | 4 | 68,413,139 | CG | 0.6786 | 1.49E-06 | 0.618 |
| 6 | 87,595 | CG | 0.7328 | 9.25E-06 | 0.626 | 4 | 68,413,144 | CG | 0.7097 | 2.93E-07 | 0.764 |
| 6 | 3,785,434 | CG | 0.6562 | 4.29E-06 | 0.503 | 4 | 68,413,282 | CG | 0.6743 | 8.93E-06 | 0.589 |
| 6 | 12,340,426 | CG | 0.6538 | 4.76E-06 | 0.653 | 4 | 68,413,289 | CG | 0.7447 | 2.90E-07 | 0.609 |
| 6 | 12,340,432 | CG | 0.7006 | 4.82E-07 | 0.453 | 4 | 68,413,309 | CG | 0.7259 | 8.04E-07 | 0.594 |
| 6 | 14,930,807 | CG | 0.8220 | 4.45E-10 | 0.634 | 4 | 68,413,312 | CG | 0.7160 | 1.93E-06 | 0.597 |
| 6 | 22,722,744 | CG | 0.7404 | 3.69E-07 | 0.599 | 4 | 68,413,319 | CG | 0.7210 | 1.03E-06 | 0.536 |
| 6 | 26,646,054 | CG | 0.6572 | 4.09E-06 | 0.540 | 4 | 68,413,353 | CG | 0.7348 | 5.02E-07 | 0.583 |
| 6 | 26,751,196 | CG | 0.6964 | 1.20E-06 | 0.498 | 4 | 70,514,287 | CG | 0.6525 | 5.05E-06 | 0.483 |
| 6 | 26,751,203 | CG | 0.7199 | 3.51E-07 | 0.606 | 4 | 71,305,487 | CG | 0.6435 | 7.50E-06 | 0.596 |
| 6 | 27,604,399 | CG | 0.7425 | 7.52E-07 | 0.558 | 4 | 71,305,488 | CG | 0.7067 | 3.47E-07 | 0.647 |
| 6 | 29,873,999 | CG | 0.8034 | 1.82E-08 | 1.073 | 4 | 75,217,726 | CG | 0.6696 | 3.13E-06 | 0.639 |
| Chromosome | position | context | Spearman Rho | P-value | $h^2$ | Chromosome | position | context | Spearman Rho | P-value | $h^2$ |
| 6 | 29,874,002 | CG | 0.7501 | 5.01E-07 | 0.778 | 4 | 76,513,906 | CG | 0.6562 | 7.72E-06 | 0.446 |
| 6 | 33,216,149 | CG | 0.7552 | 3.80E-07 | 0.754 | 4 | 77,299,118 | CG | 0.6601 | 8.78E-06 | 0.449 |
| 6 | 33,216,160 | CG | 0.7273 | 1.63E-06 | 0.743 | 4 | 81,789,648 | CG | 0.6694 | 8.03E-06 | 0.636 |
| 6 | 33,668,119 | CG | 0.7032 | 8.50E-07 | 0.584 | 4 | 85,346,739 | CG | 0.6895 | 6.48E-06 | 0.727 |
| 6 | 43,703,782 | CG | 0.7720 | 8.66E-09 | 0.978 | 4 | 95,034,643 | CG | 0.6856 | 2.04E-06 | 0.578 |
| 6 | 43,703,783 | CG | 0.7590 | 1.38E-08 | 1.067 | 5 | 9,723,543 | CG | 0.7098 | 3.73E-06 | 0.760 |
| 6 | 43,703,806 | CG | 0.7767 | 6.17E-09 | 1.113 | 5 | 9,723,575 | CG | 0.7109 | 3.55E-06 | 0.630 |
| 6 | 43,703,807 | CG | 0.8296 | 6.68E-11 | 1.120 | 5 | 23,390,391 | CG | 0.7008 | 6.76E-07 | 0.663 |
| 6 | 44,625,142 | CG | 0.6399 | 8.74E-06 | 0.581 | 5 | 26,494,855 | CG | 0.7244 | 1.26E-07 | 0.623 |
| 6 | 44,891,925 | CG | 0.7557 | 9.91E-08 | 0.697 | 5 | 26,494,856 | CG | 0.6635 | 3.07E-06 | 0.614 |
| 6 | 44,891,939 | CG | 0.8090 | 2.37E-09 | 0.795 | 5 | 28,709,370 | CG | 0.7274 | 7.80E-06 | 0.234 |
| 6 | 44,891,945 | CG | 0.7655 | 5.38E-08 | 0.700 | 5 | 28,778,228 | CG | 0.6574 | 4.05E-06 | 0.451 |
| 6 | 44,891,947 | CG | 0.7221 | 6.64E-07 | 0.726 | 5 | 32,963,403 | CG | 0.6545 | 6.20E-06 | 0.740 |
| 6 | 44,891,951 | CG | 0.7654 | 5.42E-08 | 0.766 | 5 | 32,963,404 | CG | 0.6579 | 3.96E-06 | 0.783 |
| 6 | 44,891,969 | CG | 0.7461 | 1.76E-07 | 0.765 | 5 | 44,091,167 | CG | 0.6441 | 9.66E-06 | 0.463 |
| 6 | 44,891,975 | CG | 0.7269 | 5.15E-07 | 0.677 | 5 | 46,951,272 | CG | 0.6642 | 2.96E-06 | 0.589 |
| 6 | 44,891,978 | CG | 0.7449 | 1.89E-07 | 0.768 | 5 | 52,752,004 | CG | 0.6801 | 1.38E-06 | 0.689 |
| 6 | 47,513,336 | CG | 0.6458 | 9.00E-06 | 0.236 | 5 | 52,752,005 | CG | 0.6894 | 1.21E-06 | 0.632 |
| 6 | 48,214,399 | CG | 0.6375 | 9.67E-06 | 0.494 | 5 | 60,304,380 | CG | 0.6473 | 6.35E-06 | 0.559 |
| 6 | 52,916,296 | CG | 0.6729 | 6.92E-06 | 0.521 | 5 | 67,758,924 | CG | 0.6531 | 4.92E-06 | 0.572 |
| 7 | 779,360 | CG | 0.7995 | 4.97E-09 | 0.632 | 5 | 69,436,926 | CG | 0.7645 | 2.32E-08 | 0.677 |
| 7 | 779,375 | CG | 0.7525 | 5.08E-08 | 0.508 | 5 | 72,806,747 | CG | 0.7294 | 1.38E-07 | 0.491 |
| 7 | 779,641 | CG | 0.7137 | 2.34E-07 | 0.644 | 5 | 72,806,752 | CG | 0.6925 | 1.04E-06 | 0.479 |
| 7 | 1,533,705 | CG | 0.7216 | 6.83E-07 | 0.389 | 5 | 77,344,794 | CG | 0.6954 | 8.97E-07 | 0.596 |
| 7 | 2,603,687 | CG | 0.6518 | 5.22E-06 | 0.554 | 5 | 77,344,851 | CG | 0.7731 | 3.27E-08 | 0.485 |
| 7 | 11,022,928 | CG | 0.7486 | 3.56E-07 | 0.585 | 5 | 79,391,611 | CG | 0.6404 | 8.58E-06 | 0.478 |
| 7 | 17,653,842 | CG | 0.7299 | 1.43E-06 | 0.509 | 5 | 80,817,753 | CG | 0.6504 | 7.38E-06 | 0.443 |
| 7 | 19,546,493 | CG | 0.6834 | 1.18E-06 | 0.544 | 5 | 83,924,280 | CG | 0.6996 | 2.06E-06 | 0.569 |
| 7 | 23,521,103 | CG | 0.6872 | 9.69E-07 | 0.513 | 5 | 85,095,638 | CG | 0.6630 | 4.24E-06 | 0.483 |
| Chromosome | position | context | Spearman Rho | P-value | $h^2$ | Chromosome | position | context | Spearman Rho | P-value | $h^2$ |
| 7 | 23,521,205 | CG | 0.6881 | 9.27E-07 | 0.523 | 6 | 2,086,610 | CG | 0.6481 | 6.13E-06 | 0.589 |
| 7 | 32,729,397 | CG | 0.8020 | 4.94E-10 | 0.670 | 6 | 2,461,566 | CG | 0.7067 | 4.95E-07 | 0.892 |
| 7 | 33,969,449 | CG | 0.7078 | 1.98E-06 | 0.573 | 6 | 2,461,569 | CG | 0.6450 | 9.32E-06 | 1.574 |
| 7 | 33,969,456 | CG | 0.7054 | 2.22E-06 | 0.620 | 6 | 6,188,137 | CG | 0.6464 | 8.77E-06 | 0.575 |
| 7 | 34,253,299 | CG | 0.7369 | 1.33E-07 | 0.629 | 6 | 6,771,196 | CG | 0.7243 | 1.27E-07 | 0.488 |
| 7 | 34,266,079 | CG | 0.6397 | 8.81E-06 | 0.602 | 6 | 6,875,703 | CG | 0.7775 | 6.34E-08 | 0.835 |
| 7 | 34,266,134 | CG | 0.6786 | 1.49E-06 | 0.532 | 6 | 6,875,705 | CG | 0.7046 | 3.30E-06 | 0.787 |
| 7 | 34,345,427 | CG | 0.7103 | 4.08E-07 | 0.620 | 6 | 8,664,195 | CG | 0.6735 | 3.59E-06 | 0.644 |
| 7 | 34,345,428 | CG | 0.6830 | 1.20E-06 | 0.569 | 6 | 12,340,426 | CG | 0.7107 | 2.78E-07 | 0.715 |
| 7 | 34,345,444 | CG | 0.7310 | 8.50E-08 | 0.534 | 6 | 14,433,431 | CG | 0.7802 | 2.60E-06 | 0.507 |
| 7 | 34,386,701 | CG | 0.7757 | 1.07E-08 | 0.653 | 6 | 14,930,882 | CG | 0.6599 | 3.62E-06 | 0.560 |
| 7 | 34,888,481 | CG | 0.6640 | 3.00E-06 | 0.532 | 6 | 20,122,099 | CG | 0.7062 | 7.27E-07 | 0.606 |
| 7 | 34,888,482 | CG | 0.6448 | 7.10E-06 | 0.483 | 6 | 26,646,054 | CG | 0.6380 | 9.50E-06 | 0.569 |
| 7 | 35,942,799 | CG | 0.7614 | 2.54E-06 | 0.795 | 6 | 26,646,084 | CG | 0.6630 | 3.14E-06 | 0.531 |
| 7 | 36,252,028 | CG | 0.7326 | 1.15E-07 | 0.519 | 6 | 33,668,119 | CG | 0.7162 | 2.95E-07 | 0.597 |
| 7 | 36,935,923 | CG | 0.6947 | 2.12E-06 | 0.680 | 6 | 39,683,299 | CG | 0.7690 | 1.71E-07 | 0.554 |
| 7 | 44,634,632 | CG | 0.6456 | 6.84E-06 | 0.567 | 6 | 42,854,343 | CG | 0.7531 | 5.81E-06 | 0.596 |
| 7 | 44,808,394 | CG | 0.6787 | 1.48E-06 | 0.528 | 6 | 43,703,782 | CG | 0.6869 | 1.37E-06 | 0.973 |
| 8 | 1,535,813 | CG | 0.6575 | 4.04E-06 | 0.601 | 6 | 43,703,783 | CG | 0.7122 | 3.68E-07 | 1.017 |
| 8 | 4,007,741 | CG | 0.7124 | 2.53E-07 | 0.585 | 6 | 43,703,807 | CG | 0.6943 | 9.46E-07 | 1.045 |
| 8 | 4,224,847 | CG | 0.6990 | 2.12E-06 | 0.621 | 6 | 43,821,948 | CG | 0.7809 | 1.92E-08 | 0.696 |
| 8 | 11,669,243 | CG | 0.7609 | 1.21E-08 | 0.570 | 6 | 44,423,230 | CG | 0.6649 | 2.87E-06 | 0.435 |
| 8 | 11,783,972 | CG | 0.7245 | 5.87E-07 | 0.514 | 6 | 44,752,700 | CG | 0.7206 | 1.58E-07 | 0.771 |
| 8 | 16,714,002 | CG | 0.7161 | 2.05E-07 | 0.551 | 6 | 44,792,723 | CG | 0.6532 | 6.54E-06 | 0.676 |
| 8 | 16,714,003 | CG | 0.6686 | 2.41E-06 | 0.533 | 6 | 44,792,729 | CG | 0.6768 | 2.24E-06 | 0.694 |
| 8 | 17,407,748 | CG | 0.6851 | 1.50E-06 | 0.630 | 6 | 44,844,285 | CG | 0.7423 | 4.03E-06 | 0.685 |
| 8 | 23,137,505 | CG | 0.6825 | 1.23E-06 | 0.716 | 6 | 44,844,298 | CG | 0.7699 | 1.04E-06 | 0.750 |
| 8 | 25,713,678 | CG | 0.8131 | 1.56E-08 | 0.726 | 6 | 44,891,925 | CG | 0.7066 | 7.12E-07 | 0.658 |
| 8 | 33,653,700 | CG | 0.6891 | 8.79E-07 | 0.673 | 6 | 44,891,939 | CG | 0.7937 | 2.77E-09 | 0.809 |
| Chromosome | position | context | Spearman Rho | P-value | $h^2$ | Chromosome | position | context | Spearman Rho | P-value | $h^2$ |
| 8 | 48,305,917 | CG | 0.6975 | 8.03E-07 | 0.605 | 6 | 44,891,945 | CG | 0.7661 | 2.07E-08 | 0.758 |
| 8 | 49,145,729 | CG | 0.7554 | 1.56E-07 | 0.610 | 6 | 44,891,947 | CG | 0.7999 | 1.69E-09 | 0.819 |
| 8 | 58,014,945 | CG | 0.6539 | 4.75E-06 | 0.558 | 6 | 44,891,951 | CG | 0.8114 | 6.47E-10 | 0.858 |
| 8 | 58,014,954 | CG | 0.7381 | 3.32E-07 | 0.711 | 6 | 44,891,969 | CG | 0.8113 | 6.54E-10 | 0.940 |
| 8 | 58,014,955 | CG | 0.6586 | 3.84E-06 | 0.724 | 6 | 44,891,975 | CG | 0.7278 | 2.24E-07 | 0.789 |
| 8 | 58,014,956 | CG | 0.6433 | 7.55E-06 | 0.706 | 6 | 44,891,978 | CG | 0.7660 | 2.10E-08 | 0.899 |
| 8 | 58,014,981 | CG | 0.6565 | 4.23E-06 | 0.523 | 6 | 45,716,210 | CG | 0.7247 | 1.82E-07 | 0.529 |
| 8 | 58,776,430 | CG | 0.7657 | 2.05E-06 | 0.780 | 6 | 45,716,219 | CG | 0.7422 | 4.21E-08 | 0.538 |
| 8 | 62,651,282 | CG | 0.6707 | 2.18E-06 | 0.626 | 6 | 51,995,234 | CG | 0.7929 | 1.36E-06 | 0.535 |
| 8 | 62,910,122 | CG | 0.6974 | 6.49E-06 | 0.564 | 6 | 52,916,295 | CG | 0.7233 | 6.23E-07 | 0.675 |
| 9 | 9,183,461 | CG | 0.6991 | 5.21E-07 | 0.633 | 7 | 779,375 | CG | 0.6812 | 1.31E-06 | 0.538 |
| 9 | 9,510,244 | CG | 0.7809 | 2.78E-09 | 0.565 | 7 | 1,533,735 | CG | 0.7351 | 7.38E-07 | 0.557 |
| 9 | 11,601,037 | CG | 0.7189 | 1.68E-06 | 0.548 | 7 | 3,742,239 | CG | 0.7339 | 3.93E-06 | 0.688 |
| 9 | 14,325,010 | CG | 0.7949 | 9.08E-10 | 0.684 | 7 | 5,226,521 | CG | 0.6844 | 2.99E-06 | 0.632 |
| 9 | 14,325,017 | CG | 0.8106 | 2.32E-10 | 0.768 | 7 | 8,207,154 | CG | 0.7270 | 5.13E-07 | 0.663 |
| 9 | 14,325,032 | CG | 0.7498 | 2.57E-08 | 0.757 | 7 | 10,175,706 | CG | 0.7289 | 1.43E-07 | 0.614 |
| 9 | 14,325,035 | CG | 0.8135 | 1.76E-10 | 0.761 | 7 | 11,817,509 | CG | 0.7481 | 6.70E-08 | 0.605 |
| 9 | 14,325,036 | CG | 0.8206 | 2.82E-09 | 0.736 | 7 | 12,989,161 | CG | 0.6434 | 7.52E-06 | 0.567 |
| 9 | 14,325,063 | CG | 0.8382 | 1.49E-11 | 0.735 | 7 | 23,521,205 | CG | 0.7908 | 1.27E-09 | 0.721 |
| 9 | 14,325,064 | CG | 0.8319 | 2.92E-11 | 0.686 | 7 | 27,587,547 | CG | 0.6519 | 9.27E-06 | 0.715 |
| 9 | 16,964,059 | CG | 0.6728 | 1.97E-06 | 0.490 | 7 | 34,253,352 | CG | 0.6492 | 7.78E-06 | 0.628 |
| 9 | 17,813,026 | CG | 0.6502 | 5.59E-06 | 0.547 | 7 | 34,266,134 | CG | 0.7290 | 9.58E-08 | 0.547 |
| 9 | 19,609,829 | CG | 0.6963 | 8.56E-07 | 0.581 | 7 | 34,345,427 | CG | 0.6468 | 8.62E-06 | 0.640 |
| 9 | 25,795,359 | CG | 0.7695 | 6.91E-06 | 0.716 | 7 | 34,345,428 | CG | 0.6963 | 6.07E-07 | 0.687 |
| 9 | 32,402,361 | CG | 0.7315 | 8.22E-08 | 0.587 | 7 | 34,345,444 | CG | 0.6821 | 1.25E-06 | 0.602 |
| 9 | 32,704,350 | CG | 0.6782 | 5.50E-06 | 0.586 | 7 | 35,395,393 | CG | 0.7252 | 1.77E-07 | 0.466 |
| 9 | 36,008,687 | CG | 0.7679 | 7.35E-09 | 0.672 | 7 | 36,775,415 | CG | 0.6913 | 7.88E-07 | 0.702 |
| 9 | 36,188,650 | CG | 0.7475 | 4.56E-08 | 0.552 | 7 | 36,775,416 | CG | 0.7514 | 5.46E-08 | 0.734 |
| 9 | 38,248,581 | CG | 0.6385 | 9.28E-06 | 0.587 | 7 | 38,624,494 | CG | 0.7039 | 8.20E-07 | 0.676 |
| Chromosome | position | context | Spearman Rho | P-value | $h^2$ | Chromosome | position | context | Spearman Rho | P-value | $h^2$ |
| 9 | 41,112,285 | CG | 0.6479 | 6.19E-06 | 0.511 | 7 | 44,342,017 | CG | 0.6772 | 2.20E-06 | 0.578 |
| 9 | 42,882,296 | CG | 0.6584 | 3.88E-06 | 0.602 | 7 | 44,342,018 | CG | 0.6597 | 6.64E-06 | 0.571 |
| 9 | 42,924,310 | CG | 0.7218 | 1.47E-07 | 0.402 | 8 | 1,574,991 | CG | 0.7340 | 1.17E-06 | 0.432 |
| 9 | 46,136,029 | CG | 0.6890 | 8.83E-07 | 0.613 | 8 | 1,574,992 | CG | 0.7217 | 2.14E-06 | 0.470 |
| 10 | 157,720 | CG | 0.6972 | 5.77E-07 | 0.619 | 8 | 3,516,129 | CG | 0.6923 | 2.90E-06 | 0.527 |
| 10 | 3,221,127 | CG | 0.6773 | 7.88E-06 | 0.549 | 8 | 4,007,469 | CG | 0.7386 | 7.19E-06 | 0.703 |
| 10 | 3,221,177 | CG | 0.7300 | 9.62E-07 | 0.584 | 8 | 4,007,783 | CG | 0.6444 | 7.22E-06 | 0.615 |
| 10 | 3,221,178 | CG | 0.7280 | 2.34E-06 | 0.582 | 8 | 4,007,784 | CG | 0.6614 | 3.38E-06 | 0.539 |
| 10 | 5,871,405 | CG | 0.6850 | 2.92E-06 | 0.482 | 8 | 4,607,405 | CG | 0.6582 | 5.26E-06 | 0.653 |
| 10 | 6,054,630 | CG | 0.6510 | 9.61E-06 | 0.454 | 8 | 11,744,222 | CG | 0.6806 | 9.46E-06 | 0.491 |
| 10 | 15,984,235 | CG | 0.6692 | 2.34E-06 | 0.564 | 8 | 16,714,003 | CG | 0.6887 | 9.01E-07 | 0.503 |
| 10 | 17,715,567 | CG | 0.6397 | 8.83E-06 | 0.753 | 8 | 19,711,558 | CG | 0.7447 | 8.28E-08 | 0.596 |
| 10 | 17,715,663 | CG | 0.6460 | 6.74E-06 | 0.568 | 8 | 19,711,593 | CG | 0.6970 | 1.17E-06 | 0.789 |
| 10 | 17,715,728 | CG | 0.6777 | 1.56E-06 | 0.628 | 8 | 21,666,091 | CG | 0.7089 | 3.07E-07 | 0.652 |
| 10 | 20,404,647 | CG | 0.6739 | 1.87E-06 | 0.509 | 8 | 22,063,842 | CG | 0.7058 | 5.19E-07 | 0.483 |
| 10 | 23,460,067 | CG | 0.6965 | 4.77E-06 | 0.604 | 8 | 23,137,505 | CG | 0.6503 | 5.56E-06 | 0.682 |
| 10 | 25,631,933 | CG | 0.6834 | 1.17E-06 | 0.507 | 8 | 26,857,425 | CG | 0.6420 | 8.02E-06 | 0.521 |
| 10 | 31,672,087 | CG | 0.7858 | 1.36E-08 | 0.632 | 8 | 27,974,900 | CG | 0.6983 | 5.46E-07 | 0.578 |
| 10 | 32,869,251 | CG | 0.6927 | 2.02E-06 | 0.495 | 8 | 33,653,700 | CG | 0.7425 | 4.15E-08 | 0.769 |
| 10 | 33,677,641 | CG | 0.6846 | 4.12E-06 | 0.663 | 8 | 41,842,754 | CG | 0.7098 | 2.91E-07 | 0.638 |
| 10 | 40,166,644 | CG | 0.6479 | 6.20E-06 | 0.595 | 8 | 43,122,541 | CG | 0.6482 | 8.13E-06 | 0.042 |
| 10 | 41,121,153 | CG | 0.7613 | 4.21E-07 | 0.602 | 8 | 44,644,794 | CG | 0.6608 | 6.32E-06 | 1.302 |
| 10 | 48,416,376 | CG | 0.6900 | 1.64E-06 | 0.615 | 8 | 49,074,454 | CG | 0.6760 | 1.69E-06 | 0.456 |
| 10 | 52,086,849 | CG | 0.6841 | 4.23E-06 | 0.461 | 8 | 49,145,729 | CG | 0.6840 | 8.19E-06 | 0.748 |
| 10 | 52,086,860 | CG | 0.7372 | 2.92E-07 | 0.582 | 8 | 54,302,435 | CG | 0.6859 | 1.04E-06 | 0.534 |
| 10 | 52,086,863 | CG | 0.6680 | 8.54E-06 | 0.567 | 8 | 56,313,555 | CG | 0.7018 | 1.85E-06 | 0.434 |
| 10 | 56,527,258 | CG | 0.7242 | 5.95E-07 | 0.814 | 8 | 56,726,312 | CG | 0.6434 | 9.97E-06 | 0.548 |
| 10 | 56,533,477 | CG | 0.7556 | 4.17E-08 | 0.564 | 8 | 58,014,954 | CG | 0.7047 | 1.08E-06 | 0.635 |
| 10 | 56,533,478 | CG | 0.7686 | 6.98E-09 | 0.619 | 8 | 58,014,981 | CG | 0.6718 | 2.08E-06 | 0.492 |
| Chromosome | position | context | Spearman Rho | P-value | $h^2$ | Chromosome | position | context | Spearman Rho | P-value | $h^2$ |
| 11 | 1,041,257 | CG | 0.6818 | 1.27E-06 | 0.686 | 8 | 62,910,122 | CG | 0.7305 | 3.08E-06 | 0.739 |
| 11 | 2,811,974 | CG | 0.7263 | 3.61E-07 | 0.547 | 9 | 6,478,385 | CG | 0.6895 | 1.69E-06 | 0.616 |
| 11 | 4,793,544 | CG | 0.6413 | 8.25E-06 | 0.595 | 9 | 9,510,244 | CG | 0.6532 | 4.90E-06 | 0.477 |
| 11 | 7,411,150 | CG | 0.7693 | 2.65E-08 | 0.555 | 9 | 10,577,112 | CG | 0.6687 | 4.46E-06 | 0.485 |
| 11 | 7,411,159 | CG | 0.6887 | 2.46E-06 | 0.458 | 9 | 11,600,998 | CG | 0.7778 | 6.20E-08 | 0.633 |
| 11 | 7,411,233 | CG | 0.7109 | 1.70E-06 | 0.561 | 9 | 11,601,037 | CG | 0.7343 | 5.14E-07 | 0.542 |
| 11 | 7,411,239 | CG | 0.7229 | 9.37E-07 | 0.514 | 9 | 13,406,163 | CG | 0.6760 | 2.32E-06 | 0.506 |
| 11 | 7,411,240 | CG | 0.7072 | 4.83E-07 | 0.519 | 9 | 13,406,164 | CG | 0.7106 | 2.79E-07 | 0.531 |
| 11 | 7,411,243 | CG | 0.7145 | 3.24E-07 | 0.554 | 9 | 14,325,010 | CG | 0.6691 | 2.36E-06 | 0.507 |
| 11 | 7,411,244 | CG | 0.7404 | 1.07E-07 | 0.600 | 9 | 14,325,017 | CG | 0.7275 | 1.05E-07 | 0.514 |
| 11 | 7,411,260 | CG | 0.7009 | 6.75E-07 | 0.550 | 9 | 14,325,032 | CG | 0.6901 | 8.37E-07 | 0.514 |
| 11 | 7,411,264 | CG | 0.7283 | 1.47E-07 | 0.585 | 9 | 14,325,035 | CG | 0.7242 | 1.28E-07 | 0.492 |
| 11 | 7,411,277 | CG | 0.6923 | 1.05E-06 | 0.524 | 9 | 14,325,036 | CG | 0.7902 | 1.65E-08 | 0.493 |
| 11 | 8,788,955 | CG | 0.6716 | 2.86E-06 | 1.184 | 9 | 14,325,063 | CG | 0.7372 | 5.79E-08 | 0.570 |
| 11 | 20,646,562 | CG | 0.7114 | 3.85E-07 | 0.684 | 9 | 14,325,064 | CG | 0.7555 | 1.75E-08 | 0.527 |
| 11 | 20,646,728 | CG | 0.6394 | 8.95E-06 | 0.523 | 9 | 16,964,082 | CG | 0.6761 | 2.32E-06 | 0.466 |
| 11 | 20,646,734 | CG | 0.6694 | 2.33E-06 | 0.537 | 9 | 16,964,197 | CG | 0.7383 | 4.43E-07 | 0.679 |
| 11 | 20,646,740 | CG | 0.6574 | 4.07E-06 | 0.481 | 9 | 17,813,025 | CG | 0.7284 | 9.97E-08 | 0.642 |
| 11 | 24,048,594 | CG | 0.7416 | 3.45E-07 | 0.686 | 9 | 17,983,597 | CG | 0.6536 | 4.81E-06 | 0.529 |
| 11 | 25,885,177 | CG | 0.6874 | 9.61E-07 | 0.502 | 9 | 19,609,706 | CG | 0.6829 | 4.46E-06 | 0.561 |
| 11 | 25,885,178 | CG | 0.6517 | 5.23E-06 | 0.492 | 9 | 20,810,601 | CG | 0.7024 | 6.22E-07 | 0.603 |
| 11 | 27,678,945 | CG | 0.6452 | 9.24E-06 | 0.540 | 9 | 27,499,880 | CG | 0.7303 | 6.89E-06 | 0.702 |
| 11 | 29,908,329 | CG | 0.7999 | 1.38E-08 | 0.573 | 9 | 29,579,875 | CG | 0.6532 | 6.55E-06 | 0.609 |
| 11 | 29,908,350 | CG | 0.6847 | 7.95E-06 | 0.405 | 9 | 32,704,350 | CG | 0.6801 | 1.91E-06 | 0.601 |
| 11 | 29,908,359 | CG | 0.7783 | 6.02E-08 | 0.548 | 9 | 34,578,395 | CG | 0.6647 | 3.93E-06 | 0.715 |
| 11 | 29,908,367 | CG | 0.8023 | 1.16E-08 | 0.651 | 9 | 36,008,687 | CG | 0.7013 | 4.65E-07 | 0.573 |
| 11 | 36,019,076 | CG | 0.6530 | 6.62E-06 | 0.496 | 9 | 36,188,650 | CG | 0.6934 | 9.93E-07 | 0.556 |
| 11 | 43,338,065 | CG | 0.6754 | 1.74E-06 | 0.616 | 9 | 38,263,243 | CG | 0.6466 | 6.57E-06 | 0.633 |
| 11 | 43,338,111 | CG | 0.7029 | 4.27E-07 | 0.616 | 9 | 44,566,060 | CG | 0.7726 | 5.21E-09 | 0.718 |
| Chromo-<br>some | position | context | Spearman<br>Rho | P-value | $h^2$ | Chromo-<br>some | position | context | Spearman<br>Rho | P-value | $h^2$ |
| 11 | 43,338,153 | CG | 0.6750 | 1.78E-06 | 0.619 | 9 | 44,640,742 | CG | 0.6693 | 2.34E-06 | 0.515 |
| 11 | 43,338,156 | CG | 0.6573 | 4.07E-06 | 0.624 | 9 | 44,640,745 | CG | 0.6726 | 1.99E-06 | 0.606 |
| 11 | 47,895,372 | CG | 0.7045 | 5.57E-07 | 0.453 | 10 | 157,715 | CG | 0.7022 | 4.43E-07 | 0.610 |
| 11 | 50,136,691 | CG | 0.6571 | 4.11E-06 | 0.457 | 10 | 3,221,142 | CG | 0.7417 | 3.43E-07 | 0.459 |
| 11 | 50,940,529 | CG | 0.6864 | 5.32E-06 | 0.560 | 10 | 3,882,297 | CG | 0.7282 | 5.09E-06 | 0.582 |
| 11 | 52,348,206 | CG | 0.7139 | 2.31E-07 | 0.529 | 10 | 4,729,802 | CG | 0.7965 | 2.37E-07 | 0.576 |
| 11 | 52,515,086 | CG | 0.8032 | 5.38E-08 | 0.554 | 10 | 5,871,405 | CG | 0.7106 | 4.02E-07 | 0.572 |
| 12 | 1,577,516 | CG | 0.7913 | 2.03E-09 | 0.648 | 10 | 5,871,406 | CG | 0.6553 | 4.46E-06 | 0.582 |
| 12 | 1,577,608 | CG | 0.7487 | 5.42E-07 | 0.598 | 10 | 13,673,351 | CG | 0.6939 | 6.88E-07 | 0.638 |
| 12 | 3,832,647 | CG | 0.6902 | 2.28E-06 | 0.709 | 10 | 13,673,352 | CG | 0.7302 | 8.89E-08 | 0.646 |
| 12 | 4,711,708 | CG | 0.7073 | 2.92E-06 | 0.569 | 10 | 17,715,663 | CG | 0.6649 | 2.87E-06 | 0.660 |
| 12 | 4,711,852 | CG | 0.7158 | 2.82E-06 | 0.513 | 10 | 18,573,703 | CG | 0.6526 | 6.73E-06 | 0.577 |
| 12 | 5,108,467 | CG | 0.7024 | 7.42E-06 | 0.742 | 10 | 20,404,646 | CG | 0.7332 | 1.11E-07 | 0.701 |
| 12 | 6,960,880 | CG | 0.6659 | 3.72E-06 | 0.623 | 10 | 20,404,647 | CG | 0.7511 | 2.36E-08 | 0.648 |
| 12 | 7,994,192 | CG | 0.6478 | 8.26E-06 | 0.671 | 10 | 25,631,933 | CG | 0.8303 | 3.45E-11 | 0.654 |
| 12 | 8,928,713 | CG | 0.6697 | 2.29E-06 | 0.766 | 10 | 33,059,460 | CG | 0.6723 | 2.02E-06 | 0.631 |
| 12 | 8,928,738 | CG | 0.6506 | 5.50E-06 | 0.618 | 10 | 33,184,792 | CG | 0.6699 | 2.27E-06 | 0.498 |
| 12 | 8,928,739 | CG | 0.6943 | 6.73E-07 | 0.769 | 10 | 33,184,793 | CG | 0.6819 | 1.27E-06 | 0.499 |
| 12 | 8,928,740 | CG | 0.6934 | 7.07E-07 | 0.628 | 10 | 33,677,641 | CG | 0.6577 | 9.71E-06 | 0.552 |
| 12 | 12,085,721 | CG | 0.7164 | 2.01E-07 | 0.527 | 10 | 36,875,033 | CG | 0.6778 | 2.13E-06 | 0.747 |
| 12 | 14,900,665 | CG | 0.7850 | 3.38E-06 | 0.793 | 10 | 40,753,488 | CG | 0.6855 | 1.47E-06 | 0.708 |
| 12 | 17,585,888 | CG | 0.7446 | 3.62E-08 | 0.529 | 10 | 41,264,568 | CG | 0.7047 | 3.28E-06 | 0.680 |
| 12 | 21,910,448 | CG | 0.6887 | 1.25E-06 | 0.388 | 10 | 46,702,704 | CG | 0.6745 | 3.43E-06 | 0.627 |
| 12 | 22,356,947 | CG | 0.6503 | 5.57E-06 | 0.437 | 10 | 48,416,376 | CG | 0.7599 | 4.91E-08 | 0.588 |
| 12 | 24,019,243 | CG | 0.7632 | 2.41E-07 | 0.577 | 10 | 51,760,452 | CG | 0.7771 | 9.62E-09 | 0.676 |
| 12 | 24,642,505 | CG | 0.6553 | 8.02E-06 | 0.512 | 10 | 56,533,711 | CG | 0.7567 | 9.32E-08 | 0.774 |
| 12 | 27,331,238 | CG | 0.7118 | 7.84E-07 | 0.602 | 11 | 1,041,257 | CG | 0.6838 | 1.15E-06 | 0.585 |
| 12 | 30,788,253 | CG | 0.6487 | 5.98E-06 | 0.618 | 11 | 8,739,121 | CG | 0.7085 | 2.75E-06 | 0.584 |
| 12 | 33,671,473 | CG | 0.6840 | 5.90E-06 | 0.512 | 11 | 10,385,335 | CG | 0.6944 | 1.33E-06 | 0.569 |
| Chromosome | position | context | Spearman Rho | P-value | $h^2$ | Chromosome | position | context | Spearman Rho | P-value | $h^2$ |
| 12 | 34,601,942 | CG | 0.7338 | 1.59E-07 | 0.642 | 11 | 13,552,314 | CG | 0.7386 | 1.20E-07 | 0.631 |
| 12 | 36,187,235 | CG | 0.7528 | 2.10E-08 | 0.686 | 11 | 13,585,878 | CG | 0.7152 | 9.50E-07 | 0.655 |
| 12 | 37,925,182 | CG | 0.7480 | 2.90E-08 | 0.513 | 11 | 13,673,401 | CG | 0.7602 | 1.27E-08 | 0.669 |
| 12 | 40,378,040 | CG | 0.7564 | 1.65E-08 | 0.667 | 11 | 13,673,405 | CG | 0.7531 | 2.06E-08 | 0.623 |
| 12 | 42,419,241 | CG | 0.6482 | 6.12E-06 | 0.455 | 11 | 13,673,631 | CG | 0.7497 | 2.59E-08 | 0.762 |
| 12 | 43,065,113 | CG | 0.7836 | 8.16E-07 | 0.698 | 11 | 13,673,632 | CG | 0.7615 | 1.16E-08 | 0.785 |
|  |  |  |  |  |  | 11 | 20,646,734 | CG | 0.6687 | 2.41E-06 | 0.469 |
|  |  |  |  |  |  | 11 | 21,650,661 | CG | 0.6425 | 7.85E-06 | 0.308 |
|  |  |  |  |  |  | 11 | 24,048,676 | CG | 0.6647 | 2.90E-06 | 0.586 |
|  |  |  |  |  |  | 11 | 25,885,178 | CG | 0.6593 | 3.71E-06 | 0.425 |
|  |  |  |  |  |  | 11 | 30,658,274 | CG | 0.7223 | 2.09E-07 | 0.652 |
|  |  |  |  |  |  | 11 | 32,591,404 | CG | 0.7039 | 4.03E-07 | 0.609 |
|  |  |  |  |  |  | 11 | 32,591,414 | CG | 0.6694 | 2.33E-06 | 0.565 |
|  |  |  |  |  |  | 11 | 32,591,416 | CG | 0.6575 | 4.04E-06 | 0.547 |
|  |  |  |  |  |  | 11 | 32,591,421 | CG | 0.6924 | 7.45E-07 | 0.533 |
|  |  |  |  |  |  | 11 | 32,591,422 | CG | 0.6522 | 5.13E-06 | 0.424 |
|  |  |  |  |  |  | 11 | 36,346,451 | CG | 0.7788 | 1.72E-06 | 0.759 |
|  |  |  |  |  |  | 11 | 38,639,149 | CG | 0.7145 | 2.24E-07 | 0.647 |
|  |  |  |  |  |  | 11 | 38,639,150 | CG | 0.6389 | 9.14E-06 | 0.618 |
|  |  |  |  |  |  | 11 | 43,432,220 | CG | 0.6537 | 4.79E-06 | 0.499 |
|  |  |  |  |  |  | 11 | 45,808,516 | CG | 0.7502 | 5.00E-07 | 0.563 |
|  |  |  |  |  |  | 11 | 46,825,622 | CG | 0.7009 | 1.93E-06 | 0.655 |
|  |  |  |  |  |  | 11 | 50,136,891 | CG | 0.6456 | 6.86E-06 | 0.657 |
|  |  |  |  |  |  | 11 | 52,348,125 | CG | 0.6701 | 5.71E-06 | 0.474 |
|  |  |  |  |  |  | 12 | 1,008,589 | CG | 0.7327 | 7.65E-08 | 0.763 |
|  |  |  |  |  |  | 12 | 1,008,595 | CG | 0.6769 | 1.62E-06 | 0.557 |
|  |  |  |  |  |  | 12 | 1,573,993 | CG | 0.7132 | 3.19E-06 | 0.726 |
|  |  |  |  |  |  | 12 | 1,577,516 | CG | 0.7094 | 4.29E-07 | 0.620 |
|  |  |  |  |  |  | 12 | 4,711,669 | CG | 0.6849 | 1.52E-06 | 0.765 |
| Chromo-<br>some | position | context | Spearman<br>Rho | P-value | $h^2$ | Chromo-<br>some | position | context | Spearman<br>Rho | P-value | $h^2$ |
|  |  |  |  |  |  | 12 | 4,711,852 | CG | 0.6603 | 8.73E-06 | 0.495 |
|  |  |  |  |  |  | 12 | 16,225,965 | CG | 0.6599 | 4.87E-06 | 0.554 |
|  |  |  |  |  |  | 12 | 17,585,889 | CG | 0.7080 | 9.52E-07 | 0.562 |
|  |  |  |  |  |  | 12 | 20,167,101 | CG | 0.6664 | 2.68E-06 | 0.659 |
|  |  |  |  |  |  | 12 | 21,293,828 | CG | 0.7445 | 2.39E-06 | 0.646 |
|  |  |  |  |  |  | 12 | 24,019,224 | CG | 0.7134 | 3.16E-06 | 0.629 |
|  |  |  |  |  |  | 12 | 24,019,243 | CG | 0.7308 | 1.37E-06 | 0.657 |
|  |  |  |  |  |  | 12 | 24,019,283 | CG | 0.7456 | 6.41E-07 | 0.657 |
|  |  |  |  |  |  | 12 | 24,642,505 | CG | 0.6515 | 9.41E-06 | 0.562 |
|  |  |  |  |  |  | 12 | 24,665,540 | CG | 0.6711 | 2.14E-06 | 0.357 |
|  |  |  |  |  |  | 12 | 25,365,204 | CG | 0.7491 | 2.26E-07 | 0.602 |
|  |  |  |  |  |  | 12 | 27,331,201 | CG | 0.6934 | 9.90E-07 | 0.664 |
|  |  |  |  |  |  | 12 | 27,331,237 | CG | 0.8035 | 1.54E-07 | 0.614 |
|  |  |  |  |  |  | 12 | 36,187,357 | CG | 0.7003 | 4.91E-07 | 0.652 |
|  |  |  |  |  |  | 12 | 36,187,365 | CG | 0.7378 | 5.59E-08 | 0.729 |
|  |  |  |  |  |  | 12 | 37,925,182 | CG | 0.6545 | 4.61E-06 | 0.489 |
|  |  |  |  |  |  | 12 | 39,478,979 | CG | 0.7483 | 8.46E-07 | 0.566 |
|  |  |  |  |  |  | 12 | 40,547,728 | CG | 0.6557 | 4.39E-06 | 0.659 |
| 1 | 16,747,844 | CHG | 0.7124 | 2.29E-06 | 0.590 | 1 | 3,014,526 | CHG | 0.7578 | 3.01E-06 | 0.592 |
| 1 | 26,634,624 | CHG | 0.7235 | 2.89E-06 | 0.373 | 1 | 3,014,559 | CHG | 0.7651 | 1.34E-06 | 0.563 |
| 1 | 38,054,491 | CHG | 0.6486 | 8.00E-06 | 1.578 | 1 | 3,014,565 | CHG | 0.7228 | 9.50E-06 | 0.483 |
| 2 | 12,063,886 | CHG | 0.7207 | 1.05E-06 | 0.415 | 1 | 3,014,616 | CHG | 0.7305 | 6.84E-06 | 0.553 |
| 2 | 12,063,896 | CHG | 0.7113 | 5.57E-07 | 0.374 | 1 | 3,014,718 | CHG | 0.6610 | 3.44E-06 | 0.478 |
| 2 | 12,063,898 | CHG | 0.8006 | 7.73E-09 | 0.357 | 1 | 3,014,744 | CHG | 0.6687 | 2.41E-06 | 0.496 |
| 2 | 12,063,903 | CHG | 0.7122 | 5.32E-07 | 0.419 | 1 | 3,014,746 | CHG | 0.6437 | 7.42E-06 | 0.442 |
| 2 | 12,063,905 | CHG | 0.7410 | 3.58E-07 | 0.351 | 1 | 13,559,818 | CHG | 0.6996 | 2.92E-06 | 0.410 |
| 2 | 12,063,910 | CHG | 0.6958 | 3.48E-06 | 0.508 | 1 | 20,119,685 | CHG | 0.7164 | 1.30E-06 | 0.853 |
| 2 | 32,680,717 | CHG | 0.7148 | 2.96E-06 | 0.619 | 1 | 21,485,450 | CHG | 0.6409 | 8.37E-06 | 0.537 |
| 2 | 48,467,533 | CHG | 0.7246 | 8.59E-07 | 0.592 | 1 | 21,485,483 | CHG | 0.6819 | 1.27E-06 | 0.469 |
| Chromosome | position | context | Spearman Rho | P-value | $h^2$ | Chromosome | position | context | Spearman Rho | P-value | $h^2$ |
| 2 | 63,623,271 | CHG | 0.7252 | 5.79E-06 | 0.536 | 1 | 46,365,911 | CHG | 0.7454 | 6.48E-07 | 0.531 |
| 2 | 101,304,575 | CHG | 0.6766 | 4.27E-06 | 0.525 | 1 | 54,692,867 | CHG | 0.7460 | 1.46E-06 | 0.660 |
| 3 | 28,880,879 | CHG | 0.7024 | 3.65E-06 | 0.491 | 1 | 54,692,926 | CHG | 0.7463 | 1.43E-06 | 0.702 |
| 3 | 32,073,745 | CHG | 0.7174 | 5.58E-06 | 0.588 | 2 | 12,063,886 | CHG | 0.7173 | 8.52E-07 | 0.440 |
| 4 | 16,404,054 | CHG | 0.7360 | 8.05E-06 | 0.552 | 2 | 12,063,898 | CHG | 0.7938 | 7.66E-09 | 0.402 |
| 4 | 22,955,994 | CHG | 0.7192 | 5.16E-06 | 1.246 | 2 | 12,063,905 | CHG | 0.7900 | 1.01E-08 | 0.448 |
| 4 | 33,656,275 | CHG | 0.8096 | 5.47E-07 | 0.777 | 2 | 12,063,910 | CHG | 0.7319 | 3.92E-07 | 0.466 |
| 4 | 53,005,263 | CHG | 0.6514 | 5.31E-06 | 0.731 | 4 | 10,213,343 | CHG | 0.6976 | 7.98E-07 | 0.554 |
| 4 | 66,271,684 | CHG | 0.7071 | 6.06E-06 | 0.361 | 4 | 24,845,288 | CHG | 0.6935 | 9.86E-07 | 0.792 |
| 5 | 57,608,753 | CHG | 0.6704 | 2.22E-06 | 0.424 | 4 | 48,548,899 | CHG | 0.7423 | 1.45E-07 | 0.752 |
| 5 | 67,135,192 | CHG | 0.6506 | 5.51E-06 | 0.472 | 5 | 28,709,334 | CHG | 0.7844 | 4.78E-07 | 0.216 |
| 6 | 44,891,972 | CHG | 0.6858 | 3.90E-06 | 0.506 | 5 | 28,709,358 | CHG | 0.7419 | 4.10E-06 | 0.206 |
| 6 | 47,513,375 | CHG | 0.6475 | 8.39E-06 | 0.223 | 5 | 28,709,362 | CHG | 0.7281 | 7.56E-06 | 0.236 |
| 7 | 20,554,107 | CHG | 0.6545 | 4.63E-06 | 0.398 | 5 | 28,709,377 | CHG | 0.7523 | 2.53E-06 | 0.219 |
| 7 | 35,806,803 | CHG | 0.7486 | 9.93E-08 | 0.613 | 5 | 57,991,202 | CHG | 0.7361 | 5.33E-06 | 0.276 |
| 9 | 34,656,368 | CHG | 0.6472 | 6.38E-06 | 0.103 | 5 | 67,135,192 | CHG | 0.6905 | 8.20E-07 | 0.554 |
| 10 | 2,459,641 | CHG | 0.7325 | 4.19E-06 | 0.698 | 6 | 1,116,149 | CHG | 0.7452 | 2.31E-06 | 0.519 |
| 10 | 3,221,211 | CHG | 0.7047 | 3.28E-06 | 0.427 | 6 | 44,891,958 | CHG | 0.7309 | 1.89E-07 | 0.603 |
| 10 | 38,547,322 | CHG | 0.6993 | 7.34E-07 | 0.772 | 9 | 1,480,030 | CHG | 0.7830 | 2.26E-06 | 0.747 |
| 11 | 7,411,154 | CHG | 0.6586 | 9.38E-06 | 0.362 | 9 | 29,579,870 | CHG | 0.6873 | 1.35E-06 | 0.656 |
| 11 | 7,411,270 | CHG | 0.7116 | 3.79E-07 | 0.483 | 9 | 32,704,322 | CHG | 0.6747 | 2.48E-06 | 0.563 |
| 11 | 23,344,901 | CHG | 0.7012 | 5.48E-06 | 0.574 | 9 | 32,704,332 | CHG | 0.7018 | 6.44E-07 | 0.560 |
|  |  |  |  |  |  | 9 | 32,704,400 | CHG | 0.6724 | 2.75E-06 | 0.524 |
|  |  |  |  |  |  | 10 | 38,547,309 | CHG | 0.7208 | 1.52E-06 | 0.624 |
|  |  |  |  |  |  | 10 | 49,058,871 | CHG | 0.7026 | 7.36E-06 | 0.740 |
|  |  |  |  |  |  | 11 | 27,923,239 | CHG | 0.7520 | 4.51E-07 | 0.439 |
|  |  |  |  |  |  | 11 | 36,346,450 | CHG | 0.8092 | 3.23E-07 | 0.556 |
|  |  |  |  |  |  | 11 | 38,639,144 | CHG | 0.6848 | 1.09E-06 | 0.557 |
|  |  |  |  |  |  | 12 | 24,019,257 | CHG | 0.7168 | 2.69E-06 | 0.712 |
| Chromo-<br>some | position | context | Spearman<br>Rho | P-value | $h^2$ | Chromo-<br>some | position | context | Spearman<br>Rho | P-value | $h^2$ |
| 1 | 3,014,705 | CHH | 0.6785 | 1.50E-06 | 0.375 | 1 | 3,014,540 | CHH | 0.7269 | 7.98E-06 | 0.081 |
| 1 | 7,125,376 | CHH | 0.6466 | 6.55E-06 | 0.554 | 1 | 3,014,779 | CHH | 0.6969 | 5.88E-07 | 0.157 |
| 1 | 8,066,570 | CHH | 0.6868 | 1.93E-06 | 0.527 | 1 | 8,066,570 | CHH | 0.6910 | 3.09E-06 | 0.354 |
| 2 | 38,816,761 | CHH | 0.8859 | 8.59E-09 | 0.833 | 1 | 10,546,825 | CHH | 0.7710 | 1.58E-06 | 0.662 |
| 3 | 18,181,075 | CHH | 0.6964 | 6.02E-07 | 0.289 | 1 | 14,506,740 | CHH | 0.6653 | 3.82E-06 | 0.313 |
| 3 | 18,181,098 | CHH | 0.6570 | 7.46E-06 | 0.221 | 1 | 19,332,195 | CHH | 0.6516 | 5.27E-06 | 0.332 |
| 3 | 35,023,827 | CHH | 0.7298 | 9.16E-08 | 0.548 | 1 | 21,485,411 | CHH | 0.6963 | 6.07E-07 | 0.242 |
| 3 | 50,045,982 | CHH | 0.7677 | 4.72E-06 | 0.670 | 1 | 46,366,763 | CHH | 0.7757 | 6.32E-06 | 0.540 |
| 4 | 33,540,044 | CHH | 0.6972 | 1.16E-06 | 0.556 | 1 | 47,296,506 | CHH | 0.7532 | 7.49E-08 | 0.650 |
| 4 | 48,548,901 | CHH | 0.6690 | 8.18E-06 | 0.243 | 2 | 19,650,266 | CHH | 0.6573 | 7.36E-06 | 0.552 |
| 4 | 52,271,883 | CHH | 0.7140 | 2.31E-07 | 0.561 | 2 | 63,536,637 | CHH | 0.6855 | 2.84E-06 | 0.283 |
| 4 | 76,283,198 | CHH | 0.6448 | 7.08E-06 | 0.374 | 2 | 92,135,478 | CHH | 0.7611 | 4.03E-06 | 0.670 |
| 5 | 7,557,888 | CHH | 0.6907 | 1.14E-06 | 0.390 | 3 | 33,923,261 | CHH | 0.6796 | 5.17E-06 | 0.623 |
| 5 | 19,538,109 | CHH | 0.6511 | 5.38E-06 | 0.565 | 3 | 33,923,281 | CHH | 0.7263 | 5.33E-07 | 0.643 |
| 5 | 25,804,097 | CHH | 0.7780 | 7.63E-06 | 0.509 | 4 | 35,874,624 | CHH | 0.6996 | 8.37E-06 | 0.457 |
| 5 | 62,589,248 | CHH | 0.7169 | 8.32E-06 | 0.756 | 4 | 39,038,509 | CHH | 0.7064 | 6.25E-06 | 0.283 |
| 5 | 74,598,755 | CHH | 0.8863 | 6.84E-12 | 0.656 | 4 | 52,271,883 | CHH | 0.6594 | 3.71E-06 | 0.622 |
| 5 | 85,095,650 | CHH | 0.7184 | 2.62E-07 | 0.686 | 4 | 84,964,311 | CHH | 0.6728 | 2.70E-06 | 0.715 |
| 6 | 8,038,025 | CHH | 0.6797 | 7.12E-06 | 0.894 | 4 | 84,964,398 | CHH | 0.7087 | 6.39E-07 | 0.491 |
| 6 | 8,079,812 | CHH | 0.6777 | 5.62E-06 | 0.652 | 5 | 9,723,579 | CHH | 0.7514 | 4.67E-07 | 0.252 |
| 8 | 1,535,820 | CHH | 0.7355 | 6.45E-08 | 0.769 | 5 | 52,108,873 | CHH | 0.6650 | 7.12E-06 | 0.688 |
| 9 | 9,473,094 | CHH | 0.7064 | 5.03E-07 | 0.129 | 5 | 66,146,639 | CHH | 0.6630 | 5.75E-06 | 0.587 |
| 9 | 42,363,176 | CHH | 0.7152 | 8.94E-06 | 0.189 | 5 | 82,184,125 | CHH | 0.7091 | 3.86E-06 | 0.585 |
| 10 | 31,672,089 | CHH | 0.7185 | 8.05E-07 | 0.699 | 6 | 15,228,740 | CHH | 0.6827 | 1.22E-06 | 0.551 |
| 10 | 51,218,260 | CHH | 0.6867 | 1.39E-06 | 0.335 | 6 | 51,909,478 | CHH | 0.6530 | 6.62E-06 | 0.653 |
| 11 | 7,411,249 | CHH | 0.7501 | 3.85E-08 | 0.593 | 7 | 34,253,380 | CHH | 0.6460 | 6.72E-06 | 0.586 |
| 11 | 17,852,352 | CHH | 0.7665 | 5.04E-08 | 0.881 | 7 | 41,827,782 | CHH | 0.7100 | 5.34E-06 | 0.258 |
| 11 | 24,048,552 | CHH | 0.7478 | 2.44E-07 | 1.285 | 9 | 9,510,110 | CHH | 0.6849 | 1.09E-06 | 0.611 |
| 11 | 27,782,572 | CHH | 0.8720 | 9.02E-12 | 0.584 | 9 | 18,098,761 | CHH | 0.7158 | 2.08E-07 | 0.696 |
| Chromo-<br>some | position | context | Spearman<br>Rho | P-value | $h^2$ | Chromo-<br>some | position | context | Spearman<br>Rho | P-value | $h^2$ |
| 11 | 50,940,418 | CHH | 0.6782 | 3.98E-06 | 0.483 | 9 | 35,494,182 | CHH | 0.7683 | 4.51E-07 | 0.298 |
| 12 | 1,577,478 | CHH | 0.7332 | 7.41E-08 | 0.649 | 12 | 760,192 | CHH | 0.7357 | 3.18E-07 | 0.479 |
| 12 | 1,577,865 | CHH | 0.7140 | 1.01E-06 | 0.587 | 12 | 1,577,478 | CHH | 0.7185 | 2.60E-07 | 0.684 |
| 12 | 5,169,043 | CHH | 0.7061 | 3.58E-07 | 0.257 | 12 | 7,346,751 | CHH | 0.7028 | 5.11E-06 | 0.623 |
| 12 | 22,329,759 | CHH | 0.8497 | 4.00E-08 | 0.709 | 12 | 39,401,029 | CHH | 0.6368 | 9.98E-06 | 0.569 |

